# A CRISPRi-based genetic resource to study essential *Staphylococcus aureus* genes

**DOI:** 10.1101/2022.10.31.514627

**Authors:** Patricia Reed, Moritz Sorg, Dominik Alwardt, Lúcia Serra, Helena Veiga, Simon Schäper, Mariana G. Pinho

## Abstract

We have optimized a CRISPR interference system to facilitate gene knockdown in the gram-positive bacterial pathogen *Staphylococcus aureus.* Our approach used a CRISPRi system derived from *Streptococcus pyogenes,* which involves the co-expression of the *dcas9* gene encoding a catalytically inactive Cas9 protein and a customizable single guide RNA (sgRNA). In our system, *dcas9* is expressed from a single copy in the chromosome of methicillin resistant *S. aureus* (MRSA) strains COL or JE2, under the control of a tightly regulated promoter, inducible by anhydrotetracycline. The sgRNAs are expressed from a replicative plasmid under the control of a constitutively active promoter. This system enables efficient, inducible, knockdown of both essential and non-essential genes. Using this approach, we constructed the Lisbon CRISPRi Mutant Library (LCML) comprising 261 strains, in the JE2 background, containing sgRNAs targeting 200 essential genes/operons. This library facilitates the study of the function of essential *S. aureus* genes and is complementary to the Nebraska Transposon Mutant Library which consists of nearly 2000 strains, each carrying a transposon insertion within a non-essential gene. The availability of these two libraries will facilitate the study of *S. aureus* pathogenesis and biology.

**Abstract Importance:** *Staphylococcus aureus* is an important clinical pathogen that causes a high number of antibiotic resistant infections. The study of *S. aureus* biology, and particularly of the function of essential proteins, is of particular importance to develop new approaches to combat this pathogen. We have optimized a CRISPRi system that allows efficient targeting of essential *S. aureus* genes. Furthermore, we have used that system to construct a library of 261 strains which allow the depletion of essential proteins encoded in 200 genes/operons. This library, which we have named Lisbon CRISPRi Mutant Library (LCML), should facilitate the study of *S. aureus* pathogenesis and biology.

## Introduction

*Staphylococcus aureus* is a gram-positive bacterium that frequently colonizes the skin and nares of both humans and animals. It is also an opportunistic pathogen in community and hospital settings, causing a range of clinical conditions such as skin and soft tissue infections, bacteraemia or endocarditis (1). The emergence of multi-drug resistant strains, particularly methicillin-resistant *S. aureus* (MRSA), has complicated the treatment of *S. aureus* infections and MRSA strains are currently the second most common cause of global deaths associated with bacterial antimicrobial resistance (2). The discovery of novel antibiotics with unique modes of action against *S. aureus* is crucial in combating multidrug-resistant strains. One approach to address this challenge is the study of essential genes that encode core proteins required for bacterial survival, as these genes may encode potential targets for new antimicrobial drugs.

Despite their importance, the function of many essential genes in *S. aureus* remains poorly understood. Gene inactivation or repression are common approaches to elucidate the molecular function of genes in bacteria. Genetic tools to disrupt *S. aureus* genes or impair their transcription have become increasingly available (3). However, classical approaches such as gene deletion or transposon mutagenesis cannot be used to study essential genes, as they play indispensable roles in bacterial survival. Several alternative strategies have been developed including the exchange of the endogenous promoter of a gene by an exogenous inducible promoter via allelic exchange, or the use of antisense RNA technology (4, 5). However, the former approach is labour-intensive and unsuitable for large scale studies, while the latter has limitations due to variable efficacy.

In recent years, new tools based on CRISPR (clustered regularly interspaced palindromic repeats) interference systems have emerged as powerful tools to control gene expression. The most widely used CRISPRi system is derived from *Streptococcus pyogenes* and involves the co-expression of a gene encoding a catalytically inactive or dead (d)Cas9_Spy_ protein, which lacks endonuclease activity, and a customizable single guide RNA (sgRNA) (6, 7). The dCas9-sgRNA complex binds to the DNA target that is complementary to the sgRNA. Instead of introducing double strand breaks, this complex causes a steric block that halts transcription by RNA polymerase, leading to repression of the target gene or operon. Besides the complementary nucleotide sequence, the only other prerequisite for the complex to bind DNA is a three nucleotide long (NGG) recognition motif downstream of the complementary region of the targeted DNA strand, known as the protospacer adjacent motif (PAM).

CRISPRi systems based on *S. pyogenes* Cas9 (a type II CRISPR) have been established in several model organisms, including *Escherichia coli* (8), *Bacillus subtilis* (9), *Streptococcus pneumoniae* (10) or *S. aureus* (11–13). Type II CRISPR systems have been found in other bacteria besides *S. pyogenes*, including in *S. aureus.* Similar to CRISPR-Cas9*_Spy_*, these systems can be reprogrammed to serve as molecular tools for genome editing in eukaryotic cells (14, 15). The Cas9 from *S. aureus* (Cas9*_Sa_*) has raised some interest due to its smaller size compared to *S. pyogenes* Cas9*_Spy_* (1053 vs 1368 amino acids), hence allowing the use of adeno-associated virus with restrictive cargo size as vehicles for delivering Cas9 to animal cells (15). Additionally, Cas9*_Sa_* requires a longer PAM sequence (NNGRRT) (16), which increases specificity of DNA targeting and therefore reduces the potential for off-target effects, a concern when manipulating large genomes.

Previous CRISPRi systems for genetic manipulation of *S. aureus* use plasmids to encode the sgRNA and the dCas9, enabling the system to be easily moved into various strains (11–13). However, the use of multicopy plasmids for *dCas9* expression, combined with the fact that most inducible promoters used in *S. aureus* cannot be fully repressed, leads to systems in which dCas9 production cannot be completely shut down. This becomes particularly problematic when targeting essential genes, as basal levels of dCas9 can result in impaired growth or cell death. Here we report the construction of two variations of a gene knockdown system in *S. aureus,* based on the CRISPRi system originally established in *E. coli* by Qi and colleagues (8). The first system consists of two shuttle vectors, one encoding the dCas9 of either *S. pyogenes* or *S. aureus* and the other encoding the corresponding sgRNA with the gene-specific target sequence. The second system allows for efficient regulation of dCas9*_Spy_* production, as the gene encoding this protein was integrated into the genome under the control of a tightly controlled inducible promoter. This system was subsequently used to generate a knockdown library of 261 genes from 200 reported essential genes/operons in *S. aureus*, which we named the Lisbon CRISPRi Mutant Library (LCML). The LCML serves as a complementary resource to the widely used Nebraska Transposon Mutant Library (NTML), which includes mutants in virtually all non-essential genes in *S. aureus* (17). The use of both libraries will facilitate comprehensive functional studies of staphylococcal genes/operons.

## Results

### Construction of two-plasmid systems for CRISPR interference in *Staphylococcus aureus*

To establish a CRISPRi system in *S. aureus*, we chose two *E. coli-S. aureus* shuttle vectors to enable initial propagation and genetic manipulation in *E. coli*, followed by introduction into *S. aureus* by electroporation. *S. aureus* is not naturally competent, and transformation efficiency is very low compared to other model organisms (18, 19). The genes encoding a catalytically dead Cas9 from either *S. pyogenes* (dCas9*_Spy_*) or *S. aureus* (dCas9*_Sa_*), fused or not with superfast green fluorescent protein (sGFP), were cloned downstream of the cadmium-inducible promoter of the pCNX vector (20), which harbors a kanamycin resistance cassette for selection in *S. aureus*, generating the plasmids pBCB40 (*dcas9_Spy_*), pBCB41 (*dcas9_Spy_-sgfp*) and pBCB42 (*dcas9_Sa_-sgfp*). The tailor-made single guide RNAs from *S. pyogenes* (sgRNA*_Spy_*) or *S. aureus* (sgRNA*_Sa_*) were cloned downstream of a constitutively active promoter in the pGC2 vector (21), which contains a chloramphenicol resistance cassette for selection in *S. aureus* (Figure 1). Both sgRNA*_Spy_* and sgRNA*_Sa_* consist of two regions: the 5’ end contains the variable region for targeted DNA binding (first 20 nucleotides of sgRNA*_Spy_* or first 22 nucleotides of sgRNA*_Sa_*), while the downstream region contains a constant region for dCas9 binding (different for dCas9*_Spy_* and dCas9*_Sa_*) and transcription termination. Targeting of specific genes of interest can be accomplished by replacing the 20 (or 22) nucleotides of the variable region, using inverse PCR, for the specific sequence of interest. In the presence of dCas9, the sgRNA forms a complex with the inactive nuclease and guides it to the complementary DNA, where the complex acts as a physical blockage for RNA polymerase, impairing transcription. A key requirement for an efficient CRISPRi system is the ability to shut down dCas9 production, so that the system can be turned ON/OFF. Western blot analysis of dCas9_Spy_ levels in strain BCBMS02 (expressing cadmium-inducible *dcas9_Spy_-sgfp*) showed the presence of dCas9*_Spy_*-sGFP even in samples grown in the absence of inducer, indicating that the cadmium-inducible promoter is leaky (Figure S1A). Despite this limitation, we proceeded to evaluate the efficiency of the system. For that, we constructed the reporter strains BCBMS05 (expressing *dcas9_Spy_-sgfp*) and BCBLS01 (expressing *dcas9_Sa_-sgfp*), in the background of a derivative of strain NCTC8325-4, where the non-essential gene *rodZ,* which encodes the septal protein RodZ, was fused to the gene encoding red fluorescent protein eqFP650. Both strains allow easy evaluation of eqFP650-RodZ (red fluorescence) depletion and dCas9-sGFP (green fluorescence) production by fluorescence microscopy. We then designed four different dCas9*_Spy_* sgRNAs, targeting the coding or the template strand of *fp650-rodZ,* as well as its upstream region (Figure 2A). We also designed three dCas9*_Sa_* sgRNAs, two targeting the coding strand and one targeting the template strand of *fp650-rodZ* (Figure 2A). To analyze the efficiency of the two systems, we used pairs of strains, both expressing the same sgRNA targeting *fp650-rodZ* plus either *dcas9-sgfp* expressed from a pCNX-based plasmid (test) or the empty pCNX vector (control). The two strains were grown in the presence of the inducer cadmium, and then mixed and placed on the same microscopy slide for direct comparison (Figure 2B). With dCas9*_Spy_* system, we observed that sgRNAs targeting the promoter sequence on either strand (sg-*fp650rodZ-Spy*1 and sg-*fp650rodZ-Spy*2) or the gene sequence on the coding strand (sg-*fp650rodZ-Spy*3) were efficient in depleting eqFP650-RodZ, leading to a strong reduction in red fluorescence intensity when compared to the corresponding control strains lacking dCas9*_Spy_*-sGFP (Figure 2C, E). On the contrary, the sgRNA that targeted the template strand of *fp650-rodZ* gene (sg-*fp650rodZ-Spy*4) was not effective (Figure 2C, E), in agreement with previous data for *E. coli* describing that targeting the template strand is not efficient (8). Notably, the eqFP650 signal in this strain was delocalized from the septum to the cytosol and it even increased 2.5-fold compared to control cells (expressing sg-*fp650rodZ-Spy*4 but lacking dCas9*_Spy_*-sGFP). The sgRNA of this strain targets *rodZ,* downstream of *fp650*, so it is possible that transcription and translation of *fp650* can still occur, producing a soluble, truncated eqFP650.

**Figure 1.**
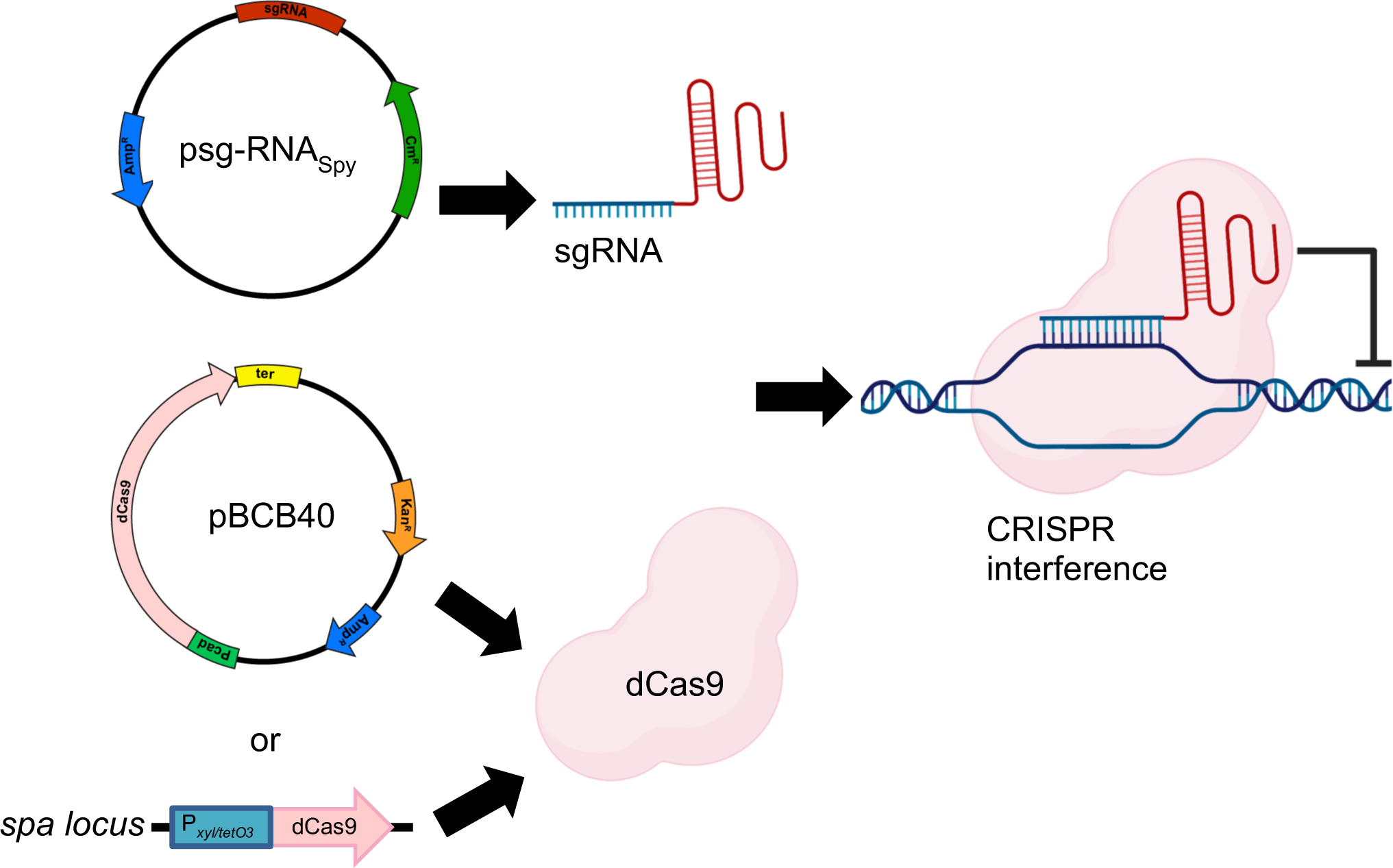
Plasmid-based CRISPRi system. The sgRNAs for *S. pyogenes* Cas9 are constitutively expressed from the psg-RNA*_Spy_* plasmid. dCas9 can either be expressed from a second plasmid, under the control of a cadmium-inducible promoter (pBCB40), or from the *spa* locus in the *S. aureus* chromosome. Transcription inhibition occurs when a complex of dCas9 and sgRNA binds specific target genes, thus blocking the RNA polymerase.

**Figure 2.**
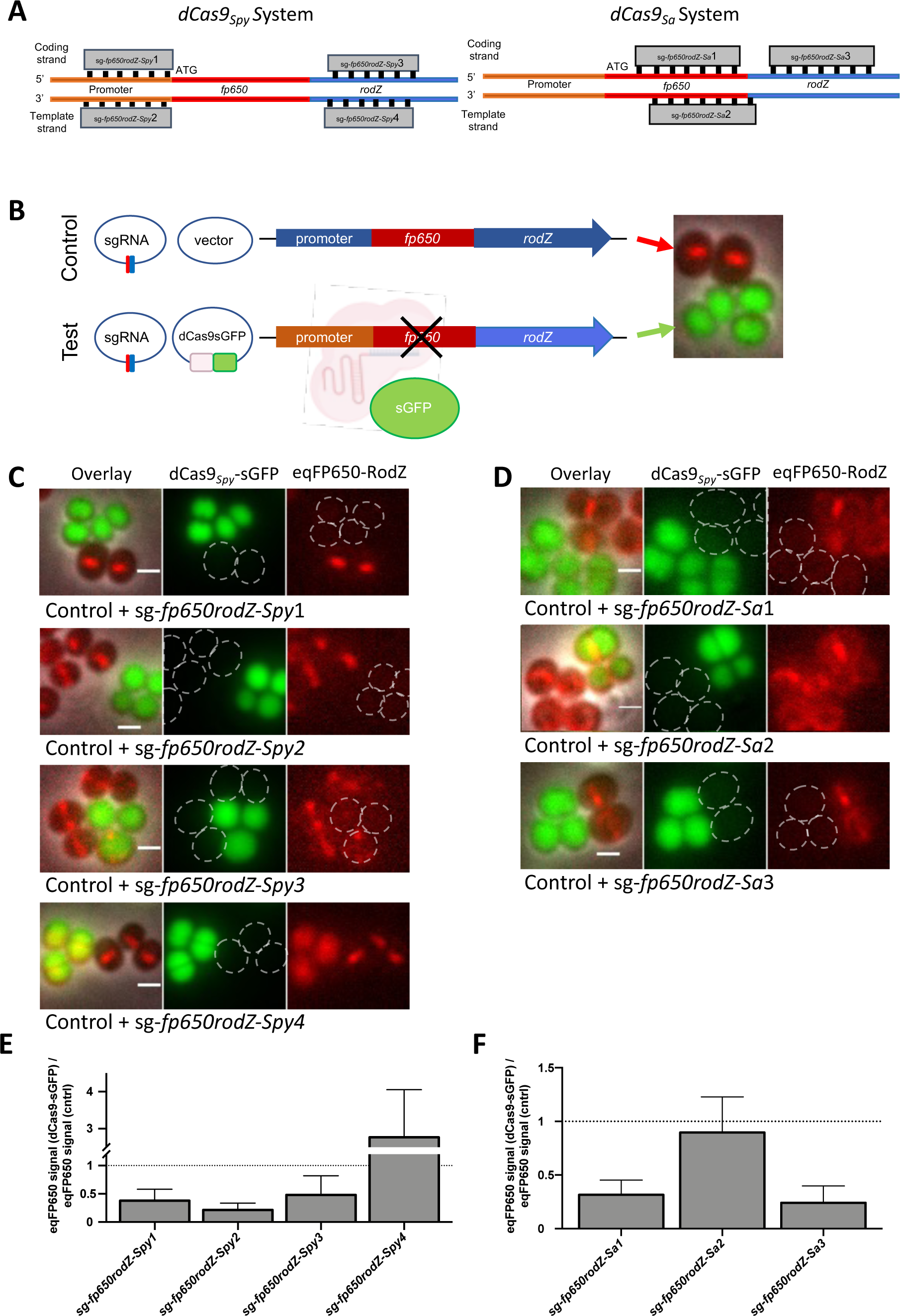
Plasmid-based CRISPRi systems using either dCas9*_Spy_* or dCas9*_Sa_* inhibit gene expression in *S. aureus*. (**A**) Schematic overview of sgRNA target sites in *fp650-rodZ* used to test the dCas9*_Spy_* (left) or dCas9*_Sa_* (right) CRISPRi systems. **(B)** Experimental setup to test CRISPRi systems. Genetically modified NCTC8324-5 cells expressing an eqFP650 fusion to the non-essential septal protein RodZ were transformed with plasmids encoding sgRNA against *fp650-rodZ* and either an empty vector control (top, control cells) or the same vector encoding dCas9-sGFP (bottom, test cells). Strains were grown separately in the presence of 0.1 µM CdCl_2_ and subsequently mixed to perform the microscope experiments. Control cells lack dCas9 and therefore do not impair production of eqFP650-RodZ, which can be seen as a septal band (red). Test cells express a cytoplasmic sGFP fusion of dCas9 (green) which is targeted by sgRNA to specific regions of *fp650-rodZ,* blocking its expression. **(C, D)** Fluorescence microscopy images of mixtures of control cells and test cells (encoding dCas9-sGFP) expressing sgRNAs shown in panel B using either the dCas9*_Spy_* (C) or dCas9*_Sa_* (D) systems. From left to right, overlay image, green channel image showing dCas9-sGFP signal and red channel image showing eqFP650-RodZ signal. Scale bars, 1µm. (**E, F**) Bar charts showing the ratio of the median eqFP650 signal of cells expressing dCas9-sGFP versus the median eqFP650 signal of the corresponding control cells, both expressing the same sgRNA. Error Bars represent SEM calculated from three independent experiments.

We noticed that depletion of eqFP650-RodZ, resulting in loss of red fluorescence signal, occured even in the absence of *dcas9* induction (Figure S1B). This indicates that the leaky *dcas9* expression results in the production of low levels of dCas9 protein, as seen by western blot (Figure S1A), that are sufficient to efficiently drive the CRISPRi system in the absence of inducer, i.e., that the system cannot be efficiently turned off.

Next, we tested the alternative version of the CRISPRi system that uses an inactive variant of a Cas9*_Sa_* native to *S. aureus*. Similarly to the dCas9*_Spy_* system, this system was also effective when the sgRNA*_Sa_* targeted the coding strand (sg-*fp650rodZ*-*Sa*1 and sg-*fp650rodZ*-*Sa*3) within the *fp650-rodZ* gene, as seen by the decreased red fluorescence of these strains compared to the corresponding control strains lacking dCas9*_Sa_* (Figure 2D, F). No inhibition of *fp650-rodZ* expression was observed when the sgRNA targeted the template strand (sg-*fp650rodZ*-*Sa*2). We achieved similar knockdown efficiencies with both the dCas9*_Spy_* and the dCas9*_Sa_* system, in both cases leading to a decrease in the relative fluorescence signal of eqFP650-RodZ (Figure 2E and F). Therefore, we decided to proceed with *S. pyogenes* dCas9 as it is less limited in target selection due to the shorter PAM sequence, a relevant criterion for work in the low GC content bacterium *S. aureus*.

### CRISPRi system with chromosome-encoded *dCas9_Spy_* enables repression of essential genes transcription in *S. aureus*

To construct an efficient knockdown system, that can be used to target essential genes in *S. aureus*, we re-designed the CRISPRi system described above, to minimize production of dCas9 in the absence of inducer. For that, we integrated *dcas9_Spy_* into the *spa* locus of the staphylococcal genome, to reduce the gene copy-number to one. This chromosomal copy of *dcas9_Spy_* was placed under the control of a tetracycline-inducible *xyl/tetO* promoter (22). Furthermore, we used a weak ribosome binding site (RBS) (23) separated from the start codon by only 3 nucleotides to decrease translation efficiency (Figure 3A and S2). This P*_xyl/tetO3_*-*dcas9_Spy_* construct was integrated into the *spa* locus of two MRSA strains, JE2 (generating strain BCBMS14) and COL (generating strain BCBMS15). In both backgrounds dCas9 was detected by western blot analysis in cells grown in the presence of the inducer anhydrotetracycline (aTc), but not in its absence, indicating that *dcas9_Spy_* expression was now under tight control (Figure 3B). Cells expressing *dcas9_Spy_* from the chromosome in the presence of aTc were only mildly impaired in growth (Figure 3C).

**Figure 3.**
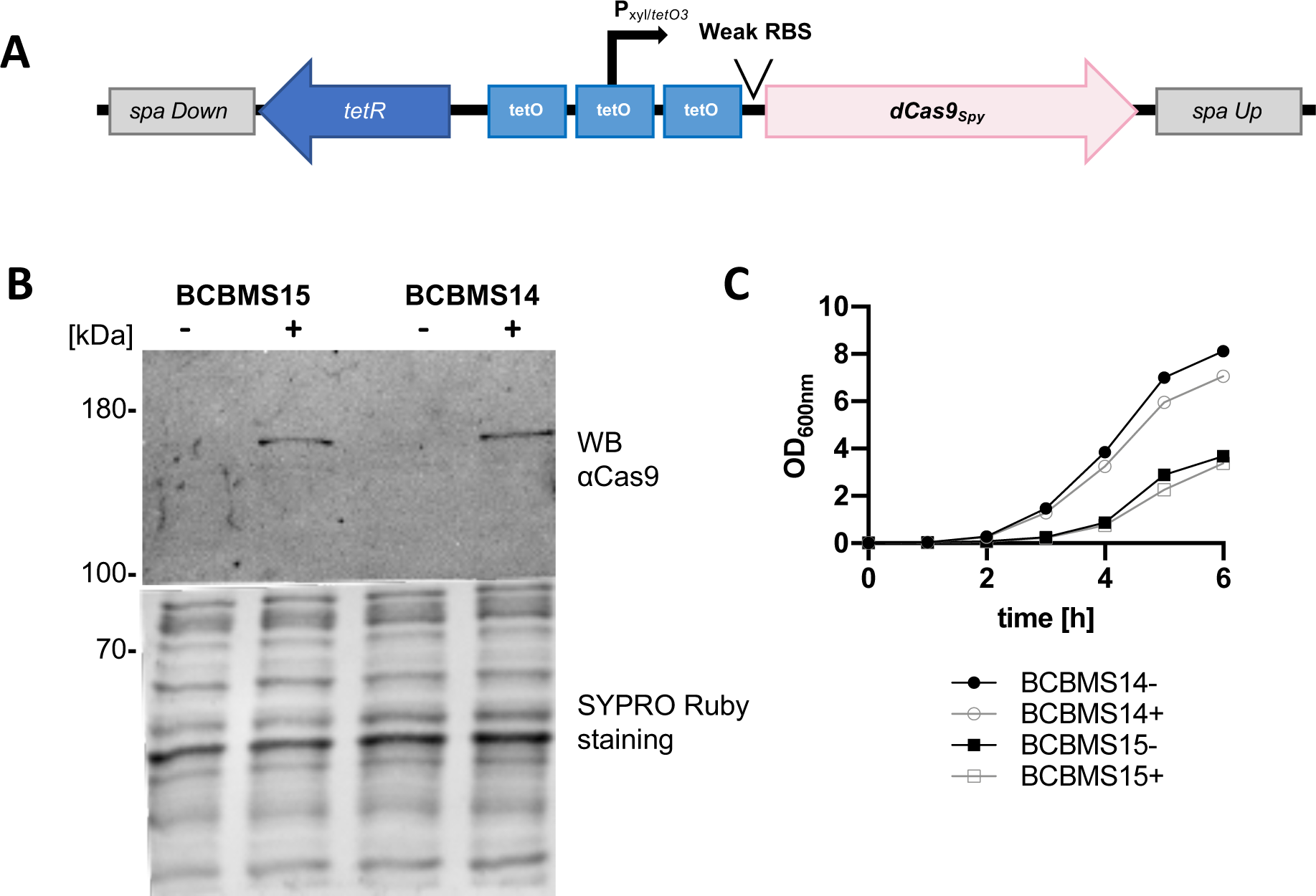
CRISPRi system with chromosome-encoded *dCas9_Spy_* allows tight regulation of dCas9*_Spy_* production. (**A**) Schematic representation of the genomic construct inserted into the *spa* locus of strains BCBMS14 (JE2 background) and BCBMS15 (COL background), which allows tight control of dCas9*_Spy_* production due to the presence of three TetR-binding sites (*tetO*) flanking and overlapping the xylose promoter and of an inefficient ribosome binding site (RBS). The sgRNA is expressed from plasmid psg-RNA_Spy_ (not shown) (**B**) Western blot using anti-Cas9 antibody, of protein extracts from induced (+) and non-induced (-) BCBMS15 and BCBMS14 cultures showing tight regulation of dCas9*_Spy_* expression. SYPRO Ruby staining was used as a loading control. **(C)** Growth curves of induced (+, 100 ng/ml aTc) and uninduced (-) BCBMS14 and BCBMS15.

We then tested the efficiency of the improved CRISPRi system when targeting essential genes. For that we designed sgRNAs targeting the essential cell division genes *ftsZ, murJ* and *pbpA,* encoding the cell division protein FtsZ, the peptidoglycan precursor lipid II flippase MurJ and the penicillin-binding protein PBP1, respectively. The resulting CRISPRi strains were analyzed by growth assays and microscopy (see Figure 4 and Figure S3 for strains in the background of JE2 and COL, respectively). Strains showed severe growth inhibition upon induction of *dcas9_Spy_* expression (but not in its absence) and the expected phenotype for the depletion of each gene was observed: while depletion of MurJ did not have a visible phenotype, depletion of FtsZ led to greatly enlarged, spherical cells without detectable septa (24) and depletion of PBP1 led to enlarged, elongated cells (25). Importantly, these phenotypes were barely observed in the absence of inducer, confirming that *dcas9* is under tight regulation in this improved CRISPRi system.

**Figure 4.**
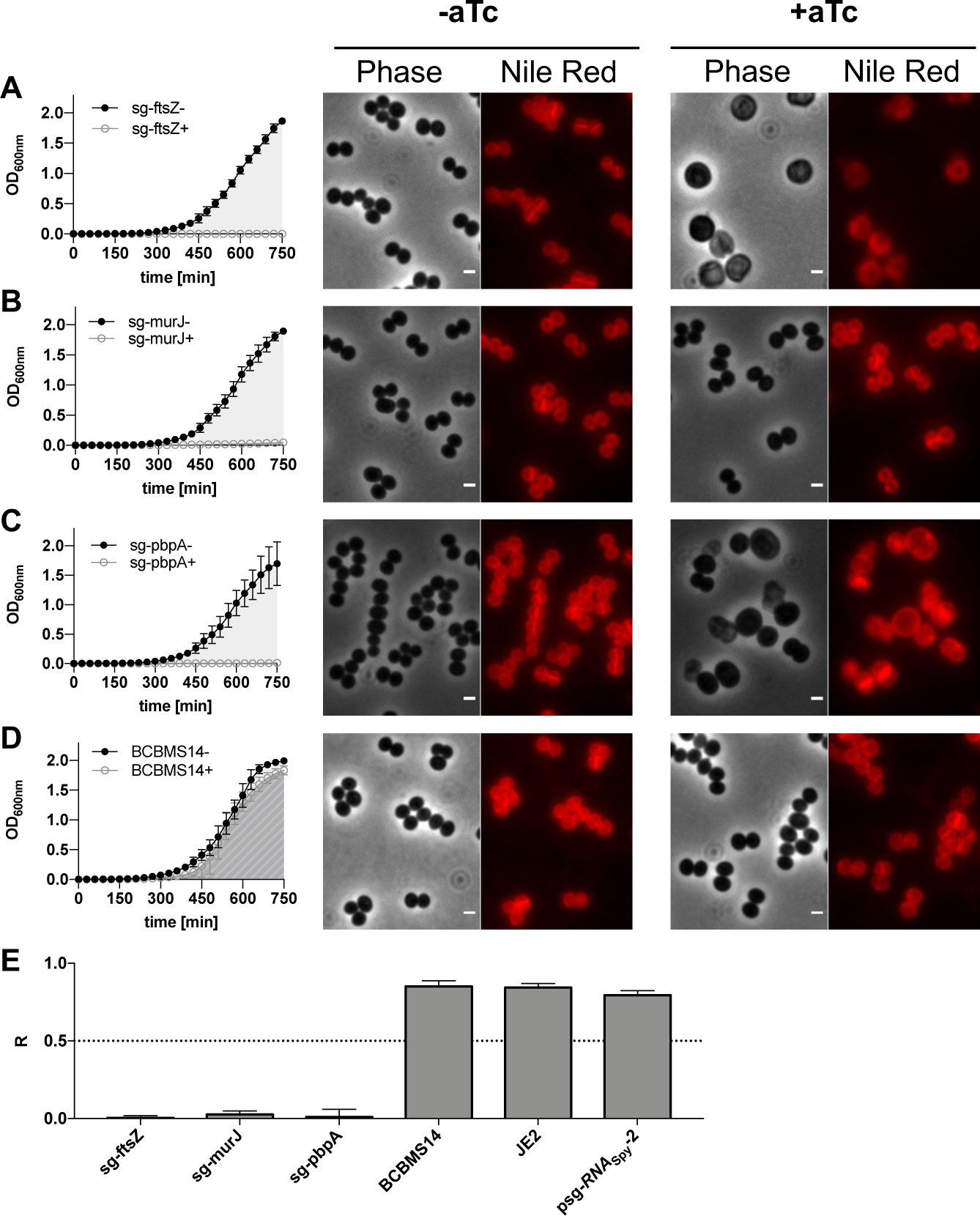
CRISPRi system with chromosome-encoded *dCas9_Spy_* is suitable to target essential genes in *S. aureus*. **(A-D)** Growth assays performed in 96-well plates at 30 °C in TSB, in the absence (-) or the presence (+) of aTc (200 ng/ml, inducer for *dCas9* expression) of BCBMS14 strains expressing sgRNAs targeting the essential genes *ftsZ* (BCBMS16, A), and *murJ* (BCBMS18, B) and *pbpA* (BCBMS17, C). The strain BCBMS14 (D) was used as a control. Phase contrast and fluorescence microscopy images of cells stained with Nile Red (membrane stain) are shown for each strain. Scale bars, 1µm. (**E**) Graphs show the ratio (R) of the area under the curve (AUC) of growth curves obtained in the presence (+) versus the absence (-) of aTc for strains shown in A-D, as well as for parental strain JE2 and for BCBMS14 containing psg-*RNA*_Spy_ lacking a specific sgRNA sequence. R can vary between 0 (complete growth inhibition in the presence of the inducer) and 1 (no growth inhibition). Error bars indicate SEM calculated with data from three independent experiments.

### Construction of a CRISPRi library of conditional mutants of essential *S. aureus* genes

The Nebraska Transposon Mutant library is an extremely useful resource for the study of *S. aureus,* as it contains transposon mutants in virtually all non-essential genes of MRSA strain JE2 (17). A complementary library containing depletion mutants for the remaining, essential, genes would be a valuable tool and therefore we decided to use the improved CRISPRi system for this purpose. Given that CRISPRi affects the expression of entire operons and not just of the target gene, we are unable to construct single mutants for each essential gene using this technology. We therefore targeted each essential operon, which allows users to perform initial screenings that may have to be followed up by detailed studies of individual genes in operons. We identified 200 predicted monocistronic or polycistronic operons (Table S1) containing genes originally described in the literature as essential using antisense RNA (26) or transposon library screenings (27, 28). Identity of genes belonging to each operon was based on the information available in aureowiki (https://aureowiki.med.uni-greifswald.de/Main_Page). We designed sgRNAs targeting the first essential gene of each essential operon. sgRNAs targeting more than one gene in an operon were designed for 46 operons resulting in a total of 261 sgRNAs tested. sgRNAs were cloned into psg-*RNA_Spy_* using inverse PCR with primer 5846-EcR and a primer containing the specific sgRNA sequence to be introduced (Table S2). The resulting library of plasmids encoding the sgRNAs was electroporated into BCBMS14, generating a library of 261 conditional mutants strains, targeting 200 operons, listed in Table S3, which we named the Lisbon CRISPRi Mutant Library (LCML). Growth of these strains in the presence or absence of the aTc inducer (200 ng/ml) was followed in a 96-well plate reader, for 750 min, at 30 °C. The ratio (R) of the area under the curve (AUC) of the growth curve obtained in the presence versus the absence of inducer was used to evaluate the efficiency of growth inhibition (Figure S4). 148 of the 261 strains showed a clear growth reduction upon induction (R ≤ 0.5, an arbitrary threshold chosen to decide if further optimization was required). However, 113 of the strains had milder growth deficits (R >0.5). For these, we designed a second sgRNA (sgRNA_B) targeting an alternative region of the gene. Of the 113 strains for which a second sgRNA was designed, 48 showed R lower than 0.5 (Figure S4 shows R data for each strain when using the sgRNA, A or B, with highest efficiency). For 65 strains for which R was higher than 0.5, growth analysis was repeated at 37 °C, given that some mutants may show a stronger growth defect at 37 °C than at 30 °C. Figure S5 shows that inhibition of the expression of a further 20 genes led to impaired growth at 37 °C. Overall we achieved a success rate of ∼83% as we observed strong growth inhibition (R ≤ 0.5) for 216 of the 261 strains constructed (Figure 5A). The LCML clones showing the greatest growth impairment are listed in Supplementary Table 4 with the sgRNA sequence and R values listed.

**Figure 5.**
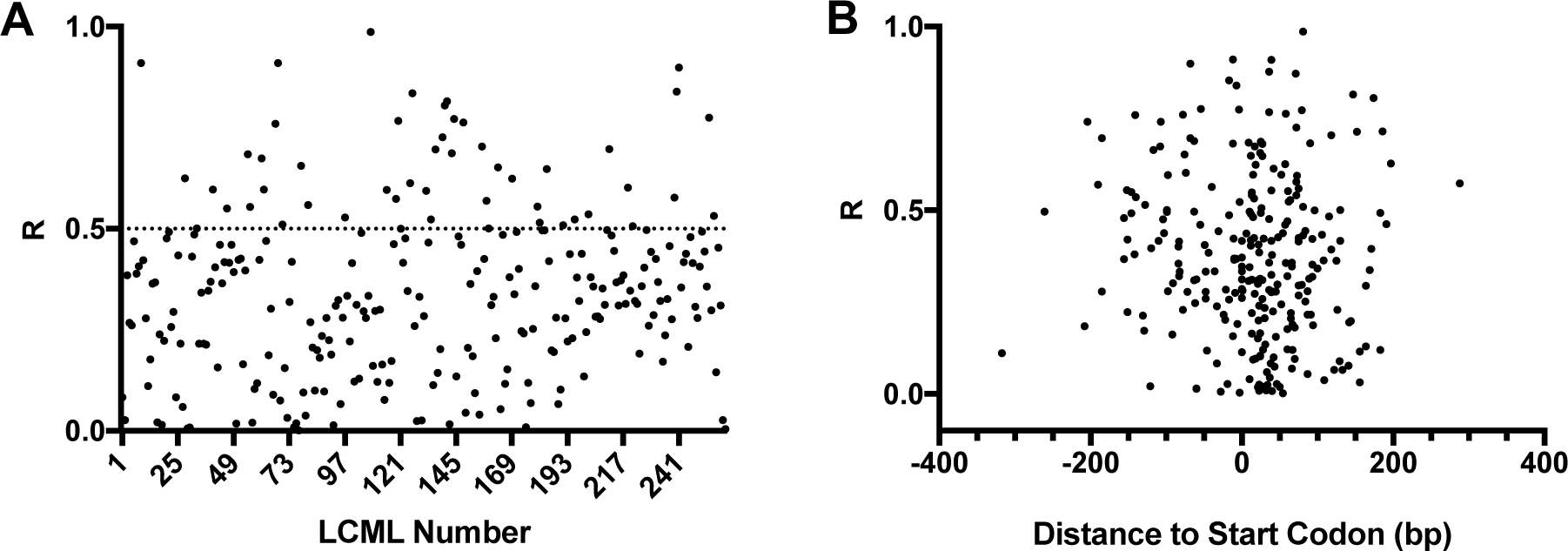
Efficiency of growth inhibition by CRISPRi system with chromosome-encoded *dCas9_Spy_*. **(A).** The ratio (R) of the area under the curve (AUC) of growth rates obtained in the presence versus the absence of aTc (inducer for *dCas9* expression) was calculated for LCML strains expressing 261 different sgRNAs. Graph shows that R is lower than 0.5 for ∼80% of the strains. **(B)** R was plotted against distance between the sgRNA target position and the start-codon of the targeted gene, showing that there is no correlation between this distance and the efficiency of growth inhibition by the CRISPRi system.

To determine if there were target regions for sgRNA that resulted in higher efficiency of the CRISPRi system, we plotted R for each sgRNA against their target position relative to the start-codon of the corresponding gene (Figure 5B) and tested those targeting the coding strand for statistically relevant correlation using a Spearman correlation test. For that we split the data into two sets: the first set included the 87 sgRNAs that bind upstream of the start codon and the second set included the 174 sgRNAs targeting in the gene. sgRNAs targeting the template strand were excluded from the analyses due to small sample size (n=8). We found no statistically significant correlation between efficiency of growth inhibition and localization of sgRNAs targeted sequence relative to start codon, for either the first set (r: -0.06365; p-value: 0.5581) or the second set (r: 0.09877; p-value: 0.1947).

## Discussion

Construction of *S. aureus* conditional knockouts or gene deletion mutants using traditional approaches that require recombination events is labour-intensive and time-consuming. In contrast, use of a CRISPRi system allows the generation of conditional mutants through the simple exchange of the target sequence of the sgRNA, so that it specifically inactivates any target gene/operon. In this study, we developed different versions of CRISPRi systems for *S. aureus*, using dCas9 proteins originated from both *S. pyogenes* and *S. aureus*. In our first approach, sgRNAs and *dcas9* from either *S. pyogenes* or *S. aureus* were expressed from two replicative vectors, allowing rapid introduction into different *S. aureus* strains and mutants. Staphylococcal Cas9*_Sa_* is significantly smaller than Cas9 from *S. pyogenes* (1053 vs 1368 amino acid residues) and it requires a longer PAM sequence. The smaller size of the nuclease is important when used in eukaryotes because of limited packaging size of adenovirus systems and lower transfection efficiency of very large plasmids. Additionally, a longer PAM sequence should decrease off-target effects, which is a problem when manipulating eukaryotic cells due to their large genome size. Both advantages are less important in bacteria. Nevertheless, we hypothesized that the expression of the smaller, native nuclease, could be advantageous in *S. aureus*. However, the two CRISPRi systems, with either *S. pyogenes* or *S. aureus* dCas9, worked similarly. Given that *S. pyogenes* dCas9 is less limited in target selection due to the shorter PAM sequence, we decided to proceed with dCas9*_Spy_* for further optimization of the system. Optimization was required because low levels of dCas9 were produced even in the absence of inducer when using these two-plasmid systems. To reduce gene transcription of *dcas9* in the absence of an inducer to a minimum we (i) integrated *dcas9* into the genome of *S. aureus* MRSA strains JE2 and COL, to reduce its copy number to one, (ii) exchanged the leaky cadmium-inducible promoter with a tighter *xyl/tetO* promoter, and (iii) decreased translation efficiency by switching to an inefficient ribosome binding site. The resulting strains, BCBMS14 (JE2 background) and BCBMS15 (COL background), produced dCas9 under tight control as it was not detected by western blot in the absence of aTc inducer, but reached sufficient levels to silence essential genes upon addition of the inducer. Targeting of essential genes *ftsZ*, *pbpA*, or *murJ* with this improved CRISPRi system led to a severe decrease of growth, showing the efficiency of the system.

Given the efficacy of the improved CRISPRi system to silence essential genes, we constructed a library of CRISPRi conditional mutants in the strain BCBMS14, each containing a specific sgRNA targeting one of 261 genes, corresponding to 200 operons which harbour at least one essential gene. Overall, we were able to construct strains to silence genes in 169 of these essential operons. It is possible that some of the tested genes for which expression inhibition did not result in severe growth defects (R>0.5) are not actually essential for bacterial survival under the growth conditions tested, which would justify the lack of efficiency of some sgRNAs. For example, it was reported that *recU* (R = 0.595) can be depleted in *S. aureus*, although mutants display chromosome segregation and DNA damage repair defects (29). Similarly, a *ltaS* mutant (R = 0.597) is viable in *S. aureus* but displays temperature sensitivity (30). Viable (although often exhibiting growth defects) mutants of other genes that displayed R higher than 0.5 when gene expression was inhibited with CRISPRi can also be found in the literature, namely *parE* (31), *dnaK* (32), *secDF* (33), *ackA* (34), *murA* (35), *ureA* (36), *xpt* (37), *mraZ* (38) and *gdpP* (39).

Large variation in sgRNA efficiency has been reported in several published studies (8, 40–42). In *E. coli*, the distance between transcription start site and sgRNA target was reported by Qi and colleagues to affect knockdown efficiency (8) but this effect was not observed in genome-wide screens (43, 44). We selected PAM sequences close to the start codon, so all the designed sgRNAs target regions between -400 to +400 nucleotides from the start codon. Given the large number of sgRNAs tested, we evaluated if there were specific locations that corresponded to more efficient sgRNAs. However, as shown in Figure 5B, we could not find a correlation between the distance of the sgRNA target to the start codon of the corresponding gene and sgRNA efficiency, and therefore could not establish guiding principles for sgRNA design.

The constructed Lisbon CRISPRi Mutant Library of conditional mutants in essential genes/operons of *S. aureus* is complementary to the widely used Nebraska Transposon Mutant library, which includes mutants in virtually all non-essential genes in *S. aureus* (17). The combined use of the two libraries will allow the evaluation of the function of any gene/operon of this important pathogen.

## Materials and Methods

### Bacterial strains and growth conditions

Plasmids and strains used in this study are described in Tables 1 and 2. *S. aureus* cells were grown at 37 °C with aeration in liquid tryptic soy broth (TSB; Difco) or on solid tryptic soy agar (TSA; VWR) medium, supplemented, when necessary, with antibiotics (chloramphenicol (Cm), Sigma-Aldrich, 10 µg/ml in liquid media, 20 µg/ml in solid medium; kanamycin (Kan), Apollo Scientific, 50 µg/ml; neomycin (Neo), Sigma-Aldrich, 50 µg/ml) and inducer (cadmium chloride, Sigma-Aldrich, 0.1 µM; or anhydrotetracycline (aTc), Sigma-Aldrich, 100-200 ng/ml). *E. coli* cells were grown at 37 °C with aeration in either LB (Luria-Bertani broth; VWR) or LA (Luria Agar; VWR), supplemented with ampicillin (Apollo Scientific, 100 µg/ml) when required. Strain growth in liquid media was followed by measurement of optical density at 600 nm (OD_600_).

**Table 1.** Plasmids used and constructed in this study.

| Name | Description | Reference |
| --- | --- | --- |
| pTRC99a-P7 | Vector containing *sgfp-p7*; Amp<sup>R</sup> | (46) |
| pFP60-F | pTnT with *fp650* in forward orientation | (56) |
| pgRNA<sub>bacteria</sub> | Vector containing sgRNA<sub>Spy</sub> | (8, 45) |
| pdCas9<sub>bacteria</sub> | Vector containing *dcas9<sub>Spy</sub>* | (8, 45) |
| pMAD | *E. coli* / *S. aureus* shuttle vector with a thermosensitive origin of replication for Gram positive bacteria; Amp<sup>R</sup>; Ery<sup>R</sup>; *lacZ* | (57) |
| pGC2 | Replicative *E. coli* / *S. aureus* shuttle vector; Amp<sup>R</sup>; Cm<sup>R</sup> | (21) |
| pBCB13 | pMAD with up- and downstream regions of *spa* and *Pspac-lacI* region from pDH88 | (48) |
| pCNX | Replicative *E. coli* / *S. aureus* shuttle vector containing the cadmium-inducible *Pcad* promoter; Amp<sup>R</sup>; Kan<sup>R</sup>/Neo<sup>R</sup> | (20) |
| pCNX<sub>sGFP</sub> | pCNX derivative containing *sgfp* from plasmid pTRC99a-P7 | This study |
| pBCB40 | pCNX derivative containing *S. pyogenes dcas9*; Amp<sup>R</sup>; Kan<sup>R</sup>/Neo<sup>R</sup> | This study |
| pBCB41 | pCNX derivative containing *S. pyogenes dcas9-sgfp*; Amp<sup>R</sup>; Kan<sup>R</sup>/Neo<sup>R</sup> | This study |
| pBCB42 | pCNX derivative containing *S. aureus dcas9-sgfp*; Amp<sup>R</sup>; Kan<sup>R</sup>/Neo<sup>R</sup> | This study |
| pBCB43 | pBCB13 derivative lacking *P<sub>spac</sub>* and *lacI*, containing *tetR*, *P<sub>xyltetO</sub>* and a downstream located *tetO* site, Amp<sup>R</sup>; Ery<sup>R</sup>; *lacZ* | This study |
| pBCB44 | pBCB43 derivative containing *dcas9<sub>Spy</sub>* under *P<sub>xyl/tetO3</sub>* control; Amp<sup>R</sup>; Ery<sup>R</sup>; *lacZ* | This study |
| psg-RNA<sub>Spy</sub> | pGC2 derivative containing the sgRNA cloned from pgRNA<sub>bacteria</sub> plasmid; Cm<sup>R</sup> | This study |
| psg-RNA<sub>Spy-2</sub> | psg-RNA<sub>Spy</sub> derivative lacking the 20 target specific base pairing nucleotides; Cm<sup>R</sup> | This study |
| psg-*fp650rodZ-Spy1* | psg-RNA<sub>Spy</sub> derivative with sgRNA altered to sg-*fp650rodZ-Spy1*; Amp<sup>R</sup>; Cm<sup>R</sup> | This study |
| psg-*fp650rodZ-Spy2* | psg-RNA<sub>Spy</sub> derivative with sgRNA altered to sg-*fp650rodZ-Spy2*; Amp<sup>R</sup>; Cm<sup>R</sup> | This study |
| psg-*fp650rodZ-Spy3* | psg-RNA<sub>Spy</sub> derivative with sgRNA altered to sg-*fp650rodZ-Spy3*; Amp<sup>R</sup>; Cm<sup>R</sup> | This study |
| psg-*fp650rodZ-Spy4* | psg-RNA<sub>Spy</sub> derivative with sgRNA altered to sg-*fp650rodZ-Spy4*; Amp<sup>R</sup>; Cm<sup>R</sup> | This study |
| psg-*fp650rodZ-Sa1* | pGC2 derivative containing the sg-*fp650rodZ-Sa1* under control of a constitutive active promoter; Amp<sup>R</sup>; Cm<sup>R</sup> | This study |
| psg-*fp650rodZ-Sa2* | pGC2 derivative containing the sg-*fp650rodZ-Sa2* under control of a constitutive active promoter; Amp<sup>R</sup>; Cm<sup>R</sup> | This study |
| psg-*fp650rodZ-Sa3* | pGC2 derivative containing the sg-*fp650rodZ-Sa3* under control of a constitutive active promoter; Amp<sup>R</sup>; Cm<sup>R</sup> | This study |
| psg-*ftsZ* | psg-RNA<sub>Spy</sub> derivative with sgRNA altered to sg-*ftsZ*; Amp<sup>R</sup>; Cm<sup>R</sup> | This study |
| psg-*pbpA* | psg-RNA<sub>Spy</sub> derivative with sgRNA altered to sg-*pbpA*; Amp<sup>R</sup>; Cm<sup>R</sup> | This study |
| psg-*murJ* | psg-RNA<sub>Spy</sub> derivative with sgRNA altered to sg-*murJ*; Amp<sup>R</sup>; Cm<sup>R</sup> | This study |
| pMAD<sub>fp650rodZ</sub> | pMAD containing *rodZ* upstream region-*eqfp650-rodZ-rodZ* downstream region; Amp<sup>R</sup> Ery<sup>R</sup> | This study |

### DNA purification and manipulation

Plasmid DNA was extracted from *E. coli* cells using Wizard SV Plus Miniprep Kit (Promega) or the QIAprep Spin Miniprep Kit (Qiagen), following the manufacturer’s instructions. DNA was digested with FastDigest restriction enzymes (ThermoFisher) by incubating 0.5-1 µg DNA at 37 °C for 1 hour, with 1x FastDigest buffer, 1 µl of the selected endonuclease and 1 µl of Fast Alkaline Phosphatase (ThermoFisher), when necessary. DNA fragments were purified using Wizard SV Plus Cleanup Kit (Promega), according to manufacturer’s instructions. DNA ligations were performed by standard molecular biology techniques using T4 DNA Ligase (ThermoFisher). PCR reactions were performed using either Phusion DNA Polymerase for cloning purposes or DreamTaq Polymerase for colony screenings (both ThermoFisher), following the manufacturer’s instructions. For inverse PCR, one primer was 5’ phosphorylated using a T4 Polynucleotide Kinase (ThermoFisher) using recommended reaction conditions. After PCR amplification, 1 µl DpnI (NEB) was added to the PCR mix, which was incubated for 1 h at 37 °C followed by purification. 5 µl of the purified PCR product were used for ligation using T4 DNA ligase followed by transformation.

### Plasmid and strain construction

Plasmids and primers used in this study are described in Tables 1 and 3, primers used to generate the different sgRNAs for the LCML are described in Table S2. To construct the pBCB40 plasmid, *dcas9_Spy_* was amplified from plasmid pdCas9_bacteria (8, 45) using primers 5450/5451. The PCR product and the vector pCNX (20) were digested with PstI and SalI and the fragment containing *dcas9_Spy_* was ligated downstream of the cadmium-inducible promoter from pCNX. To construct pBCB41, primers 3714/5450 were used to amplify *dcas9_Spy_* with a linker insertion and *sgfp* was amplified from the plasmid pTRC99a-P7 (46) using primers 3715/5452. The two fragments were joined by overlap PCR using primers 5450/5452, digested with PstI and SalI and ligated into pCNX, digested with the same enzymes.

To construct the pBCB42, we first constructed the pCNX_sGFP plasmid by amplifying the *sgfp* gene sequence from pTRC99a-P7 plasmid using the primer pair 5602/5603 and cloning it into pCNX using EcoRI and BamHI restriction sites. The *dcas9_Sa_* gene sequence was kindly provided by F. Ann Ran (15). The sequence, comprising a ribosome binding site and the *dcas9_Sa_* gene plus restriction sites SalI and BamHI, was synthesized (NZYTech), digested with the respective restriction enzymes and cloned into the pCNX_sGFP plasmid, resulting in plasmid pBCB42.

To construct plasmid pBCB43 a fragment of the antisense *secY* expression cassette in pIMAY (47) was amplified using primers 7015/7016. The PCR product, comprising *tetR*, *P_xyl-tetO_* and a downstream located *tetO* site, was digested with EcoRI/NheI and ligated into similarly digested pBCB13 (48) lacking *lacI* and P*_spac_*. The resulting plasmid, pBCB43, was SmaI/EagI digested to clone *dcas9_Spy_*, amplified from pBCB40 with the primer pair 7268/7272, downstream of *tetO* of pBCB43, via Gibson assembly (49), generating plasmid pBCB44, verified by sequencing.

Plasmid psg-*RNA_Spy_* was constructed by cloning the sgRNA sequence including the minimal constitutive promoter and transcription terminator from plasmid pgRNA_bacteria (8, 45) into pGC2 (21). For that, the multiple cloning site of pGC2 was altered via inverse PCR with primer pair 5922/5923 (5’ phosphorylated) and for a second time with primer pair 5745/5746 (5’ phosphorylated), to obtain a BglBrick (50) composition with restriction sites for EcoRI, BglII; BamHI and XhoI. The vector pgRNA_bacteria was digested with EcoRI and XhoI and the fragment was ligated into the new multiple cloning site of altered pGC2 resulting in the plasmid psg-*RNA_Spy_,* which was verified by sequencing. To construct control plasmid psg-*RNA_Spy_*-2, encoding a truncated sgRNA that lacks the target specific base pairing region but contains the dCas9 binding hairpin and *S. pyogenes* terminator sequence, the region was removed from plasmid psg-*RNA_Spy_* via inverse PCR using primers 5846 and 5845. Plasmid sequence was verified.

Plasmid pBCB44 was electroporated into RN4220 and transduced into JE2 or COL. The replacement of the *spa* gene for P*_xyl/tetO3_*-*dcas9* was completed after a two-step homologous recombination event and the resulting strains BCBMS14 and BCBMS15 were verified by PCR and sequencing.

To construct plasmids expressing different sgRNAs in the *S. pyogenes* CRISPRi systems, psg-*RNA_Spy_* was used as a template and the 20 nt target specific sequence present in psg-*RNA_Spy_* was replaced by the 20 nt sequence of interest via inverse PCR as described (45). To generate the initial test sg-*RNA_Spy_* by inverse PCR, reverse primer 5846 and forward primers 5887, 5849, 5890 and 5889 (Table 3) were used for construction of psg-*fp650rodZ-Spy*1-4. Similarly reverse primer 5846 and forward primers 6424 (sg-*ftsZ*), 6426 (sg-*pbpA*) and 6423 (sg-*murJ*), were used to construct plasmids psg-*ftsZ,* psg-*pbpA* and psg-*murJ* encoding sgRNAs against selected essential genes. These three plasmids were introduced into strain BCBMS14, generating strains BCBMS16-18 and into strain BCBMS15 generating strains BCBMS20-22. For the LCML, gene-specific base pairing regions were selected by searching for PAM sequences in/upstream of essential genes/operons and the 20 nt upstream of the first PAM sequence in the coding strand of a gene were chosen as target DNA in most cases. To generate sg-*RNA_Spy_* for the library construction, we used always 5’phosporylated reverse primer 5846 together with a gene specific forward primer (see Table S2) containing the base-pairing region as an overhang in combination with the constant part of the sgRNA gttttAGAGCTAGAAATAGCAAGTTAAAATAAGGC.

**Table 2.** Strains used in this study.

| Name | Description | Reference |
| --- | --- | --- |
| NCTC8325-4 | MSSA strain NCTC8325-4 | R. Novick |
| COL | Hospital acquired homogeneous MRSA | (59) |
| JE2 | Derivative of community acquired MRSA | (17) |
| BCBMS01 | JE2 with pBCB40; expressing *S. pyogenes dcas9<sub>spy</sub>*; Kan/Neo<sup>R</sup> | This study |
| BCBMS02 | JE2 with pBCB41; expressing *S. pyogenes dcas9<sub>spy</sub>-sgfp*; Kan/Neo<sup>R</sup> | This study |
| BCBHV200 | NCTC8325-4 *ΔrodZ::fp650rodZ*; expressing *rodZ* translationally fused to fluorescent protein eqFP650 | This study |
| BCBMS04 | BCBHV200 with pCNX; serves as control; Kan/Neo<sup>R</sup> | This study |
| BCBMS05 | BCBHV200 with pBCB41, expressing *S. pyogenes dcas9<sub>spy</sub>-sgfp*; Kan/Neo<sup>R</sup> | This study |
| BCBMS06 | BCBMS04 with *psg-fp650rodZ-Spy1*; Cm<sup>R</sup>, Kan/Neo<sup>R</sup> | This study |
| BCBMS07 | BCBMS04 with *psg-fp650rodZ-Spy2*; Cm<sup>R</sup>, Kan/Neo<sup>R</sup> | This study |
| BCBMS08 | BCBMS04 with *psg-fp650rodZ-Spy3*; Cm<sup>R</sup>, Kan/Neo<sup>R</sup> | This study |
| BCBMS09 | BCBMS04 with *psg-fp650rodZ-Spy4*; Cm<sup>R</sup>, Kan/Neo<sup>R</sup> | This study |
| BCBMS10 | BCBMS05 with *psg-fp650rodZ-Spy1*; Cm<sup>R</sup>, Kan/Neo<sup>R</sup> | This study |
| BCBMS11 | BCBMS05 with *psg-fp650rodZ-Spy2*; Cm<sup>R</sup>, Kan/Neo<sup>R</sup> | This study |
| BCBMS12 | BCBMS05 with *psg-fp650rodZ-Spy3*; Cm<sup>R</sup>, Kan/Neo<sup>R</sup> | This study |
| BCBMS13 | BCBMS05 with *psg-fp650rodZ-Spy4*; Cm<sup>R</sup>, Kan/Neo<sup>R</sup> | This study |
| BCBLS01 | BCBHV200 with pBCB42; expressing *S. aureus dcas9<sub>Sa</sub>-sgfp*; Kan/Neo<sup>R</sup> | This study |
| BCBLS02 | BCBMS04 with *psg-fp650rodZ-Sa1*; Cm<sup>R</sup>, Kan/Neo<sup>R</sup> | This study |
| BCBLS03 | BCBMS04 with *psg-fp650rodZ-Sa2*; Cm<sup>R</sup>, Kan/Neo<sup>R</sup> | This study |
| BCBLS04 | BCBMS04 with *psg-fp650rodZ-Sa3*; Cm<sup>R</sup>, Kan/Neo<sup>R</sup> | This study |
| BCBLS05 | BCBLS01 with *psg-fp650rodZ-Sa1*; Cm<sup>R</sup>, Kan/Neo<sup>R</sup> | This study |
| BCBLS06 | BCBLS01 with *psg-fp650rodZ-Sa2*; Cm<sup>R</sup>, Kan/Neo<sup>R</sup> | This study |
| BCBLS07 | BCBLS01 with *psg-fp650rodZ-Sa3*; Cm<sup>R</sup>, Kan/Neo<sup>R</sup> | This study |
| BCBMS14 | JE2 *Δspa*: P<sub>xyl/tetO3</sub>-*dcas9<sub>spy</sub>*; expressing *dcas9<sub>spy</sub>* under the control of aTc-inducible promoter P<sub>xyl/tetO3</sub> from the *spa* locus | This study |
| BCBMS14<br>psg-RNA<sub>spy</sub>-2 | BCBMS14 with *psg-RNA<sub>spy</sub>-2*; Cm<sup>R</sup> | This study |
| BCBMS15 | COL *Δspa*: P<sub>xyl/tetO3</sub>-*dcas9<sub>spy</sub>*; expressing *dcas9<sub>spy</sub>* under the control of an aTc-inducible promoter P<sub>xyl/tetO3</sub> from *spa* locus | This study |
| BCBMS16 | BCBMS14 with *psg-ftsZ*; Cm<sup>R</sup> (same as LCML 261) | This study |
| BCBMS17 | BCBMS14 with *psg-pbpA*; Cm<sup>R</sup> (same as LCML 76) | This study |
| BCBMS18 | BCBMS14 with *psg-murJ*; Cm<sup>R</sup> (same as LCML 260) | This study |
| BCBMS20 | BCBMS15 with *psg-ftsZ*; Cm<sup>R</sup> | This study |
| BCBMS21 | BCBMS15 with *psg-pbpA*; Cm<sup>R</sup> | This study |
| BCBMS22 | BCBMS15 with *psg-murJ*; Cm<sup>R</sup> | This study |
| BCBMS26 | BCBHV200 with pBCB40 and *psg-fp650rodZ-Spy2*; Cm<sup>R</sup>, Kan/Neo<sup>R</sup> | This study |

**Table 3.**
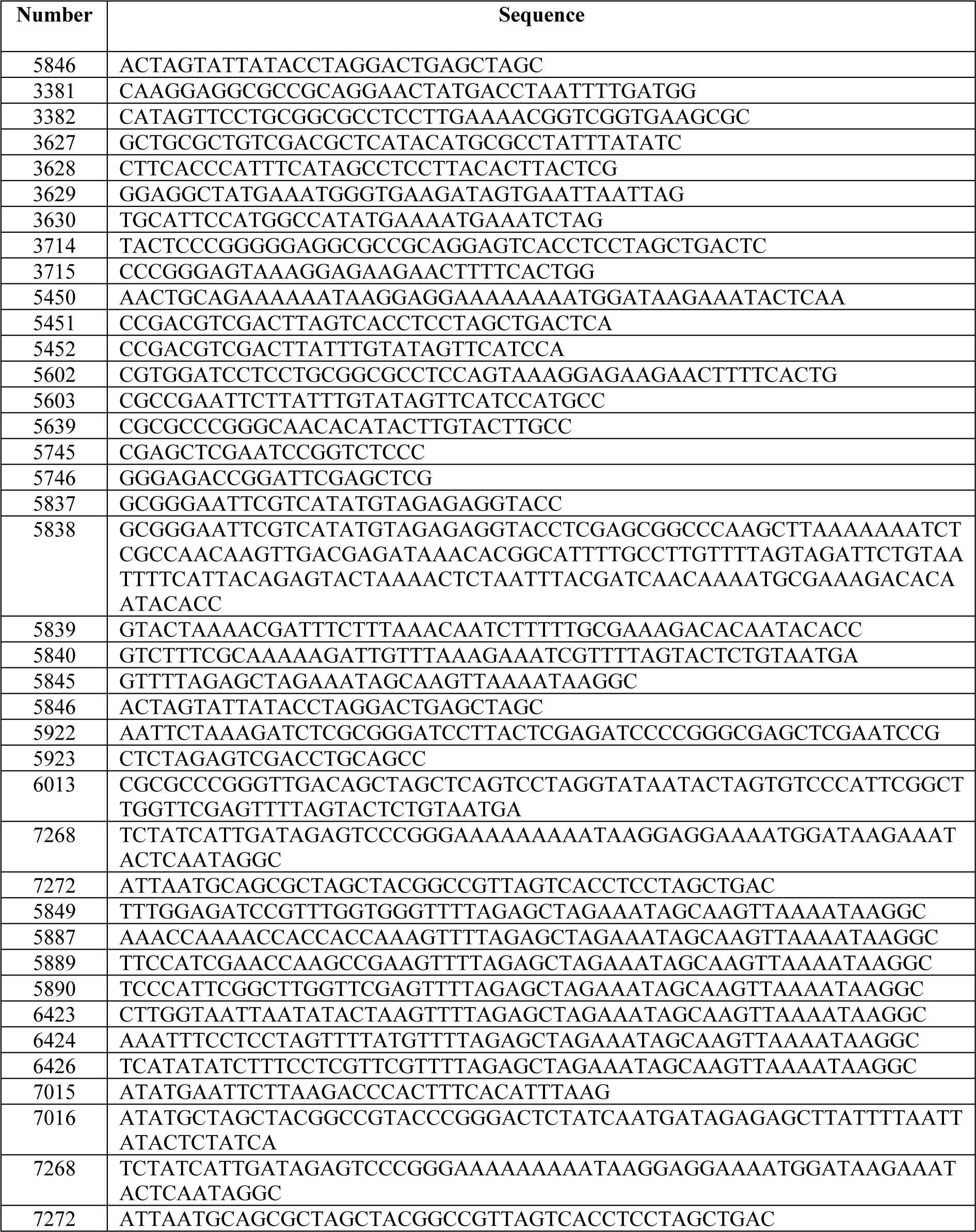
Primers used in this study.

To construct psg-*fp650rodZ-Sa1*, the *pbpb* promoter sequence (51) was amplified from NCTC8325-4 genomic DNA using primer pair 5639/5838. Primer 5838 contains the sgRNA*_Sa_* (52), a region targeting the eqFP650-RodZ specific gene sequence and a transcription terminator. The resulting PCR product was digested with SmaI and EcoRI and cloned into pGC2. Plasmid psg-*fp650rodZ-Sa2* was constructed by amplifying the *pbpb* promoter from NCTC8325-4 genomic DNA using primer pair 5639/5839 and the sgRNA*_Sa_* from *psg-fp650rodZ-Sa1* using the primer pair 5840/5837. The fragments were joined by overlap PCR with primers 5639/5837 and cloned into pCG2 using SmaI and EcoRI enzymes identical to psg-*fp650rodZ-Sa1*. For the construction of psg-*fp650rodZ-Sa3*, the *pbpb* promoter was exchanged for the synthetic minimal promoter used in psg-*RNA_Spy_* plasmids. Primer pair 6013/5837 was used to amplify the sgRNA*_Sa_* with primer 6013 containing the minimal promoter and target specific nucleotide sequence. The resulting PCR product was digested with SmaI and EcoRI and cloned into pGC2. All constructed sgRNAs were confirmed by sequencing before further utilization.

All plasmids except those encoding the sgRNA for the library were initially transformed into *E. coli* DC10B (47) with ampicillin (100 µg/ml) selection. Constructs were verified by PCR and sequencing. Plasmids were electroporated into the *S. aureus* strain RN4220 as previously described (19), followed by phage lysate preparation and transduction into other *S. aureus* strains (53). Following this protocol, the plasmids pCNX, pBCB40, pBCB41, pBCB42, psg-*fp650rodZ-Spy1-4* and psg-*fp650rodZ-Sa1-3* were electroporated into RN4220 with subsequent preparations of phage lysates. Plasmids pBCB40 and pBCB41 were transduced into JE2, generating strains BCBMS01 and BCBMS02 respectively. Strains BCBMS04, BCBMS05 and BCBLS01 were created by introducing pCNX, pBCB41 or pBCB42 respectively into the strain BCBHV200 (see below). Plasmids psg-*fp650rodZ-Spy1-4* were transduced either into BCBMS05, creating the strains BCBMS10-13, or into BCBMS04, creating control strains BCBMS06-09. Plasmids psg-*fp650rodZ-Sa1-3* were transduced either into BCBLS01, creating the strains BCBLS05-07, or into BCBMS04, creating control strains BCBLS02-04. pBCB40 and psg-*fp650rodZ-Spy2* were also transduced into BCBHV200, creating strain BCBMS26. The plasmids for the sgRNA library based on psg-*RNA_Spy_* were initially transformed into *E. coli* strain IM08B (54). IM08B has a modified DNA methylation pattern compatible with that of *S. aureus* JE2 increasing the efficiency of transformation into this strain (54). Plasmids propagated in IM08B could therefore be directly electroporated into competent BCBMS14 cells, bypassing the time-consuming steps of electroporation into RN4220 and subsequent phage transduction, frequently used to transfer plasmids into *S. aureus*. Preparation of *S. aureus* competent cells and electroporation protocol were performed as described by Johnston and colleagues (55).

We constructed a *S. aureus* strain encoding an N-terminal eqFP650-RodZ fusion as the only *rodZ* copy in the cell. For that, three DNA fragments were amplified. Fragment 1, containing a 1037 bp upstream region of *rodZ,* and fragment 3, containing a 1039 bp encompassing the *rodZ* gene and its downstream region, were amplified from NCTC8325-4 genome using the primers pairs 3627/3628 and 3382/3630, respectively. Fragment 2, encompassing the *fp650* gene without its *stop* codon and encoding a five amino acid linker (SCGAS) at the 3’ end, was amplified from plasmid pFP650-F (56) using primers 3629/3381. The three fragments were then joined by overlap PCR using primers 3627/3630 and the resulting fragment was digested with SalI and NcoI and cloned into pMAD (57). The resulting plasmid was named pMAD_*fp650rodZ* and was verified by sequencing. The pMAD_*fp650rodZ* plasmid was electroporated into RN4220 at 30 °C and subsequently transduced to NCTC8325-4 (selection at 30 °C with erythromycin in both steps). The replacement of the *rodZ* gene for *fp650-rodZ* was completed after a two-step homologous recombination event and was confirmed by PCR. The strain was named BCBHV200.

### Fluorescence microscopy NCTC *fp650-rodZ* conditional mutant

Overnight cultures of strains expressing *dcas9-sgfp* (from *S. pyogenes* or *S. aureus*) with corresponding sgRNA targeting *fp650-rodZ* (BCBMS10-13; BCBLS05-07) and control strains (BCBMS06-09; BCBLS02-04, lacking *dcas9*) were back-diluted 1:500 in 10 ml of TSB, with 10 µg/ml Cm, 50 µg/ml of both Neo and Kan and 0.1 µM CdCl_2_ and incubated at 37 °C, with aeration, to OD_600nm_ of 0.8. A 1 ml aliquot of culture was pelleted and re-suspended in 30 µl of phosphate buffered saline (PBS). A 5 µl sample of each dCas9-producing culture expressing a specific sgRNA, was mixed with an equal volume of the corresponding control culture lacking dCas9 but expressing the same sgRNA (see experimental setup in Figure 2B). 1 µl of the mixed culture was placed on a thin layer of 1.2% agarose in PBS and imaged using a Axio Observer.Z1 microscope equipped with a Photometrics CoolSNAP HQ2 camera (Roper Scientific), using phase contrast objective Plan Apo 100 x/1.4 oil Ph3. Cells were imaged using the ZEN software (Zeiss). The median fluorescence of sGFP and eqFP650 in each cell was determined using eHooke (58).

### Microscopy analysis of conditional mutants in essential genes

Overnight cultures of derivatives of BCBMS14 (JE2 background) expressing sgRNAs against *ftsZ* (BCBMS16), *pbpA* (BCBMS17) or *murJ* (BCBMS18) were each back-diluted 1:1000 into fresh media containing 10 µg/ml Cm with (induced) or without (non-induced) 100 ng/ml aTc and grown at 37 °C. After 2 hours, a second 1:100 dilution of each culture was made and cells were grown to exponential phase (OD_600nm_ 0.8). A 1 ml aliquot of each culture was incubated with 2.5 µg/ml Nile Red (membrane stain, Invitrogen) for 5 min at 37 °C with shaking. The culture was then pelleted and resuspended in 30 µl PBS. 1 µl of cell culture was placed on a thin layer of 1.2% agarose in PBS and imaged as described above.

### Western blot

*S. aureus* strains were grown overnight, diluted 1:500 in fresh medium and incubated at 37 °C with aeration. When necessary, cultures were supplemented with required antibiotics and aTc or CdCl_2_. At OD_600nm_ of approximately 0.6, cells were harvested and broken with glass beads in a SpeedMill (Analytik Jena). Samples were centrifuged for 10 min at 16000 xg and resuspended in 300 µl PBS containing protease inhibitors (cOmplete, Roche). A 16 µl sample was added to 5 µl of 5x SDS sample buffer (300mM Tris-HCl, pH 6.8, 50% glycerol, 10% SDS, 0.01% Bromophenol Blue, 10% Beta-mercaptoethanol), boiled for 5 minutes and samples were separated by SDS-PAGE (12% gel) at 120V. Proteins were then transferred to Hybond-P PVDF membrane (GE Healthcare) using a BioRad Transblot Turbo transfer system, according to the manufacturer’s instructions. After transfer, the membrane was cut at the 100 kDa marker band. The upper part was used for protein detection using a specific monoclonal antibody against Cas9 (1:1000 dilution, #MAC133, Sigma-Aldrich) and Alexa555 secondary antibody for fluorescence detection (iBright, ThermoFisher), while the lower part of the membrane was stained with Sypro Ruby Protein Blot Stain (Thermo Fisher), a total protein stain for western blot normalization.

### Growth curves

For analysis of growth of the CRISPRi conditional mutants of BCBMS20-22 (COL with aTc-inducible CRISPRi), cells were grown overnight at 37 °C in TSB medium supplemented with 10 µg/mL Cm and diluted 1:500 into 100 ml Erlenmeyer flasks containing fresh TSB medium with 10 µg/ml Cm, with or without 100 ng/ ml aTc which induces expression of *dcas9*. After four hours, cultures were diluted a second time 1:100 in fresh medium and growth was followed by measuring the OD_600nm_ every hour. All cultures were incubated at 37 °C with agitation and the OD_600nm_ was recorded.

Analysis of the library of CRISPRi conditional mutants (derived from strain BCBMS14) was performed in a 96-well plate reader (Biotek Synergy Neo2). Overnight cultures were diluted 1:1000 in fresh media containing 10 µg/ml Cm. Samples (200 µl) of each culture, with or without the addition of 200 ng/ml aTc inducer, were added to wells in a 96-well plate. Plates were incubated at 30 °C with shaking for 120 min, the OD_600nm_ was measured every 30 minutes. After two hours, the samples were diluted 1:100 into fresh media (with 10 µg/ml Cm), containing or not aTc and growth was followed in all cultures for a further 630 min. For CRISPRi conditional mutants showing less than 50% growth reduction at 30 °C, growth analysis was also performed at 37 °C.

### Data availability

Data will be made available upon reasonable request to the corresponding author.

## Acknowledgments

We thank Vincent de Bakker for advice on statistical analysis. This study was funded by the European Research Council through grant ERC-2017-CoG-771709 (to MGP), by national funds through FCT– Fundação para a Ciência e a Tecnologia through MOSTMICRO-ITQB R&D Unit (UIDB/04612/2020, UIDP/04612/2020 to ITQB) and LS4FUTURE Associated Laboratory (LA/P/0087/2020 to ITQB) and FCT fellowships PD/BD/128408/2017 (to M.S.) and 2022.12215.BD (to D.A.); by the European Union’s Horizon 2020 research and innovation programme under the Marie Sklodowska-Curie grant agreement No 839596 (to SS) and by the European Molecular Biology Organization through award ALTF 673-2018 (to SS). Figure 1 was created with Biorender.com.

## Notes

### Competing Interest Statement

The authors have declared no competing interest.

### Summary of Updates

A more complete CRISPRi library is included in the revised version

