## Supplemental Tables and Figures for "A CRISPRi-based genetic resource to study essential *Staphylococcus aureus* genes"

**Supplementary Figures**

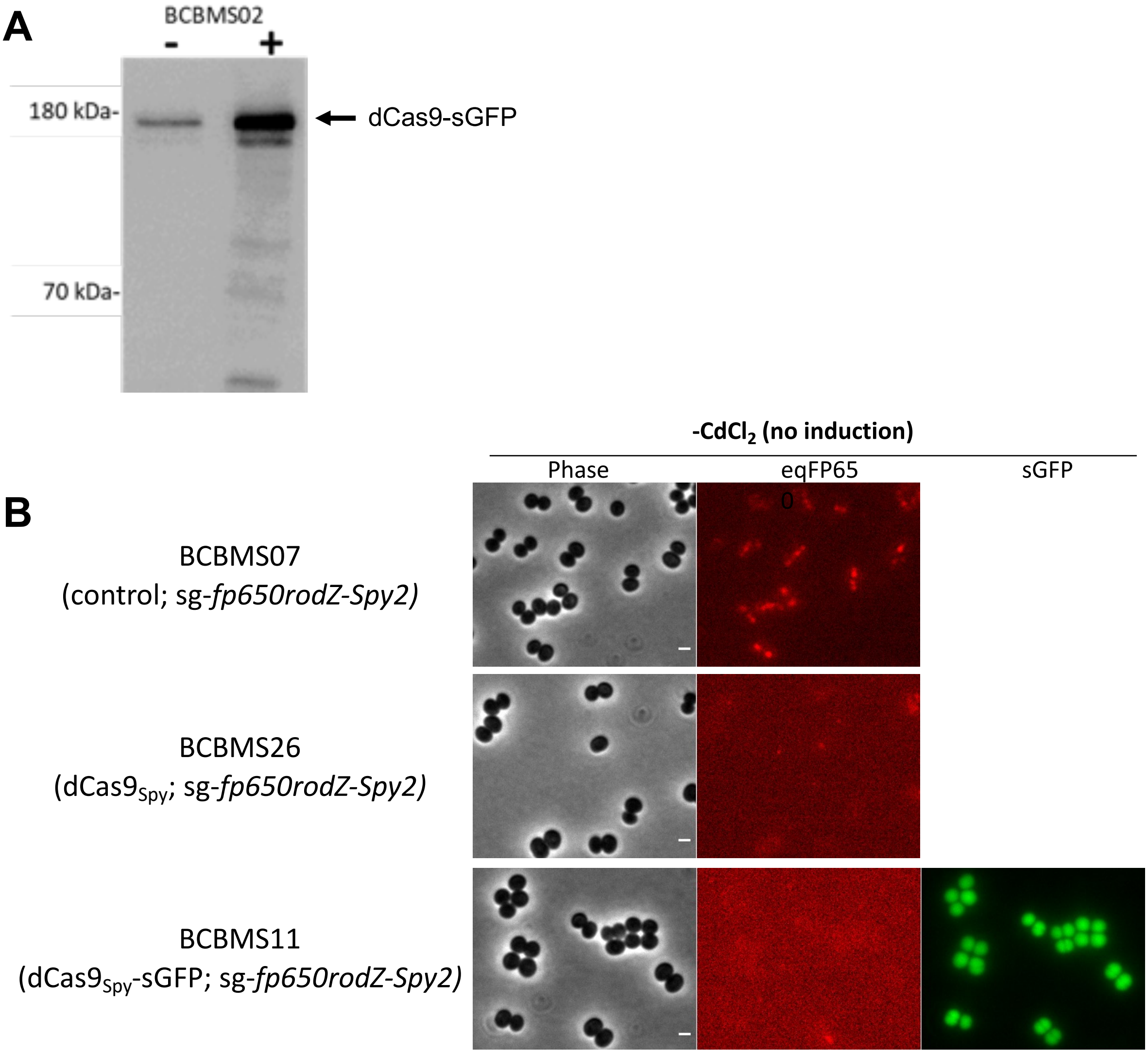

**Figure S1. Plasmid-based CRISPRi system allows leaky production of dCas9 in the absence of inducer.** (**A**) Western Blot using anti-Cas9 antibody of protein extracts from non-induced (-) and induced (+, 0.1 µM $CdCl_2$) strain BCBMS02 showing that production of dCas9-sGFP increases in the presence of $CdCl_2$, but is also present in the absence of the inducer. (**B**) Strains BCBMS26 and BCBMS11 showed depletion of eqFP650-RodZ even when *dCas9* expression was not induced, indicating leakiness of the cadmium-inducible promoter. The control strain lacking dCas9 (BCBMS07) shows eqFP650-RodZ localized at the septum.

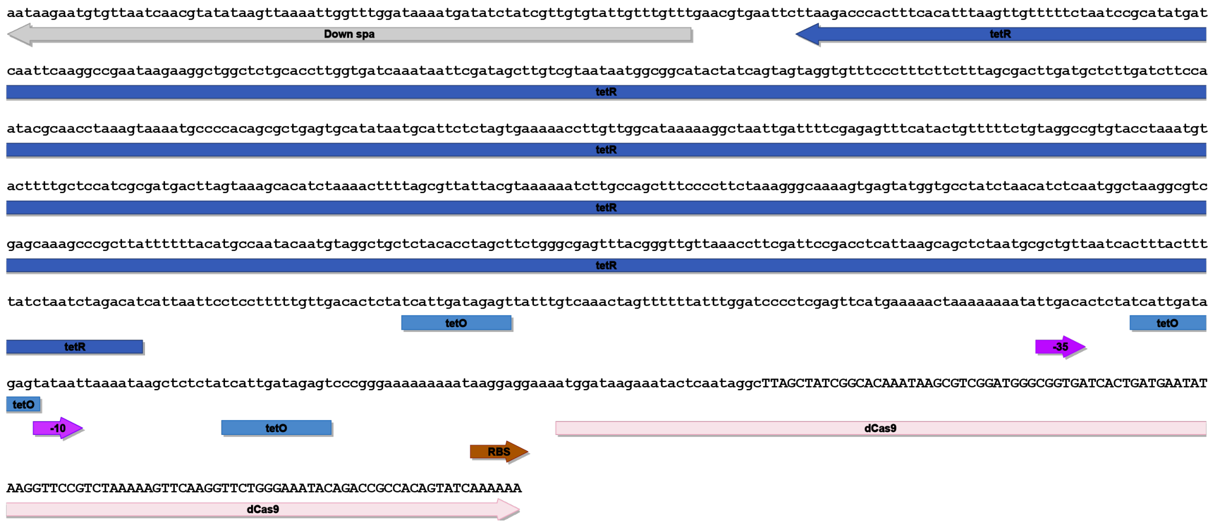

**Figure S2. Nucleotide sequence of P$_{xyl/tetO3}$ and *tetR* gene introduced in the *spa* locus of strains BCBMS14/15.** The promoter is based on the tetracycline-inducible *xyl/tetO* promoter with two additional *tetO* sites. In addition, a weak ribosome binding site (RBS) was used to reduce translation of *dCas9*.

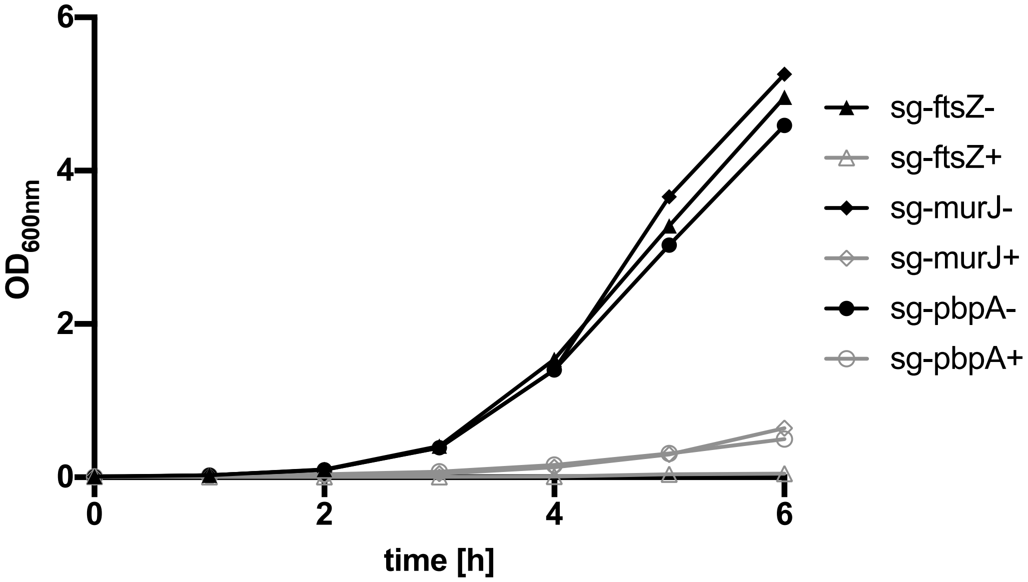

**Figure S3. CRISPRi system with chromosome-encoded *dCas9$_{Spy}$* is suitable to target essential genes in *S. aureus* strain COL.** Growth curves of strain BCBMS15 expressing sgRNAs targeting the essential genes *ftsZ* (BCBMS20), *pbpA* (BCBMS 21) and *murJ* (BCBMS 22) in the absence (-) or the presence (+) of 100 ng/ml aTc (inducer for *dCas9* expression).

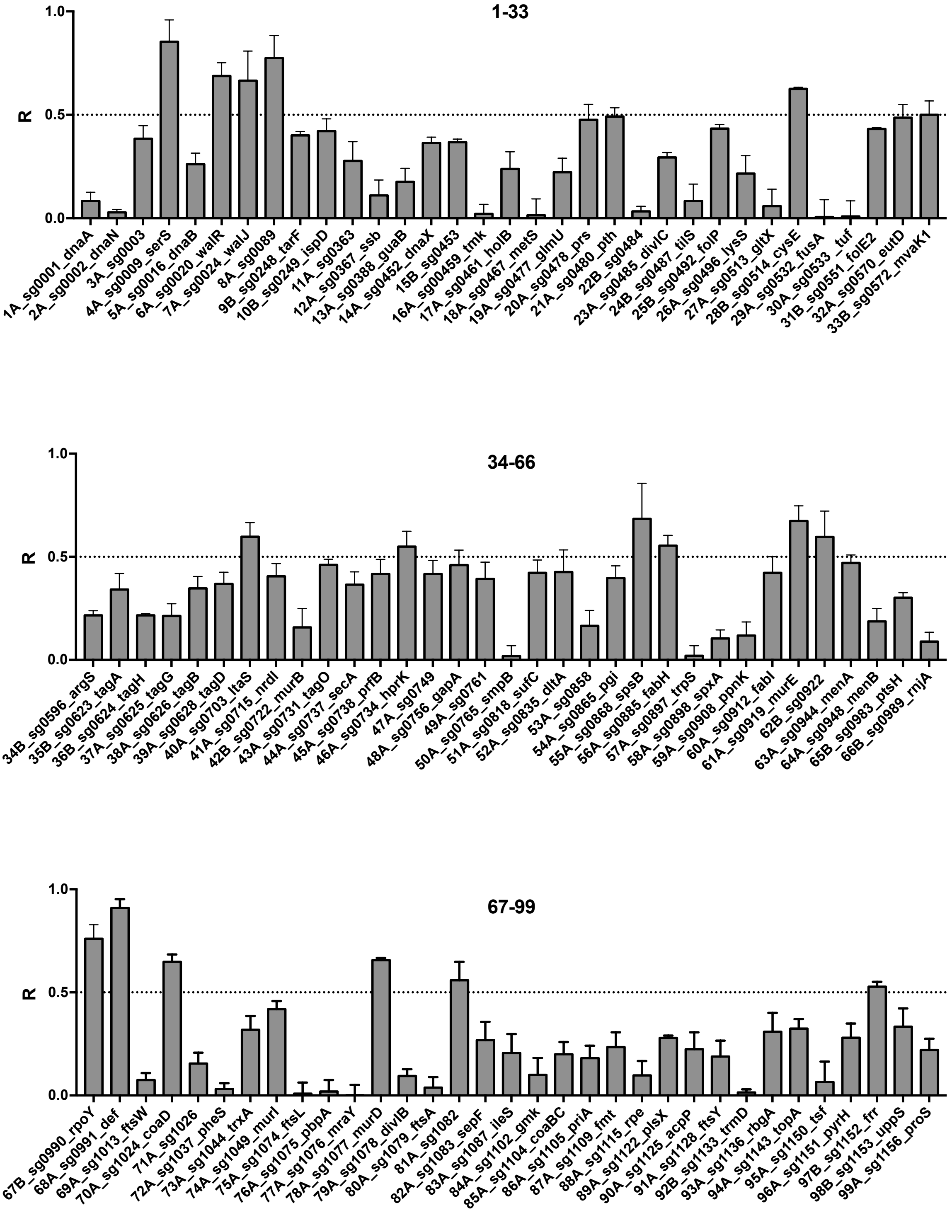

**Figure S4. Evaluation of growth inhibition in mutants from the Lisbon CRISPRi Mutant Library.** Graphs show values of R (ratio of area under the curve of growth curves obtained in the presence versus the absence of inducer aTc) for the 261 strains of the LCML containing sgRNAs targeting a predicted essential gene/operon. For each gene/operon data shown corresponds to the strain containing the sgRNA, A or B, with highest efficiency. Error bars represent SEM from three independent experiments.

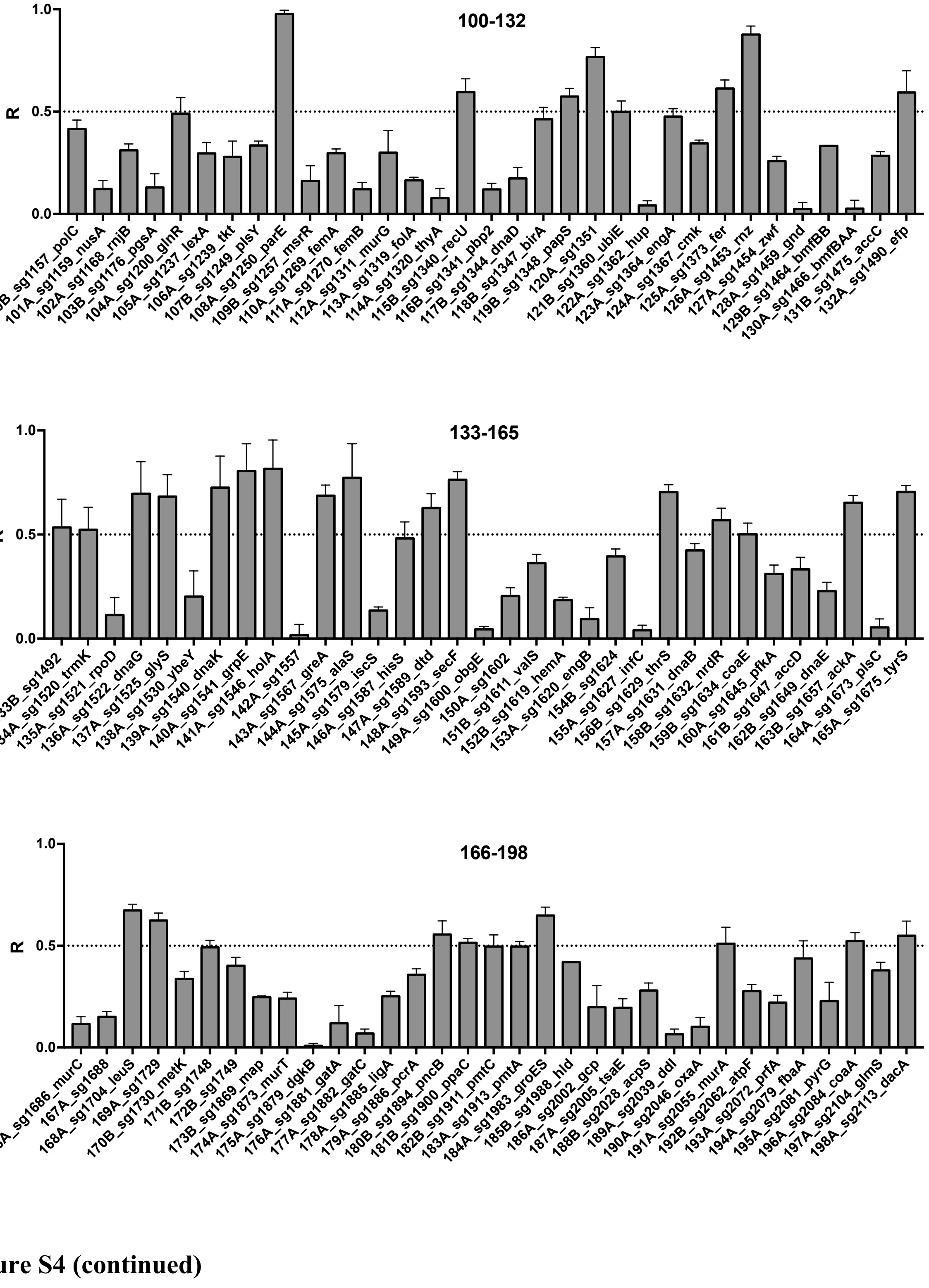

**Figure S4 (continued)**

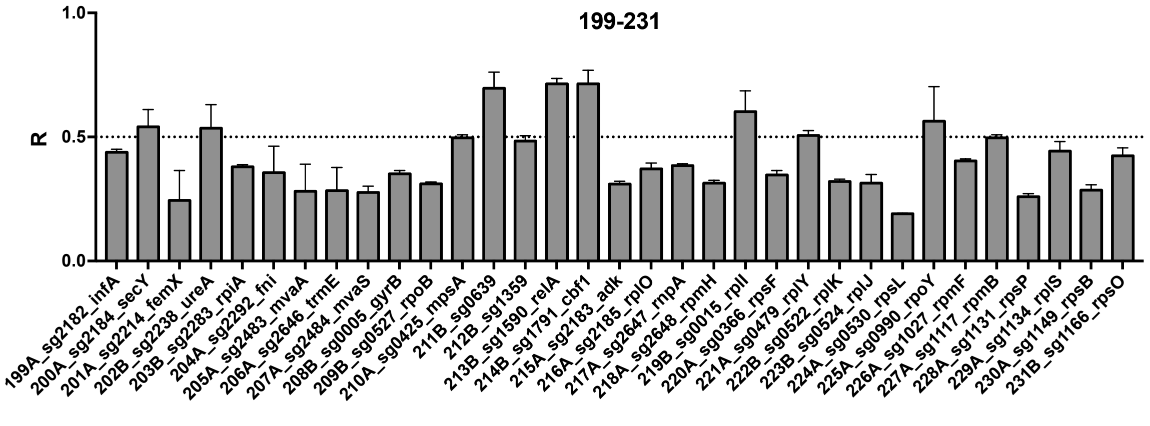

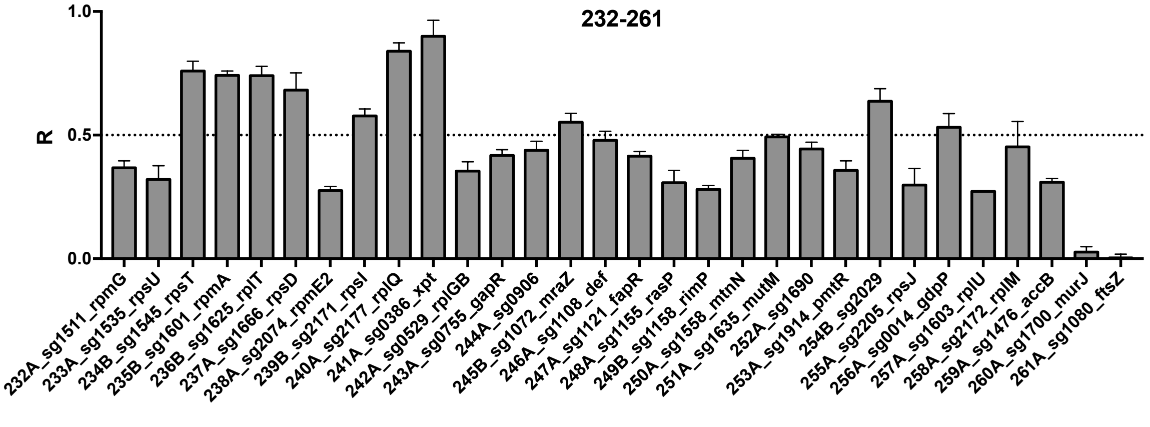

**Figure S4 (continued)**

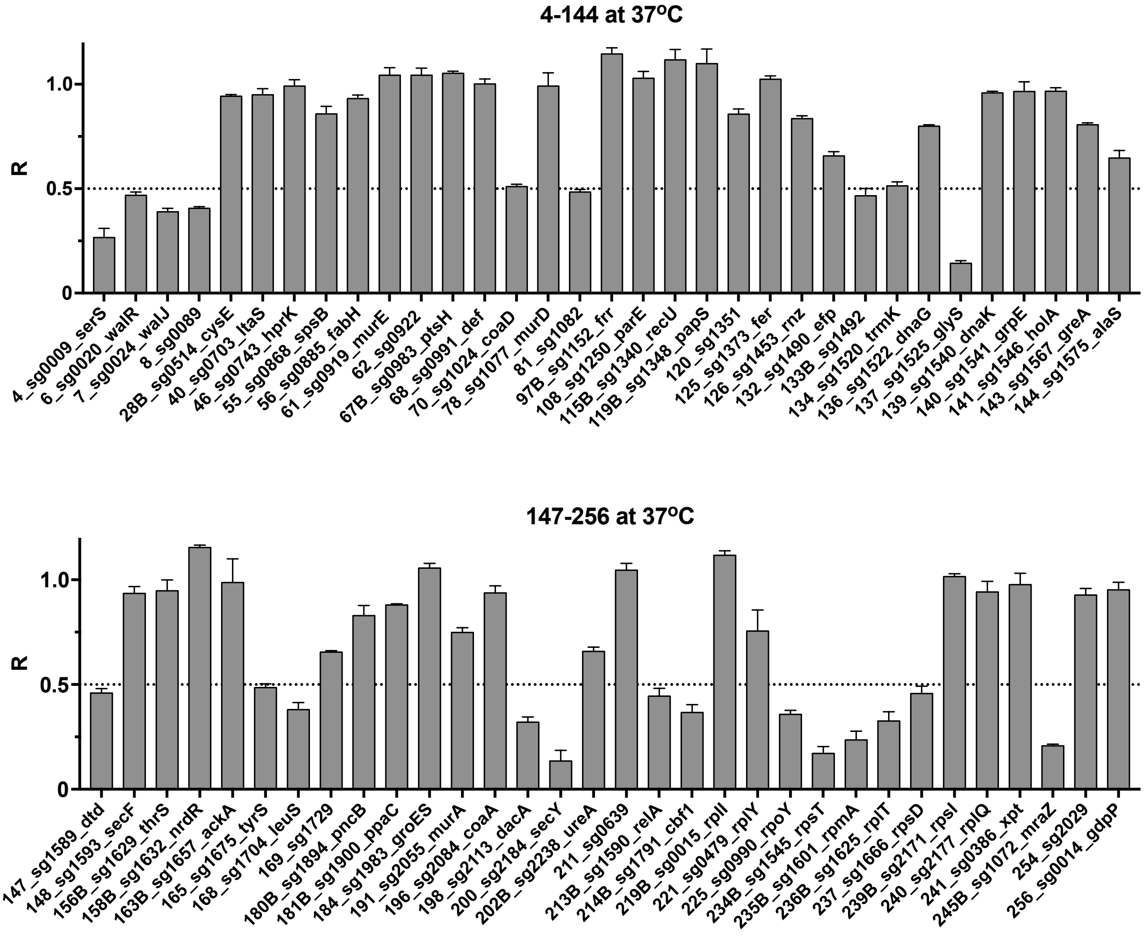

**Figure S5. Evaluation of growth inhibition at 37 ºC in selected mutants from the Lisbon CRISPRi Mutant Library.** Growth in the presence and absence of aTc of LCML strains that showed R higher than 0.5 when growth inhibition was tested at 30 ºC (see Figure S4) was also evaluated at 37 ºC**.** R is ratio of the area under the curve of the growth curve obtained in the presence versus the absence of inducer aTc. Error bars represent SEM from three independent experiments.

### Supplementary Table 1. Essential operons identified in *Staphylococcus aureus*

| Operon Number | Gene position in operon | Locus Tag | Gene Name |
|---|---|---|---|
| **1** | 1 | SAUSA300_0001 | ***dnaA*** |
| **2** | 1 | SAUSA300_0002 | ***dnaN*** |
| **3** | 1 | SAUSA300_0003 | ***0003*** |
| **3** | 2 | SAUSA300_0004 | <u>*recF*</u> |
| **3** | 3 | SAUSA300_0005 | ***gyrB*** |
| **3** | 4 | SAUSA300_0006 | *gyrA* |
| **4** | 1 | SAUSA300_0009 | ***serS*** |
| **5** | 1 | SAUSA300_0013 | <u>*0013*</u> |
| **5** | 2 | SAUSA300_0014 | ***gdpP*** |
| **5** | 3 | SAUSA300_0015 | ***rplI*** |
| **5** | 4 | SAUSA300_0016 | ***dnaB*** |
| **6** | 1 | SAUSA300_0020 | ***walR*** |
| **6** | 2 | SAUSA300_0021 | *walK* |
| **6** | 3 | SAUSA300_0022 | <u>*walH*</u> |
| **6** | 4 | SAUSA300_0023 | <u>*walI*</u> |
| **7** | 1 | SAUSA300_0024 | ***walJ*** |
| **8** | 1 | SAUSA300_0089 | ***0089*** |
| **9** | 1 | SAUSA300_0248 | ***tarF*** |
| **10** | 1 | SAUSA300_0249 | ***tarI / ispD*** |
| **10** | 2 | SAUSA300_0250 | *tarJ* |
| **10** | 3 | SAUSA300_0251 | *tarL* |
| **10** | 4 | SAUSA300_0252 | <u>*tarS*</u> |
| **11** | 1 | SAUSA300_0363 | ***0363*** |
| **12** | 1 | SAUSA300_0366 | ***rpsF*** |
| **12** | 2 | SAUSA300_0367 | ***ssb*** |
| **12** | 3 | SAUSA300_0368 | *rpsR* |
| **13** | 1 | SAUSA300_0386 | ***xpt*** |
| **13** | 2 | SAUSA300_0387 | <u>*pbuX*</u> |
| **13** | 3 | SAUSA300_0388 | ***guaB*** |
| **13** | 4 | SAUSA300_0389 | *guaA* |
| **14** | 1 | SAUSA300_0425 | ***mpsA*** |
| **14** | 2 | SAUSA300_0426 | *mpsB* |
| **15** | 1 | SAUSA300_0452 | ***dnaX*** |
| **16** | 1 | SAUSA300_0453 | ***0453*** |
| **16** | 2 | SAUSA300_0454 | *recR* |
| **17** | 1 | SAUSA300_0458 | <u>*0458*</u> |
| **17** | 2 | SAUSA300_0459 | ***tmk*** |
| **17** | 3 | SAUSA300_0460 | *pstA* |
| **18** | 1 | SAUSA300_0461 | ***holB*** |
| **18** | 2 | SAUSA300_0462 | <u>*0462*</u> |
| **18** | 3 | SAUSA300_0463 | *0463* |
| **19** | 1 | SAUSA300_0467 | ***metS*** |
| **19** | 2 | SAUSA300_0468 | <u>*0468*</u> |

| Operon Number | Gene position in operon | Locus Tag | Gene Name |
|---|---|---|---|
| **20** | 1 | SAUSA300_0477 | ***glmU*** |
| **21** | 1 | SAUSA300_0478 | ***prs*** |
| **22** | 1 | SAUSA300_0479 | ***rplY*** |
| **22** | 2 | SAUSA300_0480 | ***pth*** |
| **22** | 3 | SAUSA300_0481 | *mfd* |
| **22** | 4 | SAUSA300_0482 | *0482* |
| **22** | 5 | SAUSA300_0483 | *0483* |
| **22** | 6 | SAUSA300_0484 | ***0484*** |
| **22** | 7 | SAUSA300_0485 | ***divIC*** |
| **23** | 1 | SAUSA300_0487 | ***tilS*** |
| **24** | 1 | SAUSA300_0492 | ***folP*** |
| **24** | 2 | SAUSA300_0493 | *folB* |
| **24** | 3 | SAUSA300_0494 | *folK* |
| **25** | 1 | SAUSA300_0496 | ***lysS*** |
| **26** | 1 | SAUSA300_0513 | ***gltX*** |
| **27** | 1 | SAUSA300_0514 | ***cysE*** |
| **27** | 2 | SAUSA300_0515 | *cysS* |
| **27** | 3 | SAUSA300_0516 | *0516* |
| **27** | 4 | SAUSA300_0517 | *0517* |
| **27** | 5 | SAUSA300_0518 | *0518* |
| **28** | 1 | SAUSA300_0522 | ***rplK*** |
| **28** | 2 | SAUSA300_0523 | *rplA* |
| **29** | 1 | SAUSA300_0524 | ***rplJ*** |
| **29** | 2 | SAUSA300_0525 | *rplL* |
| **30** | 1 | SAUSA300_0527 | ***rpoB*** |
| **30** | 2 | SAUSA300_0528 | *rpoC* |
| **31** | 1 | SAUSA300_0529 | ***rplGB*** |
| **31** | 2 | SAUSA300_0530 | ***rpsL*** |
| **31** | 3 | SAUSA300_0531 | *0531* |
| **31** | 4 | SAUSA300_0532 | ***fusA*** |
| **31** | 5 | SAUSA300_0533 | ***tuf*** |
| **32** | 1 | SAUSA300_0553 | *0553* |
| **32** | 2 | SAUSA300_0552 | *0552* |
| **32** | 3 | SAUSA300_0551 | ***folE2*** |
| **33** | 1 | SAUSA300_0570 | ***eutD*** |
| **33** | 2 | SAUSA300_0571 | *lipL* |
| **34** | 1 | SAUSA300_0572 | ***mvaK1*** |
| **34** | 2 | SAUSA300_0573 | *mvaD* |
| **34** | 3 | SAUSA300_0574 | *mvaK2* |
| **35** | 1 | SAUSA300_0595 | *0595* |
| **35** | 2 | SAUSA300_0596 | ***argS*** |
| **36** | 1 | SAUSA300_0623 | ***tagA*** |
| **37** | 1 | SAUSA300_0624 | ***tagH*** |
| **38** | 1 | SAUSA300_0625 | ***tagG*** |
| **38** | 2 | SAUSA300_0626 | ***tagB*** |

| Operon Number | Gene position in operon | Locus Tag | Gene Name |
|---|---|---|---|
| **38** | 3 | SAUSA300_0627 | *tagX* |
| **39** | 1 | SAUSA300_0628 | ***tagD*** |
| **40** | 1 | SAUSA300_0636 | *dhaK* |
| **40** | 2 | SAUSA300_0637 | *dhaI* |
| **40** | 3 | SAUSA300_0638 | *dhaM* |
| **40** | 4 | SAUSA300_0639 | ***0639*** |
| **41** | 1 | SAUSA300_0703 | ***ltaS*** |
| **42** | 1 | SAUSA300_0715 | ***nrdI*** |
| **42** | 2 | SAUSA300_0716 | *nrdE* |
| **42** | 3 | SAUSA300_0717 | *nrdF* |
| **43** | 1 | SAUSA300_0722 | ***murB*** |
| **44** | 1 | SAUSA300_0731 | ***tagO*** |
| **45** | 1 | SAUSA300_0737 | ***secA*** |
| **46** | 1 | SAUSA300_0738 | ***prfB*** |
| **47** | 1 | SAUSA300_0743 | ***hprK*** |
| **47** | 2 | SAUSA300_0744 | *lgt* |
| **47** | 3 | SAUSA300_0745 | *0745* |
| **47** | 4 | SAUSA300_0746 | *0746* |
| **47** | 5 | SAUSA300_0747 | *trxB* |
| **48** | 1 | SAUSA300_0748 | *0748* |
| **48** | 2 | SAUSA300_0749 | ***0749*** |
| **48** | 3 | SAUSA300_0750 | *0750* |
| **49** | 1 | SAUSA300_0755 | ***gapR*** |
| **49** | 2 | SAUSA300_0756 | ***gapA*** |
| **49** | 3 | SAUSA300_0757 | *pgk* |
| **49** | 4 | SAUSA300_0758 | *tpiA* |
| **49** | 5 | SAUSA300_0759 | *pgm* |
| **49** | 6 | SAUSA300_0760 | *eno* |
| **50** | 1 | SAUSA300_0761 | ***0761*** |
| **51** | 1 | SAUSA300_0763 | *est* |
| **51** | 2 | SAUSA300_0764 | *rnr* |
| **51** | 3 | SAUSA300_0765 | ***smpB*** |
| **52** | 1 | SAUSA300_0818 | ***sufC*** |
| **52** | 2 | SAUSA300_0819 | *sufD* |
| **52** | 3 | SAUSA300_0820 | *sufS* |
| **52** | 4 | SAUSA300_0821 | *sufU* |
| **52** | 5 | SAUSA300_0822 | *sufB* |
| **53** | 1 | SAUSA300_0835 | ***dltA*** |
| **53** | 2 | SAUSA300_0836 | *dltB* |
| **53** | 3 | SAUSA300_0837 | *dltC* |
| **53** | 4 | SAUSA300_0838 | *dltD* |
| **54** | 1 | SAUSA300_0858 | ***0858*** |
| **55** | 1 | SAUSA300_0865 | ***pgi*** |
| **56** | 1 | SAUSA300_0866 | *0866* |
| **56** | 2 | SAUSA300_0867 | *spsA* |

| Operon Number | Gene position in operon | Locus Tag | Gene Name |
|---|---|---|---|
| **56** | 3 | SAUSA300_0868 | ***spsB*** |
| **57** | 1 | SAUSA300_0885 | ***fabH*** |
| **57** | 2 | SAUSA300_0886 | *fabF* |
| **58** | 1 | SAUSA300_0897 | ***trpS*** |
| **59** | 1 | SAUSA300_0898 | ***spxA*** |
| **60** | 1 | SAUSA300_0906 | ***0906*** |
| **60** | 2 | SAUSA300_0907 | *relQ* |
| **60** | 3 | SAUSA300_0908 | ***ppnK*** |
| **60** | 4 | SAUSA300_0909 | *0909* |
| **60** | 5 | SAUSA300_0910 | *mgtE* |
| **60** | 6 | SAUSA300_0911 | *cpaA* |
| **61** | 1 | SAUSA300_0912 | ***fabI*** |
| **62** | 1 | SAUSA300_0919 | ***murE*** |
| **62** | 2 | SAUSA300_0920 | *0920* |
| **62** | 3 | SAUSA300_0921 | *prfC* |
| **63** | 1 | SAUSA300_0922 | ***0922*** |
| **64** | 1 | SAUSA300_0944 | ***menA*** |
| **65** | 1 | SAUSA300_0945 | *0945* |
| **65** | 2 | SAUSA300_0946 | *menD* |
| **65** | 3 | SAUSA300_0947 | *0947* |
| **65** | 4 | SAUSA300_0948 | ***menB*** |
| **66** | 1 | SAUSA300_0983 | ***ptsH*** |
| **66** | 2 | SAUSA300_0984 | *ptsI* |
| **67** | 1 | SAUSA300_0989 | ***rnjA*** |
| **67** | 2 | SAUSA300_0990 | ***rpoY*** |
| **68** | 1 | SAUSA300_0991 | ***def*** |
| **69** | 1 | SAUSA300_1013 | ***ftsW*** |
| **70** | 1 | SAUSA300_1023 | *rsmD* |
| **70** | 2 | SAUSA300_1024 | ***coaD*** |
| **71** | 1 | SAUSA300_1026 | ***1026*** |
| **71** | 2 | SAUSA300_1027 | ***rpmF*** |
| **72** | 1 | SAUSA300_1037 | ***pheS*** |
| **72** | 2 | SAUSA300_1038 | *pheT* |
| **73** | 1 | SAUSA300_1044 | ***trxA*** |
| **74** | 1 | SAUSA300_1049 | ***murI*** |
| **74** | 2 | SAUSA300_1050 | *1050* |
| **74** | 3 | SAUSA300_1051 | *1051* |
| **75** | 1 | SAUSA300_1072 | ***mraZ*** |
| **75** | 2 | SAUSA300_1073 | *mraW* |
| **75** | 3 | SAUSA300_1074 | ***ftsL*** |
| **75** | 4 | SAUSA300_1075 | ***pbpA*** |
| **76** | 1 | SAUSA300_1076 | ***mraY*** |
| **76** | 2 | SAUSA300_1077 | ***murD*** |
| **76** | 3 | SAUSA300_1078 | ***divIB*** |
| **76** | 4 | SAUSA300_1079 | ***ftsA*** |

| Operon Number | Gene position in operon | Locus Tag | Gene Name |
|---|---|---|---|
| **76** | 5 | SAUSA300_1080 | ***ftsZ*** |
| **77** | 1 | SAUSA300_1081 | <u>*1081*</u> |
| **77** | 2 | SAUSA300_1082 | ***1082*** |
| **77** | 3 | SAUSA300_1083 | ***sepF*** |
| **77** | 4 | SAUSA300_1084 | <u>*1084*</u> |
| **77** | 5 | SAUSA300_1085 | <u>*1085*</u> |
| **77** | 6 | SAUSA300_1086 | *1086* |
| **78** | 1 | SAUSA300_1087 | ***ileS*** |
| **79** | 1 | SAUSA300_1102 | ***gmk*** |
| **79** | 2 | SAUSA300_1103 | *rpoZ* |
| **80** | 1 | SAUSA300_1104 | ***coaBC*** |
| **80** | 2 | SAUSA300_1105 | ***priA*** |
| **81** | 1 | SAUSA300_1108 | ***def*** |
| **81** | 2 | SAUSA300_1109 | ***fmt*** |
| **81** | 3 | SAUSA300_1110 | <u>*sun*</u> |
| **81** | 4 | SAUSA300_1111 | <u>*rlmN*</u> |
| **81** | 5 | SAUSA300_1112 | <u>*stpI*</u> |
| **81** | 6 | SAUSA300_1113 | <u>*pknB*</u> |
| **82** | 1 | SAUSA300_1114 | <u>*rsgA*</u> |
| **82** | 2 | SAUSA300_1115 | ***rpe*** |
| **82** | 3 | SAUSA300_1116 | *thiN* |
| **83** | 1 | SAUSA300_1117 | ***rpmB*** |
| **84** | 1 | SAUSA300_1121 | ***fapR*** |
| **84** | 2 | SAUSA300_1122 | ***plsX*** |
| **84** | 3 | SAUSA300_1123 | *fabD* |
| **84** | 4 | SAUSA300_1124 | *fabG* |
| **85** | 1 | SAUSA300_1125 | ***acpP*** |
| **86** | 1 | SAUSA300_1127 | <u>*smc*</u> |
| **86** | 2 | SAUSA300_1128 | ***ftsY*** |
| **86** | 3 | SAUSA300_1129 | *1129* |
| **86** | 4 | SAUSA300_1130 | *ffh* |
| **87** | 1 | SAUSA300_1131 | ***rpsP*** |
| **87** | 2 | SAUSA300_1132 | *rimM* |
| **87** | 3 | SAUSA300_1133 | ***trmD*** |
| **87** | 4 | SAUSA300_1134 | ***rplS*** |
| **88** | 1 | SAUSA300_1136 | ***rbgA*** |
| **88** | 2 | SAUSA300_1137 | *rnhB* |
| **89** | 1 | SAUSA300_1142 | <u>*dprA*</u> |
| **89** | 2 | SAUSA300_1143 | ***topA*** |
| **90** | 1 | SAUSA300_1149 | ***rpsB*** |
| **90** | 2 | SAUSA300_1150 | ***tsf*** |
| **91** | 1 | SAUSA300_1151 | ***pyrH*** |
| **91** | 2 | SAUSA300_1152 | ***frr*** |
| **92** | 1 | SAUSA300_1153 | ***uppS*** |
| **92** | 2 | SAUSA300_1154 | *cdsA* |

| Operon Number | Gene position in operon | Locus Tag | Gene Name |
|---|---|---|---|
| **93** | 1 | SAUSA300_1155 | ***rasP*** |
| **93** | 2 | SAUSA300_1156 | ***proS*** |
| **94** | 1 | SAUSA300_1157 | ***polC*** |
| **95** | 1 | SAUSA300_1158 | ***rimP*** |
| **95** | 2 | SAUSA300_1159 | ***nusA*** |
| **95** | 3 | SAUSA300_1160 | *1160* |
| **95** | 4 | SAUSA300_1161 | *1161* |
| **95** | 5 | SAUSA300_1162 | *infB* |
| **96** | 1 | SAUSA300_1166 | ***rpsO*** |
| **97** | 1 | SAUSA300_1168 | ***rnjB*** |
| **98** | 1 | SAUSA300_1169 | <u>*ftsK*</u> |
| **98** | 2 | SAUSA300_1170 | <u>*1170*</u> |
| **98** | 3 | SAUSA300_1171 | <u>*1171*</u> |
| **98** | 4 | SAUSA300_1172 | <u>*1172*</u> |
| **98** | 5 | SAUSA300_1173 | <u>*1173*</u> |
| **98** | 6 | SAUSA300_1174 | <u>*1174*</u> |
| **98** | 7 | SAUSA300_1175 | ***rodZ*** |
| **98** | 8 | SAUSA300_1176 | ***pgsA*** |
| **99** | 1 | SAUSA300_1200 | ***glnR*** |
| **99** | 2 | SAUSA300_1201 | *glnA* |
| **100** | 1 | SAUSA300_1237 | ***lexA*** |
| **101** | 1 | SAUSA300_1239 | ***tkt*** |
| **102** | 1 | SAUSA300_1249 | ***plsY*** |
| **103** | 1 | SAUSA300_1250 | ***parE*** |
| **103** | 2 | SAUSA300_1251 | *parC* |
| **104** | 1 | SAUSA300_1257 | ***msrR*** |
| **105** | 1 | SAUSA300_1269 | ***femA*** |
| **105** | 2 | SAUSA300_1270 | ***femB*** |
| **106** | 1 | SAUSA300_1312 | <u>*1312*</u> |
| **106** | 2 | SAUSA300_1311 | ***murG*** |
| **106** | 3 | SAUSA300_1310 | *1310* |
| **107** | 1 | SAUSA300_1319 | ***folA*** |
| **107** | 2 | SAUSA300_1318 | *1318* |
| **108** | 1 | SAUSA300_1320 | ***thyA*** |
| **109** | 1 | SAUSA300_1340 | ***recU*** |
| **109** | 2 | SAUSA300_1341 | ***pbp2*** |
| **110** | 1 | SAUSA300_1344 | ***dnaD*** |
| **110** | 2 | SAUSA300_1343 | *nth* |
| **110** | 3 | SAUSA300_1342 | <u>*1342*</u> |
| **111** | 1 | SAUSA300_1349 | <u>*1349*</u> |
| **111** | 2 | SAUSA300_1348 | ***papS*** |
| **111** | 3 | SAUSA300_1347 | ***birA*** |
| **111** | 4 | SAUSA300_1346 | <u>*1346*</u> |
| **112** | 1 | SAUSA300_1354 | <u>*1354*</u> |
| **112** | 2 | SAUSA300_1353 | <u>*1353*</u> |

| Operon Number | Gene position in operon | Locus Tag | Gene Name |
|---|---|---|---|
| **112** | 3 | SAUSA300_1352 | *1352* |
| **112** | 4 | SAUSA300_1351 | ***1351*** |
| **113** | 1 | SAUSA300_1360 | ***ubiE*** |
| **113** | 2 | SAUSA300_1359 | ***1359*** |
| **114** | 1 | SAUSA300_1362 | ***hup*** |
| **115** | 1 | SAUSA300_1364 | ***engA*** |
| **115** | 2 | SAUSA300_1363 | *gpsA* |
| **116** | 1 | SAUSA300_1367 | ***cmk*** |
| **116** | 2 | SAUSA300_1366 | *1366* |
| **117** | 1 | SAUSA300_1373 | ***fer*** |
| **118** | 1 | SAUSA300_1453 | ***rnz*** |
| **119** | 1 | SAUSA300_1454 | ***zwf*** |
| **120** | 1 | SAUSA300_1460 | *1460* |
| **120** | 2 | SAUSA300_1459 | ***gnd*** |
| **121** | 1 | SAUSA300_1467 | *ipdA* |
| **121** | 2 | SAUSA300_1466 | ***bfmBAA*** |
| **121** | 3 | SAUSA300_1465 | *1465* |
| **121** | 4 | SAUSA300_1464 | ***bmfBB*** |
| **122** | 1 | SAUSA300_1476 | ***accB*** |
| **122** | 2 | SAUSA300_1475 | ***accC*** |
| **122** | 3 | SAUSA300_1474 | *1474* |
| **122** | 4 | SAUSA300_1473 | *nusB* |
| **122** | 5 | SAUSA300_1472 | *xseA* |
| **122** | 6 | SAUSA300_1471 | *xseB* |
| **122** | 7 | SAUSA300_1470 | *1470* |
| **123** | 1 | SAUSA300_1491 | *pepQ2* |
| **123** | 2 | SAUSA300_1490 | ***efp*** |
| **124** | 1 | SAUSA300_1492 | ***1492*** |
| **124** | 2 | SAUSA300_1493 | *1493* |
| **125** | 1 | SAUSA300_1511 | ***rpmG*** |
| **126** | 1 | SAUSA300_1520 | ***trmK*** |
| **126** | 2 | SAUSA300_1519 | *1519* |
| **127** | 1 | SAUSA300_1521 | ***rpoD*** |
| **128** | 1 | SAUSA300_1522 | ***dnaG*** |
| **129** | 1 | SAUSA300_1525 | ***glyS*** |
| **130** | 1 | SAUSA300_1531 | *phoH* |
| **130** | 2 | SAUSA300_1530 | ***ybeY*** |
| **130** | 3 | SAUSA300_1529 | *dgcA* |
| **130** | 4 | SAUSA300_1528 | *cdd* |
| **130** | 5 | SAUSA300_1527 | *era* |
| **130** | 6 | SAUSA300_1526 | *recO* |
| **131** | 1 | SAUSA300_1535 | ***rpsU*** |
| **132** | 1 | SAUSA300_1542 | *hrcA* |
| **132** | 2 | SAUSA300_1541 | ***grpE*** |
| **132** | 3 | SAUSA300_1540 | ***dnaK*** |

| **Operon Number** | **Gene position in operon** | **Locus Tag** | **Gene Name** |
|---|---|---|---|
| **132** | 4 | SAUSA300_1539 | *dnaJ* |
| **132** | 5 | SAUSA300_1538 | *prmA* |
| **132** | 6 | SAUSA300_1537 | *rsmE* |
| **132** | 7 | SAUSA300_1536 | *mtaB* |
| **133** | 1 | SAUSA300_1545 | ***rpsT*** |
| **134** | 1 | SAUSA300_1548 | *comEB* |
| **134** | 2 | SAUSA300_1547 | *comEC* |
| **134** | 3 | SAUSA300_1546 | ***holA*** |
| **135** | 1 | SAUSA300_1558 | ***mtnN*** |
| **135** | 2 | SAUSA300_1557 | ***1557*** |
| **135** | 3 | SAUSA300_1556 | *1556* |
| **135** | 4 | SAUSA300_1555 | *aroE* |
| **135** | 5 | SAUSA300_1554 | *1554* |
| **135** | 6 | SAUSA300_1553 | *nadD* |
| **135** | 7 | SAUSA300_1552 | *1552* |
| **135** | 8 | SAUSA300_1551 | *1551* |
| **135** | 9 | SAUSA300_1550 | *1550* |
| **135** | 10 | SAUSA300_1549 | *1549* |
| **136** | 1 | SAUSA300_1571 | *1571* |
| **136** | 2 | SAUSA300_1570 | *1570* |
| **136** | 3 | SAUSA300_1569 | *1569* |
| **136** | 4 | SAUSA300_1568 | *udk* |
| **136** | 5 | SAUSA300_1567 | ***greA*** |
| **137** | 1 | SAUSA300_1575 | ***alaS*** |
| **137** | 2 | SAUSA300_1574 | *1574* |
| **137** | 3 | SAUSA300_1573 | *1573* |
| **137** | 4 | SAUSA300_1572 | *1572* |
| **138** | 1 | SAUSA300_1579 | ***iscS*** |
| **138** | 2 | SAUSA300_1578 | *mnmA* |
| **139** | 1 | SAUSA300_1587 | ***hisS*** |
| **139** | 2 | SAUSA300_1586 | *aspS* |
| **140** | 1 | SAUSA300_1590 | ***relA*** |
| **140** | 2 | SAUSA300_1589 | ***dtd*** |
| **140** | 3 | SAUSA300_1588 | *lytH* |
| **141** | 1 | SAUSA300_1593 | ***secF*** |
| **142** | 1 | SAUSA300_1600 | ***obgE*** |
| **142** | 2 | SAUSA300_1599 | *1599* |
| **142** | 3 | SAUSA300_1598 | *ruvA* |
| **142** | 4 | SAUSA300_1597 | *ruvB* |
| **142** | 5 | SAUSA300_1596 | *queA* |
| **142** | 6 | SAUSA300_1595 | *tgt* |
| **142** | 7 | SAUSA300_1594 | *yajC* |
| **143** | 1 | SAUSA300_1603 | ***rplU*** |
| **143** | 2 | SAUSA300_1602 | ***1602*** |

| Operon Number | Gene position in operon | Locus Tag | Gene Name |
|---|---|---|---|
| **143** | 3 | SAUSA300_1601 | ***rpmA*** |
| **144** | 1 | SAUSA300_1611 | ***valS*** |
| **144** | 2 | SAUSA300_1610 | *folC* |
| **145** | 1 | SAUSA300_1619 | ***<u>hemA</u>*** |
| **145** | 2 | SAUSA300_1618 | *<u>hemX</u>* |
| **145** | 3 | SAUSA300_1617 | *hemC* |
| **145** | 4 | SAUSA300_1616 | *hemD* |
| **145** | 5 | SAUSA300_1615 | *<u>hemB</u>* |
| **145** | 6 | SAUSA300_1614 | *<u>hemL</u>* |
| **146** | 1 | SAUSA300_1620 | ***engB*** |
| **147** | 1 | SAUSA300_1624 | ***1624*** |
| **147** | 2 | SAUSA300_1623 | *<u>1623</u>* |
| **148** | 1 | SAUSA300_1627 | ***infC*** |
| **148** | 2 | SAUSA300_1626 | *rpmI* |
| **148** | 3 | SAUSA300_1625 | ***rplT*** |
| **149** | 1 | SAUSA300_1629 | ***thrS*** |
| **150** | 1 | SAUSA300_1632 | ***nrdR*** |
| **150** | 2 | SAUSA300_1631 | ***dnaB*** |
| **150** | 3 | SAUSA300_1630 | *dnaI* |
| **151** | 1 | SAUSA300_1636 | *<u>polA</u>* |
| **151** | 2 | SAUSA300_1635 | ***mutM*** |
| **151** | 3 | SAUSA300_1634 | ***coaE*** |
| **152** | 1 | SAUSA300_1645 | ***pfkA*** |
| **152** | 2 | SAUSA300_1644 | *pykA* |
| **153** | 1 | SAUSA300_1647 | ***accD*** |
| **153** | 2 | SAUSA300_1646 | *accA* |
| **154** | 1 | SAUSA300_1650 | *<u>1650</u>* |
| **154** | 2 | SAUSA300_1649 | ***dnaE*** |
| **155** | 1 | SAUSA300_1658 | *<u>1658</u>* |
| **155** | 2 | SAUSA300_1657 | ***ackA*** |
| **156** | 1 | SAUSA300_1666 | ***rpsD*** |
| **157** | 1 | SAUSA300_1673 | ***plsC*** |
| **158** | 1 | SAUSA300_1675 | ***tyrS*** |
| **159** | 1 | SAUSA300_1691 | *<u>1691</u>* |
| **159** | 2 | SAUSA300_1690 | ***1690*** |
| **159** | 3 | SAUSA300_1689 | *1689* |
| **159** | 4 | SAUSA300_1688 | ***1688*** |
| **159** | 5 | SAUSA300_1687 | *1687* |
| **159** | 6 | SAUSA300_1686 | ***murC*** |
| **160** | 1 | SAUSA300_1700 | ***murJ*** |
| **160** | 2 | SAUSA300_1699 | *<u>1699</u>* |
| **160** | 3 | SAUSA300_1698 | *<u>1698</u>* |
| **161** | 1 | SAUSA300_1704 | ***leuS*** |
| **161** | 2 | SAUSA300_1703 | *1703* |
| **162** | 1 | SAUSA300_1729 | ***1729*** |

| Operon Number | Gene position in operon | Locus Tag | Gene Name |
|---|---|---|---|
| **163** | 1 | SAUSA300_1730 | ***metK*** |
| **164** | 1 | SAUSA300_1748 | ***1748*** |
| **165** | 1 | SAUSA300_1749 | ***1749*** |
| **166** | 1 | SAUSA300_1793 | <u>*1793*</u> |
| **166** | 2 | SAUSA300_1792 | <u>*1792*</u> |
| **166** | 3 | SAUSA300_1791 | ***cbf1*** |
| **167** | 1 | SAUSA300_1869 | ***map*** |
| **168** | 1 | SAUSA300_1873 | ***murT*** |
| **168** | 2 | SAUSA300_1872 | *gatD* |
| **169** | 1 | SAUSA300_1879 | ***dgkB*** |
| **169** | 2 | SAUSA300_1878 | <u>*rumA*</u> |
| **170** | 1 | SAUSA300_1882 | ***gatC*** |
| **170** | 2 | SAUSA300_1881 | ***gatA*** |
| **170** | 3 | SAUSA300_1880 | *gatB* |
| **171** | 1 | SAUSA300_1887 | <u>*pcrB*</u> |
| **171** | 2 | SAUSA300_1886 | ***pcrA*** |
| **171** | 3 | SAUSA300_1885 | ***ligA*** |
| **171** | 4 | SAUSA300_1884 | <u>*1884*</u> |
| **172** | 1 | SAUSA300_1894 | ***pncB*** |
| **172** | 2 | SAUSA300_1893 | *nadE* |
| **173** | 1 | SAUSA300_1899 | <u>***pncA***</u> |
| **173** | 2 | SAUSA300_1900 | ***ppaC*** |
| **174** | 1 | SAUSA300_1914 | ***pmtR*** |
| **174** | 2 | SAUSA300_1913 | ***pmtA*** |
| **174** | 3 | SAUSA300_1912 | <u>*pmtB*</u> |
| **174** | 4 | SAUSA300_1911 | <u>***pmtC***</u> |
| **174** | 5 | SAUSA300_1910 | *pmtD* |
| **175** | 1 | SAUSA300_1983 | ***groES*** |
| **175** | 2 | SAUSA300_1982 | *groEL* |
| **176** | 1 | SAUSA300_1988 | ***hld*** |
| **177** | 1 | SAUSA300_2005 | ***tsaE*** |
| **177** | 2 | SAUSA300_2004 | *tsaB* |
| **177** | 3 | SAUSA300_2003 | <u>*rimI*</u> |
| **177** | 4 | SAUSA300_2002 | ***gcp*** |
| **178** | 1 | SAUSA300_2031 | <u>*2031*</u> |
| **178** | 2 | SAUSA300_2030 | <u>*2030*</u> |
| **178** | 3 | SAUSA300_2029 | ***2029*** |
| **178** | 4 | SAUSA300_2028 | ***acpS*** |
| **178** | 5 | SAUSA300_2027 | *alr* |
| **179** | 1 | SAUSA300_2039 | ***ddl*** |
| **179** | 2 | SAUSA300_2038 | *murF* |
| **180** | 1 | SAUSA300_2046 | ***oxaA*** |
| **181** | 1 | SAUSA300_2056 | <u>*2056*</u> |
| **181** | 2 | SAUSA300_2055 | <u>***murA***</u> |
| **181** | 3 | SAUSA300_2054 | *fabZ* |

| Operon Number | Gene position in operon | Locus Tag | Gene Name |
|---|---|---|---|
| **182** | 1 | SAUSA300_2062 | ***atpF*** |
| **182** | 2 | SAUSA300_2061 | *atpH* |
| **182** | 3 | SAUSA300_2060 | *atpA* |
| **182** | 4 | SAUSA300_2059 | *atpG* |
| **182** | 5 | SAUSA300_2058 | *atpD* |
| **182** | 6 | SAUSA300_2057 | *atpC* |
| **183** | 1 | SAUSA300_2073 | *tdk* |
| **183** | 2 | SAUSA300_2072 | ***prfA*** |
| **183** | 3 | SAUSA300_2071 | *prmC* |
| **183** | 4 | SAUSA300_2070 | *2070* |
| **183** | 5 | SAUSA300_2069 | *ptpB* |
| **184** | 1 | SAUSA300_2074 | ***rpmE2*** |
| **185** | 1 | SAUSA300_2079 | ***fbaA*** |
| **186** | 1 | SAUSA300_2081 | ***pyrG*** |
| **187** | 1 | SAUSA300_2084 | ***coaA*** |
| **188** | 1 | SAUSA300_2104 | ***glmS*** |
| **189** | 1 | SAUSA300_2113 | ***dacA*** |
| **189** | 2 | SAUSA300_2112 | *ybbR* |
| **189** | 3 | SAUSA300_2111 | *glmM* |
| **190** | 1 | SAUSA300_2172 | ***rplM*** |
| **190** | 2 | SAUSA300_2171 | ***rpsI*** |
| **191** | 1 | SAUSA300_2182 | ***infA*** |
| **191** | 2 | SAUSA300_2181 | *rpmJ* |
| **191** | 3 | SAUSA300_2180 | *rpsM* |
| **191** | 4 | SAUSA300_2179 | *rpsK* |
| **191** | 5 | SAUSA300_2178 | *rpoA* |
| **191** | 6 | SAUSA300_2177 | ***rplQ*** |
| **192** | 1 | SAUSA300_2205 | ***rpsJ*** |
| **192** | 2 | SAUSA300_2204 | *rplC* |
| **192** | 3 | SAUSA300_2203 | *rplD* |
| **192** | 4 | SAUSA300_2202 | *rplW* |
| **192** | 5 | SAUSA300_2201 | *rplB* |
| **192** | 6 | SAUSA300_2200 | *rpsS* |
| **192** | 7 | SAUSA300_2199 | *rplV* |
| **192** | 8 | SAUSA300_2198 | *rpsC* |
| **192** | 9 | SAUSA300_2197 | *rplP* |
| **192** | 10 | SAUSA300_2196 | *rpmC* |
| **192** | 11 | SAUSA300_2195 | *rpsQ* |
| **192** | 12 | SAUSA300_2194 | *rplN* |
| **192** | 13 | SAUSA300_2193 | *rplX* |
| **192** | 14 | SAUSA300_2192 | *rplE* |
| **192** | 15 | SAUSA300_2191 | *rpsN* |
| **192** | 16 | SAUSA300_2190 | *rpsH* |
| **192** | 17 | SAUSA300_2189 | *rplF* |
| **192** | 18 | SAUSA300_2188 | *rplR* |

| Operon Number | Gene position in operon | Locus Tag | Gene Name |
|---|---|---|---|
| **192** | 19 | SAUSA300_2187 | *rpsE* |
| **192** | 20 | SAUSA300_2186 | *rpmD* |
| **192** | 21 | SAUSA300_2185 | ***rplO*** |
| **192** | 22 | SAUSA300_2184 | ***secY*** |
| **192** | 23 | SAUSA300_2183 | ***adK*** |
| **193** | 1 | SAUSA300_2214 | ***femX*** |
| **193** | 2 | SAUSA300_2213 | <u>*2213*</u> |
| **194** | 1 | SAUSA300_2238 | ***ureA*** |
| **194** | 2 | SAUSA300_2239 | <u>*ureB*</u> |
| **194** | 3 | SAUSA300_2240 | <u>*ureC*</u> |
| **194** | 4 | SAUSA300_2241 | *ureE* |
| **194** | 5 | SAUSA300_2242 | <u>*ureF*</u> |
| **194** | 6 | SAUSA300_2243 | <u>*ureG*</u> |
| **194** | 7 | SAUSA300_2244 | <u>*ureD*</u> |
| **195** | 1 | SAUSA300_2283 | ***rpiA*** |
| **196** | 1 | SAUSA300_2293 | <u>*corA*</u> |
| **196** | 2 | SAUSA300_2292 | ***fni*** |
| **197** | 1 | SAUSA300_2483 | ***mvaA*** |
| **198** | 1 | SAUSA300_2484 | ***mvaS*** |
| **199** | 1 | SAUSA300_2646 | <u>***trmE***</u> |
| **199** | 2 | SAUSA300_2645 | <u>*gidA*</u> |
| **199** | 3 | SAUSA300_2644 | <u>*gidB*</u> |
| **199** | 4 | SAUSA300_2643 | <u>*2643*</u> |
| **200** | 1 | SAUSA300_2647 | ***rnpA*** |
| **200** | 2 | SAUSA300_2648 | ***rpmH*** |

Genes listed in bold were targeted with specific sgRNAs in the LCML library. Those appearing in the Nebraska transposon mutant library are underlined.

**Supplementary Table 2. Lisbon CRISPRi Mutant Library primer list**

| LCML N° | Plasmid | Primer N° | Primer Sequence |
|---|---|---|---|
| | | 5846 - EcR (reverse) | ACTAGTATTATACCTAGGACTGAGCTAGC |
| 1 | psg0001_dnaA | 8276 | ATACACAAAAATAACAGCCGgttttAGAGCTAGAAATAGCAAGTTAAAATAAGGC |
| 2 | psg0002_dnaN | 8277 | TCGTTTATATATAATTATATgttttAGAGCTAGAAATAGCAAGTTAAAATAAGGC |
| 3 | psg0003 | 8278 | ACAAATTTCATTTAAAATAGgttttAGAGCTAGAAATAGCAAGTTAAAATAAGGC |
| 4 | psg0009_serS | 8279 | CAAATAATTATCATTTATTAgttttAGAGCTAGAAATAGCAAGTTAAAATAAGGC |
| 5 | psg0016_dnaB | 8280 | TCATACATTCTATCCATGAAgttttAGAGCTAGAAATAGCAAGTTAAAATAAGGC |
| 6 | psg0020_walR | 8281 | ATGTTTGTGTAAAAAATCACgttttAGAGCTAGAAATAGCAAGTTAAAATAAGGC |
| 7 | psg0024_walJ | 8282 | TGTAACTTTAGTTCATCGACgttttAGAGCTAGAAATAGCAAGTTAAAATAAGGC |
| 8 | psg0089 | 8283 | GCCAAAATAAAAAATGGACGgttttAGAGCTAGAAATAGCAAGTTAAAATAAGGC |
| 9 | psg0248_tarF | 8284 | ATATAATTACAAAAACACGTgttttAGAGCTAGAAATAGCAAGTTAAAATAAGGC |
| 10 | psg0249_ispD | 8285 | ATATAGATATAGTTGAATGGgttttAGAGCTAGAAATAGCAAGTTAAAATAAGGC |
| 11 | psg0363 | 8286 | TGATTATAAGCAGTCATAATgttttAGAGCTAGAAATAGCAAGTTAAAATAAGGC |
| 12 | psg0367_ssb | 8287 | TTTATATTTGCACCTCCTTGgttttAGAGCTAGAAATAGCAAGTTAAAATAAGGC |
| 13 | psg0388_guaB | 8288 | GGTAAAATATCAGATTGTGCgttttAGAGCTAGAAATAGCAAGTTAAAATAAGGC |
| 14 | psg0452_dnaX | 8289 | GGAAATTTTACGATTCCGTGgttttAGAGCTAGAAATAGCAAGTTAAAATAAGGC |
| 15 | psg0453 | 8290 | ATTGATTTTCAAGGAGGAAACgttttAGAGCTAGAAATAGCAAGTTAAAATAAGGC |
| 16 | psg00459_tmk | 8291 | GTTTCTCCTTTGAAAATAATgttttAGAGCTAGAAATAGCAAGTTAAAATAAGGC |
| 17 | psg0461_holB | 8292 | CCTTTTGCTTTTATATAAAAgttttAGAGCTAGAAATAGCAAGTTAAAATAAGGC |
| 18 | psg0467_metS | 8303 | AAGATTTCTATGCATTTCAAgttttAGAGCTAGAAATAGCAAGTTAAAATAAGGC |
| 19 | psg0477_glmU | 8304 | TTCAGCACCATGTCCTACGAgttttAGAGCTAGAAATAGCAAGTTAAAATAAGGC |
| 20 | psg0478_prs | 8305 | AGATTACATTAATATTACATgttttAGAGCTAGAAATAGCAAGTTAAAATAAGGC |
| 21 | psg0480_pth | 8306 | GACAAATTCATCATTGTCATgttttAGAGCTAGAAATAGCAAGTTAAAATAAGGC |
| 22 | psg0484 | 8398 | TTCTAGCCACTGCTTTAACGgttttAGAGCTAGAAATAGCAAGTTAAAATAAGGC |
| 23 | psg0485_divIC | 8399 | CAATATCATTGCGATGTTTTgttttAGAGCTAGAAATAGCAAGTTAAAATAAGGC |
| 24 | psg0487_tilS | 8400 | ACTTTTACAATTTTCAAAAAgttttAGAGCTAGAAATAGCAAGTTAAAATAAGGC |
| 25 | psg0492_folP | 8494 | TTTATTATCTTATCATTCACgttttAGAGCTAGAAATAGCAAGTTAAAATAAGGC |
| 26 | psg0496_lysS | 8495 | TAATCCTTTAGTATTCCAACgttttAGAGCTAGAAATAGCAAGTTAAAATAAGGC |
| 27 | psg0513_gltX | 8496 | TGAAGATACCCAGTTGGACTgttttAGAGCTAGAAATAGCAAGTTAAAATAAGGC |
| 28 | psg0514_cysE | 8497 | CATACAAGTTGTATTGTTAGgttttAGAGCTAGAAATAGCAAGTTAAAATAAGGC |
| 29 | psg0532_fusA | 8498 | CTGAATAAATACGATAGATAgttttAGAGCTAGAAATAGCAAGTTAAAATAAGGC |
| 30 | psg0533 _tuf | 8499 | GTGACCGATAGTACCGATATgttttAGAGCTAGAAATAGCAAGTTAAAATAAGGC |
| 31 | psg0551_folE2 | 8500 | TTAAAGTGATATGTCCAATAgttttAGAGCTAGAAATAGCAAGTTAAAATAAGGC |

| | | | |
|---|---|---|---|
| 32 | psg0570_eutD | 8501 | TAAATCTTATTAATCATTCAgttttAGAGCTAGAAATAGCAAGTTAAAATAAGGC |
| 33 | psg0572_mvaK1 | 8502 | AAAAATAATGAATCAAGTATgttttAGAGCTAGAAATAGCAAGTTAAAATAAGGC |
| 34 | psg0596_argS | 8503 | ATATCAAGATAATGAAAAAAgttttAGAGCTAGAAATAGCAAGTTAAAATAAGGC |
| 35 | psg0623_tagA | 8504 | CTTTCTTCAACAGTCATAACgttttAGAGCTAGAAATAGCAAGTTAAAATAAGGC |
| 36 | psg0624_tagH | 8505 | AATCTAGATAAATGTGAATAgttttAGAGCTAGAAATAGCAAGTTAAAATAAGGC |
| 37 | psg0625_tagG | 8674 | AAACCATAATTTGCATAACAgttttAGAGCTAGAAATAGCAAGTTAAAATAAGGC |
| 38 | psg0626_tagB | 8675 | TGCGGTTTATCAATCACTTGgttttAGAGCTAGAAATAGCAAGTTAAAATAAGGC |
| 39 | psg0628_tagD | 8676 | TCCTCATATTGTCACATCATgttttAGAGCTAGAAATAGCAAGTTAAAATAAGGC |
| 40 | psg0703_ltaS | 8677 | CTTCAAGGTAATCGTTATTAgttttAGAGCTAGAAATAGCAAGTTAAAATAAGGC |
| 41 | psg0715_nrdI | 8678 | TAAAAAATGCTTTAACAACAgttttAGAGCTAGAAATAGCAAGTTAAAATAAGGC |
| 42 | psg0722_murB | 8679 | TCAACTTTAATTTTTTCATTgttttAGAGCTAGAAATAGCAAGTTAAAATAAGGC |
| 43 | psg0731_tagO | 8680 | TTCGATATTGCAATAACAATgttttAGAGCTAGAAATAGCAAGTTAAAATAAGGC |
| 44 | psg0737_secA | 8681 | CCTTTAGCTAAAAAACTGTTgttttAGAGCTAGAAATAGCAAGTTAAAATAAGGC |
| 45 | psg0738_prfB | 8682 | GTTTGGTTATCCCAAAAATTgttttAGAGCTAGAAATAGCAAGTTAAAATAAGGC |
| 46 | psg0743_hprK | 8683 | TATTGCTTGCTTAATTTACAgttttAGAGCTAGAAATAGCAAGTTAAAATAAGGC |
| 47 | psg0749 | 8684 | TTGTCCACCCGTACAACCGAgttttAGAGCTAGAAATAGCAAGTTAAAATAAGGC |
| 48 | psg0756_gapA | 8685 | TTCAAGTATTATCTTTGCTGgttttAGAGCTAGAAATAGCAAGTTAAAATAAGGC |
| 49 | psg0761 | 8755 | CGTTAAATTAAGTAATGCTTgttttAGAGCTAGAAATAGCAAGTTAAAATAAGGC |
| 50 | psg0765_smpB | 8756 | CGATTTTCCGCTAATGTACCgttttAGAGCTAGAAATAGCAAGTTAAAATAAGGC |
| 51 | psg0818_sufC | 8757 | TATCCTCAATAGACACATGTgttttAGAGCTAGAAATAGCAAGTTAAAATAAGGC |
| 52 | psg0835_dltA | 8758 | TGTCTAACAGCAATGCTTTGgttttAGAGCTAGAAATAGCAAGTTAAAATAAGGC |
| 53 | psg0858 | 8759 | GTCTCAACAAACGCACCGTAgttttAGAGCTAGAAATAGCAAGTTAAAATAAGGC |
| 54 | psg0865_pgi | 8760 | TTTCAACAAACATTTCAAACgttttAGAGCTAGAAATAGCAAGTTAAAATAAGGC |
| 55 | psg0868_spsB | 8761 | CGCTCGCCATCTTTCAAAGTgttttAGAGCTAGAAATAGCAAGTTAAAATAAGGC |
| 56 | psg0885_fabH | 8762 | GCATTGTCAATAATCTTTTCgttttAGAGCTAGAAATAGCAAGTTAAAATAAGGC |
| 57 | psg0897_trpS | 8763 | TAGTAGGAATTCCACTAGGTgttttAGAGCTAGAAATAGCAAGTTAAAATAAGGC |
| 58 | psg0898_spxA | 8764 | TTACGGCAAGATGTGCAACTgttttAGAGCTAGAAATAGCAAGTTAAAATAAGGC |
| 59 | psg0908_ppnK | 8765 | GTTCATCATTTTATGCTTTAgttttAGAGCTAGAAATAGCAAGTTAAAATAAGGC |
| 60 | psg0912_fabI | 8766 | CCTTATAATAATTAATTTAAgttttAGAGCTAGAAATAGCAAGTTAAAATAAGGC |
| 61 | psg0919_murE | 9735 | ACTATTAAAAATAAAACATAgttttAGAGCTAGAAATAGCAAGTTAAAATAAGGC |
| 62 | psg0922 | 9220 | ACGACAAGTACCCATAAATAgttttAGAGCTAGAAATAGCAAGTTAAAATAAGGC |
| 63 | psg0944_menA | 9736 | ACGGAAGCAGTTAATGTATGgttttAGAGCTAGAAATAGCAAGTTAAAATAAGGC |
| 64 | psg0948_menB | 9737 | GTAAACGCATTGCGTACTTCgttttAGAGCTAGAAATAGCAAGTTAAAATAAGGC |

| | | | |
|---|---|---|---|
| 65 | psg0983_ptsH | 9221 | GTTTGTACTAACATTGTTGCgttttAGAGCTAGAAATAGCAAGTTAAAATAAGGC |
| 66 | psg0989_rnjA | 9286 | GCATATACACCTACTTCATTgttttAGAGCTAGAAATAGCAAGTTAAAATAAGGC |
| 67 | psg0990_rpoY | 9287 | AAACTTTAAATACTGCCATAgttttAGAGCTAGAAATAGCAAGTTAAAATAAGGC |
| 68 | psg0991_def | 9288 | GCTGCTTTTTGACGCAAAGTgttttAGAGCTAGAAATAGCAAGTTAAAATAAGGC |
| 69 | psg1013_ftsW | 9738 | CGGATAATCAATAAACTTTGgttttAGAGCTAGAAATAGCAAGTTAAAATAAGGC |
| 70 | psg1024_coaD | 9289 | GTAATGGGGTCAAAACTACCgttttAGAGCTAGAAATAGCAAGTTAAAATAAGGC |
| 71 | psg1026 | 9739 | ACCGTTTGATCAAATTCAAAgttttAGAGCTAGAAATAGCAAGTTAAAATAAGGC |
| 72 | psg1037_pheS | 9740 | AACGCAGGTTTATCTTCATTgttttAGAGCTAGAAATAGCAAGTTAAAATAAGGC |
| 73 | psg1044_trxA | 8877 | ACCGGAGCGATCATTTTACAgttttAGAGCTAGAAATAGCAAGTTAAAATAAGGC |
| 74 | psg1049_murI | 8878 | CCAGAGTCTATTACACCTATgttttAGAGCTAGAAATAGCAAGTTAAAATAAGGC |
| 75 | psg1074_ftsL | 8879 | AAACTTGTTCGTCATATGGTgttttAGAGCTAGAAATAGCAAGTTAAAATAAGGC |
| 76 | psg1075_pbp1_b | 8880 | GTCCGAATAAACCAACAAGTgttttAGAGCTAGAAATAGCAAGTTAAAATAAGGC |
| 77 | psg1076_mraY | 8881 | TTTAATGTAGGTATTAAAACgttttAGAGCTAGAAATAGCAAGTTAAAATAAGGC |
| 78 | psg1077_murD | 8882 | AATAAAAATGTATTAGTTGTgttttAGAGCTAGAAATAGCAAGTTAAAATAAGGC |
| 79 | psg1078_divIB | 8883 | GGAATACATCTAAGAAAAGAgttttAGAGCTAGAAATAGCAAGTTAAAATAAGGC |
| 80 | psg1079_ftsA | 8884 | ATTTTTTATACCGCTCGTGTgttttAGAGCTAGAAATAGCAAGTTAAAATAAGGC |
| 81 | psg1082 | 8885 | TTTGTAACTGCAATCACGTTgttttAGAGCTAGAAATAGCAAGTTAAAATAAGGC |
| 82 | psg1083_sepF | 8886 | TTTACCTGTTGTTGTTTGTCgttttAGAGCTAGAAATAGCAAGTTAAAATAAGGC |
| 83 | psg1087_ileS | 8887 | CGCATTGGGAAATCTGTTTTgttttAGAGCTAGAAATAGCAAGTTAAAATAAGGC |
| 84 | psg1102_gmk | 8888 | GTACCTTTACCTACTCCAGAgttttAGAGCTAGAAATAGCAAGTTAAAATAAGGC |
| 85 | psg1104_coaBC | 8889 | TGCCGCAATGCCACCTGTAAgttttAGAGCTAGAAATAGCAAGTTAAAATAAGGC |
| 86 | psg1105_priA | 8890 | GATGACAGATTCGAGTTGTTgttttAGAGCTAGAAATAGCAAGTTAAAATAAGGC |
| 87 | psg1109_fmt | 8891 | AAAACAGTTGTTGAAAAGTCgttttAGAGCTAGAAATAGCAAGTTAAAATAAGGC |
| 88 | psg1115_rpe | 8892 | AAATCAACAGATAATAATGAgttttAGAGCTAGAAATAGCAAGTTAAAATAAGGC |
| 89 | psg1122_plsX | 8893 | GATATCGTATTAGAAGCCGTgttttAGAGCTAGAAATAGCAAGTTAAAATAAGGC |
| 90 | psg1125_acpP | 8894 | TATCAGCGTCTACACCTAAAgttttAGAGCTAGAAATAGCAAGTTAAAATAAGGC |
| 91 | psg1128_ftsY | 8895 | TTGACCTTGTTCTTCTGTTAgttttAGAGCTAGAAATAGCAAGTTAAAATAAGGC |
| 92 | psg1133_trmD | 8896 | AAAACACCATCAAACATTTCgttttAGAGCTAGAAATAGCAAGTTAAAATAAGGC |
| 93 | psg1136_rbgA | 8897 | TTGGCTTTCGCCATATGTCCgttttAGAGCTAGAAATAGCAAGTTAAAATAAGGC |
| 94 | psg1143_topA | 8898 | TCAATGGTTTTTGCTTTTGCgttttAGAGCTAGAAATAGCAAGTTAAAATAAGGC |
| 95 | psg1150_tsf | 8899 | TAGCAATACCTTTTTCACGTgttttAGAGCTAGAAATAGCAAGTTAAAATAAGGC |
| 96 | psg1151_pyrH | 8900 | GCAACACTTTTAATAATTACgttttAGAGCTAGAAATAGCAAGTTAAAATAAGGC |
| 97 | psg1152_frr | 8981 | AATTAGCTAACATCAGTGCAgttttAGAGCTAGAAATAGCAAGTTAAAATAAGGC |

| | | | |
|---|---|---|---|
| 98 | psg1153_uppS | 8982 | ATTTATTAGCTTTTTAAACAgttttAGAGCTAGAAATAGCAAGTTAAAATAAGGC |
| 99 | psg1156_proS | 8983 | CATCGTTGGTATAAAAACTTgttttAGAGCTAGAAATAGCAAGTTAAAATAAGGC |
| 100 | psg1157_polC | 8984 | TCTTCATGAGCTAAGAATTGgttttAGAGCTAGAAATAGCAAGTTAAAATAAGGC |
| 101 | psg1159_nusA | 8985 | GCATCAATTAATACTGCTCTgttttAGAGCTAGAAATAGCAAGTTAAAATAAGGC |
| 102 | psg1168_rnjB | 8986 | TGAATAGTGAGTTTATATATgttttAGAGCTAGAAATAGCAAGTTAAAATAAGGC |
| 103 | psg1176_pgsA | 8987 | CTAAAAACCGTAATCTGGTTgttttAGAGCTAGAAATAGCAAGTTAAAATAAGGC |
| 104 | psg1200_glnR | 8988 | TGATGCAATCAGACGAAATAgttttAGAGCTAGAAATAGCAAGTTAAAATAAGGC |
| 105 | psg1237_lexA | 8989 | CCAATTTCGCGAACACTAGGgttttAGAGCTAGAAATAGCAAGTTAAAATAAGGC |
| 106 | psg1239_tkt | 8990 | TTGAAGTAATCTTTAGATTGgttttAGAGCTAGAAATAGCAAGTTAAAATAAGGC |
| 107 | psg1249_plsY | 8991 | TTTCCAATTACGAATCCACTgttttAGAGCTAGAAATAGCAAGTTAAAATAAGGC |
| 108 | psg1250_parE | 8992 | GTTGATCCAATATACATACCgttttAGAGCTAGAAATAGCAAGTTAAAATAAGGC |
| 109 | psg1257_msrR | 9018 | TTCTTCTTCTTTTTTCGCTTgttttAGAGCTAGAAATAGCAAGTTAAAATAAGGC |
| 110 | psg1269_femA | 9019 | TTACAGATAGCATGCCATACgttttAGAGCTAGAAATAGCAAGTTAAAATAAGGC |
| 111 | psg1270_femB | 9020 | TTGTACAAAGTTGTCAAATTgttttAGAGCTAGAAATAGCAAGTTAAAATAAGGC |
| 112 | psg1311_murG | 9021 | TGTCCAACTGTTCCCCCTCCgttttAGAGCTAGAAATAGCAAGTTAAAATAAGGC |
| 113 | psg1319_folA | 9022 | CAAGTCATGTGCAACTAGAAgttttAGAGCTAGAAATAGCAAGTTAAAATAAGGC |
| 114 | psg1320_thyA | 9023 | ACTTTCTTTGTCGTTAATAGgttttAGAGCTAGAAATAGCAAGTTAAAATAAGGC |
| 115 | psg1340_recU | 9024 | TTACGATATGGTTTACCATTgttttAGAGCTAGAAATAGCAAGTTAAAATAAGGC |
| 116 | psg1341_pbp2 | 9025 | CCATTATTACCGTTTTTCTTgttttAGAGCTAGAAATAGCAAGTTAAAATAAGGC |
| 117 | psg1344_dnaD | 9026 | AATTCTCTTCGTATCACTACgttttAGAGCTAGAAATAGCAAGTTAAAATAAGGC |
| 118 | psg1347_birA | 9027 | CTTTGTCCAGATATATAATTgttttAGAGCTAGAAATAGCAAGTTAAAATAAGGC |
| 119 | psg1348_papS | 9028 | TCTTGAATTTGTTCTAATATgttttAGAGCTAGAAATAGCAAGTTAAAATAAGGC |
| 120 | psg1351 | 9029 | ACTTTGTGTTGTGCCCATAAgttttAGAGCTAGAAATAGCAAGTTAAAATAAGGC |
| 121 | psg1360_ubiE | 9030 | AAAACGCGTCATGAAAGACAgttttAGAGCTAGAAATAGCAAGTTAAAATAAGGC |
| 122 | psg1362_hup | 9031 | CATTAGACATTCACCTCCTGgttttAGAGCTAGAAATAGCAAGTTAAAATAAGGC |
| 123 | psg1364_engA | 9032 | ATTGTAGATTTACCTACATTgttttAGAGCTAGAAATAGCAAGTTAAAATAAGGC |
| 124 | psg1367_cmk | 9033 | ACCATCTAATGCAATATTAAgttttAGAGCTAGAAATAGCAAGTTAAAATAAGGC |
| 125 | psg1373_fer | 9034 | TCGTCGTAATCATATATATCgttttAGAGCTAGAAATAGCAAGTTAAAATAAGGC |
| 126 | psg1453_rnz | 9035 | TGTGTATTTCTCTCTTTTGTgttttAGAGCTAGAAATAGCAAGTTAAAATAAGGC |
| 127 | psg1454_zwf | 9036 | CCAAAGATTGTGATTAAACAgttttAGAGCTAGAAATAGCAAGTTAAAATAAGGC |
| 128 | psg1459_gnd | 9037 | AGCTAGGTTTTTACCCATAAgttttAGAGCTAGAAATAGCAAGTTAAAATAAGGC |
| 129 | psg1464_bmfBB | 9038 | TGAACACTCTCACCTAACTTgttttAGAGCTAGAAATAGCAAGTTAAAATAAGGC |
| 130 | psg1466_bmfBAA | 9039 | CTTTTAGGTCTTCTTCGCTAgttttAGAGCTAGAAATAGCAAGTTAAAATAAGGC |

| | | | |
|---|---|---|---|
| 131 | psg1475_accC | 9040 | TCCTAACTGCGATTTCACCGgttttAGAGCTAGAAATAGCAAGTTAAAATAAGGC |
| 132 | psg1490_efp | 9041 | CTTTACCAGGCTTTACATGTgttttAGAGCTAGAAATAGCAAGTTAAAATAAGGC |
| 133 | psg1492 | 8530 | TCTGTTGTTTTTTCTTCTAAgttttAGAGCTAGAAATAGCAAGTTAAAATAAGGC |
| 134 | psg1520_trmK | 8531 | ATTCTCCTTTATGAAAAAAGgttttAGAGCTAGAAATAGCAAGTTAAAATAAGGC |
| 135 | psg1521_rpoD | 8532 | CTGTGTTATCAGACATGAAAgttttAGAGCTAGAAATAGCAAGTTAAAATAAGGC |
| 136 | psg1522_dnaG | 8533 | ATTTATTTTGTTCAATATAAgttttAGAGCTAGAAATAGCAAGTTAAAATAAGGC |
| 137 | psg1525_glyS | 8534 | TCATACATGAAAACGCCCCAgttttAGAGCTAGAAATAGCAAGTTAAAATAAGGC |
| 138 | psg1530_ybeY | 8535 | TCAATGATCTTACTTACCAAgttttAGAGCTAGAAATAGCAAGTTAAAATAAGGC |
| 139 | psg1540_dnaK | 8536 | TCAGGGTTTTGAATTACTTTgttttAGAGCTAGAAATAGCAAGTTAAAATAAGGC |
| 140 | psg1541_grpE | 8537 | TCATTAATTTTTTGATCTTTgttttAGAGCTAGAAATAGCAAGTTAAAATAAGGC |
| 141 | psg1546_holA | 8538 | CTTTGTTTTTCAACCAATTCgttttAGAGCTAGAAATAGCAAGTTAAAATAAGGC |
| 142 | psg1557 | 8539 | ATTGATTGAACATATGAATTgttttAGAGCTAGAAATAGCAAGTTAAAATAAGGC |
| 143 | psg1567_greA | 8540 | TCAAAACCTTCTTGAGTCATgttttAGAGCTAGAAATAGCAAGTTAAAATAAGGC |
| 144 | psg1575_alaS | 8541 | ATTGGCACTAATGGTGCAGAgttttAGAGCTAGAAATAGCAAGTTAAAATAAGGC |
| 145 | psg1579_iscS | 8738 | TACTTCAGGTTTTACTGGTGgttttAGAGCTAGAAATAGCAAGTTAAAATAAGGC |
| 146 | psg1587_hisS | 8739 | AAAATATCCTGCGTCCCTCTgttttAGAGCTAGAAATAGCAAGTTAAAATAAGGC |
| 147 | psg1589_dtd | 8740 | GTATTTCACCATTCATTTGTgttttAGAGCTAGAAATAGCAAGTTAAAATAAGGC |
| 148 | psg1593_secF | 8741 | TTTATAAGTTGCAGCCATTCgttttAGAGCTAGAAATAGCAAGTTAAAATAAGGC |
| 149 | psg1600_obgE | 8742 | AATACCATTACCACCATCACgttttAGAGCTAGAAATAGCAAGTTAAAATAAGGC |
| 150 | psg1602 | 8743 | CATATTCACCATGGTCAGCAgttttAGAGCTAGAAATAGCAAGTTAAAATAAGGC |
| 151 | psg1611_valS | 8744 | ACTTCACGAGGATCATATTTgttttAGAGCTAGAAATAGCAAGTTAAAATAAGGC |
| 152 | psg1619_hemA | 8745 | ATCTTCATGGGCAATTCGTAgttttAGAGCTAGAAATAGCAAGTTAAAATAAGGC |
| 153 | psg1620_engB | 8746 | ATGATTAATTCTATATTATTgttttAGAGCTAGAAATAGCAAGTTAAAATAAGGC |
| 154 | psg1624 | 8747 | TTATTAAAAATAATTTCTCTgttttAGAGCTAGAAATAGCAAGTTAAAATAAGGC |
| 155 | psg1627_infC | 8748 | TTGAGTTTGATCTTTTGCTAgttttAGAGCTAGAAATAGCAAGTTAAAATAAGGC |
| 156 | psg1629_thrS | 8749 | AACGCCTTTTTATTACCATCgttttAGAGCTAGAAATAGCAAGTTAAAATAAGGC |
| 157 | psg1631_dnaB | 8792 | TGGTCTTAAGCCGAATTCGAgttttAGAGCTAGAAATAGCAAGTTAAAATAAGGC |
| 158 | psg1632_nrdR | 8793 | ATTGTGTAGAATTACATTTCgttttAGAGCTAGAAATAGCAAGTTAAAATAAGGC |
| 159 | psg1634_coaE | 8794 | CCTGTTAGACCAATAACTTTgttttAGAGCTAGAAATAGCAAGTTAAAATAAGGC |
| 160 | psg1645_pfkA | 8795 | TGTACGAACAACTGCTCTTAgttttAGAGCTAGAAATAGCAAGTTAAAATAAGGC |
| 161 | psg1647_accD | 8796 | CACTTAGTCATAATACCTGCgttttAGAGCTAGAAATAGCAAGTTAAAATAAGGC |
| 162 | psg1649_dnaE | 8797 | TTCAGACACAGCAAGTCTTAgttttAGAGCTAGAAATAGCAAGTTAAAATAAGGC |
| 163 | psg1657_ackA | 8798 | TTTGTTACTAATTCCTCTTCgttttAGAGCTAGAAATAGCAAGTTAAAATAAGGC |

| | | | |
|---|---|---|---|
| 164 | psg1673_plsC | 8799 | ACGACATATTTACTATCCTTgttttAGAGCTAGAAATAGCAAGTTAAAATAAGGC |
| 165 | psg1675_tyrS | 8800 | TAAACTATCTGCCGTTGGATgttttAGAGCTAGAAATAGCAAGTTAAAATAAGGC |
| 166 | psg1686_murC | 8801 | TTTATGTTATTAGCATCAAAgttttAGAGCTAGAAATAGCAAGTTAAAATAAGGC |
| 167 | psg1688 | 8802 | GCGACATCTCCTACATATTTgttttAGAGCTAGAAATAGCAAGTTAAAATAAGGC |
| 168 | psg1704_leuS | 8803 | ATTTCTTTTCAATTTGATTGgttttAGAGCTAGAAATAGCAAGTTAAAATAAGGC |
| 169 | psg1729 | 8804 | ATAATGAGTCCAATGACTATgttttAGAGCTAGAAATAGCAAGTTAAAATAAGGC |
| 170 | psg1730_metK | 8805 | CTTGGTCAGCGATTTTATCTgttttAGAGCTAGAAATAGCAAGTTAAAATAAGGC |
| 171 | psg1748 | 8806 | TCGCGTATAATTTCATTCCTgttttAGAGCTAGAAATAGCAAGTTAAAATAAGGC |
| 172 | psg1749 | 8807 | TAATAGAATAACCATCCATTgttttAGAGCTAGAAATAGCAAGTTAAAATAAGGC |
| 173 | psg1869_map | 8808 | TTTCGTAGTGATACCTGGTTgttttAGAGCTAGAAATAGCAAGTTAAAATAAGGC |
| 174 | psg1873_murT | 8809 | TACGCGCCAATTTCGCTAGAgttttAGAGCTAGAAATAGCAAGTTAAAATAAGGC |
| 175 | psg1879_dgkB | 8810 | ATAGCTCTTTACCTGATGTCgttttAGAGCTAGAAATAGCAAGTTAAAATAAGGC |
| 176 | psg1881_gatA | 8811 | ATATCTTTAACAACATCAGAgttttAGAGCTAGAAATAGCAAGTTAAAATAAGGC |
| 177 | psg1882_gatC | 8812 | GCCATTTCTTCCGTTTCTTCgttttAGAGCTAGAAATAGCAAGTTAAAATAAGGC |
| 178 | psg1885_ligA | 8813 | TATTCACTATCTGGTACAGAgttttAGAGCTAGAAATAGCAAGTTAAAATAAGGC |
| 179 | psg1886_pcrA | 8814 | GCACCTGCCATAATTAACAAgttttAGAGCTAGAAATAGCAAGTTAAAATAAGGC |
| 180 | psg1894_pncB | 8815 | TTAAACTGTCGTCTTCTAATgttttAGAGCTAGAAATAGCAAGTTAAAATAAGGC |
| 181 | psg1900_ppaC | 8970 | GATGAAATTGCATCAGTGTCgttttAGAGCTAGAAATAGCAAGTTAAAATAAGGC |
| 182 | psg1911_pmtC | 8971 | ACCATTCATTACTTTCATAAgttttAGAGCTAGAAATAGCAAGTTAAAATAAGGC |
| 183 | psg1913_pmtA | 8972 | ATTAACATTACTTAATTCTAgttttAGAGCTAGAAATAGCAAGTTAAAATAAGGC |
| 184 | psg1983_groES | 8973 | ATAATCACACGATTTCCAATgttttAGAGCTAGAAATAGCAAGTTAAAATAAGGC |
| 185 | psg1988_hld | 8974 | ACTAGATCACAGAGATGTGAgttttAGAGCTAGAAATAGCAAGTTAAAATAAGGC |
| 186 | psg2002_gcp | 8975 | TGATGTCTACTTGCCACTTCgttttAGAGCTAGAAATAGCAAGTTAAAATAAGGC |
| 187 | psg2005_tsaE | 8976 | TTTAATGATGTTAAATGTCGgttttAGAGCTAGAAATAGCAAGTTAAAATAAGGC |
| 188 | psg2028_acpS | 8977 | AAAATCCGCTCAACCAATTTgttttAGAGCTAGAAATAGCAAGTTAAAATAAGGC |
| 189 | psg2039_ddl | 8978 | CTTTCTCCAATCACCATCATgttttAGAGCTAGAAATAGCAAGTTAAAATAAGGC |
| 190 | psg2046_oxaA | 8979 | ACCATAATACCTAAAAATAAgttttAGAGCTAGAAATAGCAAGTTAAAATAAGGC |
| 191 | psg2055_murA | 8980 | AATAAAGATGCTGTCAATATgttttAGAGCTAGAAATAGCAAGTTAAAATAAGGC |
| 192 | psg2062_atpF | 8981 | TAAGTTAGCTGTTTCAGTCAgttttAGAGCTAGAAATAGCAAGTTAAAATAAGGC |
| 193 | psg2072_prfA | 9003 | CTGAATCATTTACAACATCTgttttAGAGCTAGAAATAGCAAGTTAAAATAAGGC |
| 194 | psg2079_fbaA | 9004 | ATTTCTTTCATTGAAACTAAgttttAGAGCTAGAAATAGCAAGTTAAAATAAGGC |
| 195 | psg2081_pyrG | 9005 | CCTGGGTCAACATTTAAGTAgttttAGAGCTAGAAATAGCAAGTTAAAATAAGGC |
| 196 | psg2084_coaA | 9006 | ATTCAGTTTTAAAAGTACGTgttttAGAGCTAGAAATAGCAAGTTAAAATAAGGC |

| | | | |
|---|---|---|---|
| 197 | psg2104_glmS | 9007 | ACCTTTTAATAATAATTCTTgttttAGAGCTAGAAATAGCAAGTTAAAATAAGGC |
| 198 | psg2113_dacA | 9008 | ACTGAGGTTTTGAAAAAAGTgttttAGAGCTAGAAATAGCAAGTTAAAATAAGGC |
| 199 | psg2182_infA | 9009 | AACGCAATGTTTAAAGTAGAgttttAGAGCTAGAAATAGCAAGTTAAAATAAGGC |
| 200 | psg2184_secY | 9010 | TGTTCTAAAGAAGTTCACAAgttttAGAGCTAGAAATAGCAAGTTAAAATAAGGC |
| 201 | psg2214_femX | 9011 | TAATAAATCTCCATTTGGGTgttttAGAGCTAGAAATAGCAAGTTAAAATAAGGC |
| 202 | psg2238_ureA | 9012 | CTGATTAAAGCTAATGCCTCgttttAGAGCTAGAAATAGCAAGTTAAAATAAGGC |
| 203 | psg2283_rpiA | 9013 | TTAATTAGTTGCGCCATTTGgttttAGAGCTAGAAATAGCAAGTTAAAATAAGGC |
| 204 | psg2292_fni | 9014 | TGAATGCATTGCGTCAGATTgttttAGAGCTAGAAATAGCAAGTTAAAATAAGGC |
| 205 | psg2483_mvaA | 9015 | ATTCTTATCTAAATTTTGCAgttttAGAGCTAGAAATAGCAAGTTAAAATAAGGC |
| 206 | psg2646_trmE | 9017 | CCAATTGCCCCTTCACCCATgttttAGAGCTAGAAATAGCAAGTTAAAATAAGGC |
| 207 | psg2484_mvaS | 9016 | CCATGTCTACATAGTACTTgttttAGAGCTAGAAATAGCAAGTTAAAATAAGGC |
| 208 | psg0005_gyrB | 9484 | TGATTAATACGATACAATTTgttttAGAGCTAGAAATAGCAAGTTAAAATAAGGC |
| 209 | psg0527_rpoB | 9485 | GTTTACGATGTCTTCCATATgttttAGAGCTAGAAATAGCAAGTTAAAATAAGGC |
| 210 | psg0425_mpsA | 9486 | ATAGTAGATTCTGTACATAAgttttAGAGCTAGAAATAGCAAGTTAAAATAAGGC |
| 211 | psg0639 | 9487 | TCAATACATGTTTTTTTATAgttttAGAGCTAGAAATAGCAAGTTAAAATAAGGC |
| 212 | psg1359 | 9488 | TTTGCCACGTTAATCACCTTgttttAGAGCTAGAAATAGCAAGTTAAAATAAGGC |
| 213 | psg1590_relA | 9489 | TTCGCCTTAAACAATGATTTgttttAGAGCTAGAAATAGCAAGTTAAAATAAGGC |
| 214 | psg1791_cbf1 | 9490 | AATGTAATCACTTCTTTTGAgttttAGAGCTAGAAATAGCAAGTTAAAATAAGGC |
| 215 | psg2183_adk | 9491 | TGAGTTCCTTTACCTGCGCCgttttAGAGCTAGAAATAGCAAGTTAAAATAAGGC |
| 216 | psg2185_rplO | 9492 | TAACTCATGTAATTTCATTTgttttAGAGCTAGAAATAGCAAGTTAAAATAAGGC |
| 217 | psg2647_rnpA | 9493 | AATCTGCATTCTTTTTAATTgttttAGAGCTAGAAATAGCAAGTTAAAATAAGGC |
| 218 | psg2648_rpmH | 9494 | TTACTATGTTTACGTTTATTgttttAGAGCTAGAAATAGCAAGTTAAAATAAGGC |
| 219 | psg0015_rplI | 9495 | AAGTTATTTGCATAACCTACgttttAGAGCTAGAAATAGCAAGTTAAAATAAGGC |
| 220 | psg0366_rpsF | 9496 | TTTATATTTGCACCTCCTTGgttttAGAGCTAGAAATAGCAAGTTAAAATAAGGC |
| 221 | psg0479_rplY | 9497 | AACGTGTTTGTTTACCTTGAgttttAGAGCTAGAAATAGCAAGTTAAAATAAGGC |
| 222 | psg0522_rplK | 9498 | TGATATCGTGATGTGGTCACgttttAGAGCTAGAAATAGCAAGTTAAAATAAGGC |
| 223 | psg0524_rplJ | 9499 | GACACCTCCATTTAAATTTTgttttAGAGCTAGAAATAGCAAGTTAAAATAAGGC |
| 224 | psg0530_rpsL | 9500 | CGTACTAATTGGTTAATAGTgttttAGAGCTAGAAATAGCAAGTTAAAATAAGGC |
| 225 | psg0990_rpoY | 9501 | TAACCCTAGTAAAATCGTATgttttAGAGCTAGAAATAGCAAGTTAAAATAAGGC |
| 226 | psg1027_rpmF | 9502 | GTTTTAGAAGTTCTTCTTTTgttttAGAGCTAGAAATAGCAAGTTAAAATAAGGC |
| 227 | psg1117_rpmB | 9503 | CAGTTTACTCAAAATATAATgttttAGAGCTAGAAATAGCAAGTTAAAATAAGGC |
| 228 | psg1131_rpsP | 9504 | GCTACTACGATACGATAGAAgttttAGAGCTAGAAATAGCAAGTTAAAATAAGGC |
| 229 | psg1134_rplS | 9505 | CCTCGACAAATATATAGCAGgttttAGAGCTAGAAATAGCAAGTTAAAATAAGGC |

| | | | |
|---|---|---|---|
| 230 | psg1149_rpsB | 9506 | CCATTATAAATTCCTCCTATgttttAGAGCTAGAAATAGCAAGTTAAAATAAGGC |
| 231 | psg1166_rpsO | 9507 | AATACTCCTTAATCCGAGTTgttttAGAGCTAGAAATAGCAAGTTAAAATAAGGC |
| 232 | psg1511_rpmG | 9508 | TAAAGTTACGTTTACGCGCAgttttAGAGCTAGAAATAGCAAGTTAAAATAAGGC |
| 233 | psg1535_rpsU | 9509 | TCCCTCCCTCCAAATATCAAgttttAGAGCTAGAAATAGCAAGTTAAAATAAGGC |
| 234 | psg1545_rpsT | 9510 | TTGTGCAATCATTTATTTGAgttttAGAGCTAGAAATAGCAAGTTAAAATAAGGC |
| 235 | psg1601_rpmA | 9511 | TGTAAGTTTAATTTTAACATgttttAGAGCTAGAAATAGCAAGTTAAAATAAGGC |
| 236 | psg1625_rplT | 9512 | GGCATAAAAAATTCCTCCTTgttttAGAGCTAGAAATAGCAAGTTAAAATAAGGC |
| 237 | psg1666_rpsD | 9513 | CCATGTTGTCCTGGTGCGTAgttttAGAGCTAGAAATAGCAAGTTAAAATAAGGC |
| 238 | psg2074_rpmE2 | 9514 | CTCCTTTGCCCTGAACCATCgttttAGAGCTAGAAATAGCAAGTTAAAATAAGGC |
| 239 | psg2171_rpsI | 9515 | CCACGTAATTCGTAGTTTTCgttttAGAGCTAGAAATAGCAAGTTAAAATAAGGC |
| 240 | psg2177_rplQ | 9516 | GAAAAGAAGATTGATAAAGGgttttAGAGCTAGAAATAGCAAGTTAAAATAAGGC |
| 241 | psg0386_xpt | 10048 | CCAAACTAAATAATAGTTTCgttttAGAGCTAGAAATAGCAAGTTAAAATAAGGC |
| 242 | psg0529_rplGB | 10050 | ATTTACATAAAAATAACAAGgttttAGAGCTAGAAATAGCAAGTTAAAATAAGGC |
| 243 | psg0755_gapR | 10052 | ATTTACATAAAAATAACAAGgttttAGAGCTAGAAATAGCAAGTTAAAATAAGGC |
| 244 | psg0906 | 10054 | CTTACGAACACTATTTTTAAgttttAGAGCTAGAAATAGCAAGTTAAAATAAGGC |
| 245 | psg1072_mraZ | 10056 | GATTTATCATTAAGATAAATgttttAGAGCTAGAAATAGCAAGTTAAAATAAGGC |
| 246 | psg1108_def | 10058 | GTAATCCTAAATTTATTGTAgttttAGAGCTAGAAATAGCAAGTTAAAATAAGGC |
| 247 | psg1121_fapR | 10060 | AGTTCATGGTCTGTGATGAAgttttAGAGCTAGAAATAGCAAGTTAAAATAAGGC |
| 248 | psg1155_rasP | 10062 | CCCATACCGATCGCAAATTCgttttAGAGCTAGAAATAGCAAGTTAAAATAAGGC |
| 249 | psg1158_rimP | 10064 | AAATTCAAGTCTTCCATAATgttttAGAGCTAGAAATAGCAAGTTAAAATAAGGC |
| 250 | psg1558_mtnM | 10066 | TATTGTTACTTCTTCTTCCAgttttAGAGCTAGAAATAGCAAGTTAAAATAAGGC |
| 251 | psg1635_mutM | 10068 | CTTTTTACATGTTCTACTTCgttttAGAGCTAGAAATAGCAAGTTAAAATAAGGC |
| 252 | psg1690 | 10070 | GGTTCTATCACTCTACAATCgttttAGAGCTAGAAATAGCAAGTTAAAATAAGGC |
| 253 | psg1914_pmtR | 10072 | TGCTTAATCTGTTCATAAATgttttAGAGCTAGAAATAGCAAGTTAAAATAAGGC |
| 254 | psg2029 | 10074 | CTCATCACTTTTTTAGCGTGgttttAGAGCTAGAAATAGCAAGTTAAAATAAGGC |
| 255 | psg2205_rpsJ | 10076 | CTGCTGATTGATCAATTACGgttttAGAGCTAGAAATAGCAAGTTAAAATAAGGC |
| 256 | psg0014_gdpP | 10078 | TATTAGTAAAGCTTTCTTAGgttttAGAGCTAGAAATAGCAAGTTAAAATAAGGC |
| 257 | psg1603_rplU | 10080 | TAATAAGTCACGCCATACATgttttAGAGCTAGAAATAGCAAGTTAAAATAAGGC |
| 258 | psg2171_rplM | 10082 | ATGATAAACGACCTAATGTTgttttAGAGCTAGAAATAGCAAGTTAAAATAAGGC |
| 259 | psg1476_accB | 9891 | TTCAAGCATGTTCATATTGCgttttAGAGCTAGAAATAGCAAGTTAAAATAAGGC |
| 260 | psg1700_murJ | 6423 | CTTGGTAATTAATATACTAAgttttAGAGCTAGAAATAGCAAGTTAAAATAAGGC |
| 261 | psg1080_ftsZ | 6424 | AAATTTCCTCCTAGTTTTAgttttAGAGCTAGAAATAGCAAGTTAAAATAAGGC |
| | **B Clones** | | |
| 4B | psg0009_serS_B | 9062 | ATTTTGCTCTTAACTGTGTCgttttAGAGCTAGAAATAGCAAGTTAAAATAAGGC |
| 6B | psg0020_walR_B | 9063 | AATTCTAAAATATCAGCAATgttttAGAGCTAGAAATAGCAAGTTAAAATAAGGC |

| | | | |
|---|---|---|---|
| 7B | psg0024_walJ_B | 9064 | ATACACTCATGCGTATCAAGgttttAGAGCTAGAAATAGCAAGTTAAAATAAGGC |
| 8B | psg0089_B | 9065 | AATTCAGTGAAAAACACATCgttttAGAGCTAGAAATAGCAAGTTAAAATAAGGC |
| 9B | psg0248_tarF_B | 9066 | AAATAAATTAGATTCTTGTTgttttAGAGCTAGAAATAGCAAGTTAAAATAAGGC |
| 10B | psg0249_ispD_B | 9276 | TCTAATGTATGGATTAAAATgttttAGAGCTAGAAATAGCAAGTTAAAATAAGGC |
| 15B | psg0453_B | 9546 | GTTTGTTGTGCTTGTTGCTTgttttAGAGCTAGAAATAGCAAGTTAAAATAAGGC |
| 24B | psg0487_tilS_B | 8545 | TAGAAACAGCGACAACAATAgttttAGAGCTAGAAATAGCAAGTTAAAATAAGGC |
| 25B | psg0492_folP_B | 8695 | GTCAGCACCTTCATCTATCAgttttAGAGCTAGAAATAGCAAGTTAAAATAAGGC |
| 28B | psg0514_cysE_B | 8696 | TCTAATGTTGAACGTGCCGCgttttAGAGCTAGAAATAGCAAGTTAAAATAAGGC |
| 31B | psg0551_folE2_B | 8697 | GTCATTTCGTTCTTAGTAGTgttttAGAGCTAGAAATAGCAAGTTAAAATAAGGC |
| 32B | psg0570_eutD_B | 8698 | ACACGTTCGTCCTCTCCTTCgttttAGAGCTAGAAATAGCAAGTTAAAATAAGGC |
| 33B | psg0572_mvaK1_B | 9277 | CGAATAGTTCCCGCTCTCTAgttttAGAGCTAGAAATAGCAAGTTAAAATAAGGC |
| 34B | psg0596_argS_B | 8699 | GGAACTTCAATTTTAATATCgttttAGAGCTAGAAATAGCAAGTTAAAATAAGGC |
| 35B | psg0623_tagA_B | 8700 | CGTTGATTGATTTGCAAAAAgttttAGAGCTAGAAATAGCAAGTTAAAATAAGGC |
| 36B | psg0624_tagH_B | 8701 | AAATGTTTTGTTTTTATGTTgttttAGAGCTAGAAATAGCAAGTTAAAATAAGGC |
| 40B | psg0703_ltaS_B | 9278 | CTTTGATTTAACAAGTATTTgttttAGAGCTAGAAATAGCAAGTTAAAATAAGGC |
| 42B | psg0722_murB_B | 9210 | TGATATTAAAGACATGAGAAgttttAGAGCTAGAAATAGCAAGTTAAAATAAGGC |
| 46B | psg0743_hprK_B | 9213 | AGTTTTTCTGTCGTTAACATgttttAGAGCTAGAAATAGCAAGTTAAAATAAGGC |
| 55B | psg0868_spsB_B | 9216 | GATTCACCTTTAATTGTATAgttttAGAGCTAGAAATAGCAAGTTAAAATAAGGC |
| 56B | psg0885_fabH_B | 9217 | TTAATAAAATTTTTAATACCgttttAGAGCTAGAAATAGCAAGTTAAAATAAGGC |
| 62B | psg0922_B | 9220 | AAATAAGGTAAGATCAAACTgttttAGAGCTAGAAATAGCAAGTTAAAATAAGGC |
| 65B | psg0983_ptsH_B | 9221 | AAATTCAATTCGTTTAAAACgttttAGAGCTAGAAATAGCAAGTTAAAATAAGGC |
| 66B | psg0989_rnjA_B | 9286 | ATCCCTAATAAGTTATCATCgttttAGAGCTAGAAATAGCAAGTTAAAATAAGGC |
| 67B | psg0990_rpoY_B | 9287 | GGATAAAAAGTTTGGTAGAAgttttAGAGCTAGAAATAGCAAGTTAAAATAAGGC |
| 68B | psg0991_def _B | 9288 | TTAAAAAATGATTAAAGTGTgttttAGAGCTAGAAATAGCAAGTTAAAATAAGGC |
| 70B | psg1024_coaD_B | 9289 | TAACGTTTAATCATATTAAAgttttAGAGCTAGAAATAGCAAGTTAAAATAAGGC |
| 92B | psg1133_trmD_B | 9291 | TTTACAACATCAGCAATATAgttttAGAGCTAGAAATAGCAAGTTAAAATAAGGC |
| 97B | psg1152_frr_B | 8999 | GCTAATTGTTGTACAGGTGTgttttAGAGCTAGAAATAGCAAGTTAAAATAAGGC |
| 98B | psg1153_uppS_B | 9292 | TCGTAATGACCTTTAATTCTgttttAGAGCTAGAAATAGCAAGTTAAAATAAGGC |
| 100B | psg1157_polC_B | 9293 | AAAATTAAATGCTTTATATTgttttAGAGCTAGAAATAGCAAGTTAAAATAAGGC |
| 103B | psg1176_pgsA_B | 9294 | AACCATCAACAAAATCGCTAgttttAGAGCTAGAAATAGCAAGTTAAAATAAGGC |
| 107B | psg1249_plsY_B | 9295 | TATCATATTCATTTAAATTAgttttAGAGCTAGAAATAGCAAGTTAAAATAAGGC |
| 109B | psg1257_msrR_B | 9296 | TTTCTTAATAAATCGTACTAgttttAGAGCTAGAAATAGCAAGTTAAAATAAGGC |
| 115B | psg1340_recU_B | 9299 | AATACATTTACCGTTAATGTgttttAGAGCTAGAAATAGCAAGTTAAAATAAGGC |

| | | | |
|---|---|---|---|
| 116B | psg1341_pbp2_B | 9300 | AATTTAGCTTCGGTAAAAGCgttttAGAGCTAGAAATAGCAAGTTAAAATAAGGC |
| 117B | psg1344_dnaD_B | 9301 | GATTTTAACAATGAACTTTAgttttAGAGCTAGAAATAGCAAGTTAAAATAAGGC |
| 118B | psg1347_birA_B | 9302 | ATACCTTGATACCAAATATCgttttAGAGCTAGAAATAGCAAGTTAAAATAAGGC |
| 119B | psg1348_papS_B | 9303 | CGAACAAATGTAACACCACTgttttAGAGCTAGAAATAGCAAGTTAAAATAAGGC |
| 120B | psg1351_B | 9304 | GTAGTTATAAAATCATAATAgttttAGAGCTAGAAATAGCAAGTTAAAATAAGGC |
| 121B | psg1360_ubiE_B | 9305 | ATACCAGTAACTTCACCTGTgttttAGAGCTAGAAATAGCAAGTTAAAATAAGGC |
| 129B | psg1464_bmfBB_B | 9990 | ACAGGAGAAAATGGCATAGAgttttAGAGCTAGAAATAGCAAGTTAAAATAAGGC |
| 131B | psg1475_accC_B | 9308 | TGCACATATGCTTCTTCGTCgttttAGAGCTAGAAATAGCAAGTTAAAATAAGGC |
| 132B | psg1490_efp_B | 9309 | ACGCTTATGTTAAAACTATAgttttAGAGCTAGAAATAGCAAGTTAAAATAAGGC |
| 133B | psg1492_B | 9552 | AAATGCACAAACGTTTCACTgttttAGAGCTAGAAATAGCAAGTTAAAATAAGGC |
| 134B | psg1520_trmK_B | 9553 | TTCAAACGTTTACTACGACTgttttAGAGCTAGAAATAGCAAGTTAAAATAAGGC |
| 136B | psg1522_dnaG_B | 9067 | ACTTACCAAGTCTAAAATGTgttttAGAGCTAGAAATAGCAAGTTAAAATAAGGC |
| 137B | psg1525_glyS_B | 9068 | CACCGTAAATATCACTACCAgttttAGAGCTAGAAATAGCAAGTTAAAATAAGGC |
| 139B | psg1540_dnaK_B | 9554 | TTTCGCCAAATTTAAGTTATgttttAGAGCTAGAAATAGCAAGTTAAAATAAGGC |
| 140B | psg1541_grpE_B | 9069 | CGTTCTATATTGTCTATTGCgttttAGAGCTAGAAATAGCAAGTTAAAATAAGGC |
| 141B | psg1546_holA_B | 9070 | GTTAATGTTTCTTCAACAATgttttAGAGCTAGAAATAGCAAGTTAAAATAAGGC |
| 143B | psg1567_greA_B | 9071 | TTAATTTTCTCTACAACTTCgttttAGAGCTAGAAATAGCAAGTTAAAATAAGGC |
| 144B | psg1575_alaS_B | 9072 | TTTACAATTCTTGGCTTTTTgttttAGAGCTAGAAATAGCAAGTTAAAATAAGGC |
| 147B | psg1589_dtd_B | 9555 | TGAAACTTTAATTAAATTGCgttttAGAGCTAGAAATAGCAAGTTAAAATAAGGC |
| 148B | psg1593_secF_B | 9073 | ATTTTGTCGCCTTTATTTAAgttttAGAGCTAGAAATAGCAAGTTAAAATAAGGC |
| 151B | psg1611_valS_B | 9323 | TACCAGTTACATTTGGTGGCgttttAGAGCTAGAAATAGCAAGTTAAAATAAGGC |
| 152B | psg1619_hemA_B | 9324 | TACAAAAACTATGAAATATAgttttAGAGCTAGAAATAGCAAGTTAAAATAAGGC |
| 154B | psg1624_B | 9325 | AAATGAGTTGTTTATATGAgttttAGAGCTAGAAATAGCAAGTTAAAATAAGGC |
| 156B | psg1629_thrS_B | 9326 | ATTGATCCATCAGTTTCAAGgttttAGAGCTAGAAATAGCAAGTTAAAATAAGGC |
| 157B | psg1631_dnaB_B | 9327 | ACTGCTTGCGTTCCAATTAGgttttAGAGCTAGAAATAGCAAGTTAAAATAAGGC |
| 158B | psg1632_nrdR_B | 9328 | CTAACTCCGAAGTCAGAGTTgttttAGAGCTAGAAATAGCAAGTTAAAATAAGGC |
| 159B | psg1634_coaE_B | 9329 | CTTCCACACACTTTGCATACgttttAGAGCTAGAAATAGCAAGTTAAAATAAGGC |
| 161B | psg1647_accD_B | 9330 | TGAACTTGACCAGTTAAAAAgttttAGAGCTAGAAATAGCAAGTTAAAATAAGGC |
| 162B | psg1649_dnaE_B | 9331 | ACTGAAACTCCTGACGCATTgttttAGAGCTAGAAATAGCAAGTTAAAATAAGGC |
| 163B | psg1657_ackA_B | 9332 | ACTATTACAGATTATTTTTTgttttAGAGCTAGAAATAGCAAGTTAAAATAAGGC |
| 165B | psg1675_tyrS_B | 9333 | ACCCTTACAACAAATATGTAgttttAGAGCTAGAAATAGCAAGTTAAAATAAGGC |
| 168B | psg1704_leuS_B | 9334 | AAACCAGCACCTGATGGATAgttttAGAGCTAGAAATAGCAAGTTAAAATAAGGC |
| 169B | psg1729_B | 9335 | ACTTTGTCTAATTTTTCCCAgttttAGAGCTAGAAATAGCAAGTTAAAATAAGGC |

| | | | |
|---|---|---|---|
| 170B | psg1730_metK_B | 9336 | TGTTGTTGTAGAAATTTCGCgttttAGAGCTAGAAATAGCAAGTTAAAATAAGGC |
| 171B | psg1748_B | 9337 | AGTAGAGTCGCCTATCTCTCgttttAGAGCTAGAAATAGCAAGTTAAAATAAGGC |
| 172B | psg1749_B | 9338 | GTTACAACCCATATGATTGTgttttAGAGCTAGAAATAGCAAGTTAAAATAAGGC |
| 173B | psg1869_map_B | 9339 | AAGATGTTACTTTAGTATTTgttttAGAGCTAGAAATAGCAAGTTAAAATAAGGC |
| 180B | psg1894_pncB_B | 9340 | GTAGACTATAATATAAAGCGgttttAGAGCTAGAAATAGCAAGTTAAAATAAGGC |
| 181B | psg1900_ppaC_B | 9341 | AATGAAGCCACTCCCTCAGCgttttAGAGCTAGAAATAGCAAGTTAAAATAAGGC |
| 182B | psg1911_pmtC_B | 9342 | AAAATATATGAATATAAATCgttttAGAGCTAGAAATAGCAAGTTAAAATAAGGC |
| 185B | psg1988_hld_B | 9345 | AGTATTTATTTCCTACAGTTgttttAGAGCTAGAAATAGCAAGTTAAAATAAGGC |
| 188B | psg2028_acpS_B | 9346 | CTTGACCACCCGCTGTATAAgttttAGAGCTAGAAATAGCAAGTTAAAATAAGGC |
| 191B | psg2055_murA_B | 9347 | ATTTTACTAACATATTCATAgttttAGAGCTAGAAATAGCAAGTTAAAATAAGGC |
| 192B | psg2062_atpF_B | 9348 | CTGAATTTATGCGATAGGCAgttttAGAGCTAGAAATAGCAAGTTAAAATAAGGC |
| 198B | psg2113_dacA_B | 9559 | TGATTATTACTGTCAATCAAgttttAGAGCTAGAAATAGCAAGTTAAAATAAGGC |
| 200B | psg2184_secY_B | 9560 | CCAGATTCTACTAATAAAGCgttttAGAGCTAGAAATAGCAAGTTAAAATAAGGC |
| 202B | psg2238_ureA_B | 9349 | ATCTAATCGAAAACAAATAGgttttAGAGCTAGAAATAGCAAGTTAAAATAAGGC |
| 203B | psg2283_rpiA_B | 9350 | ATTTTAAATAATACTCGTTAgttttAGAGCTAGAAATAGCAAGTTAAAATAAGGC |
| 208B | psg005_gyrB_B | 9608 | GTCGATCCTATATACATACCgttttAGAGCTAGAAATAGCAAGTTAAAATAAGGC |
| 209B | psg0527_rpoB_B | 9609 | ATAAAAAGACAAAAAGAAAAgttttAGAGCTAGAAATAGCAAGTTAAAATAAGGC |
| 211B | psg0639_B | 9611 | ATATATCTCATTGGCATAACgttttAGAGCTAGAAATAGCAAGTTAAAATAAGGC |
| 212B | psg1359_B | 9612 | TACTCAGAATAACAAATGCTgttttAGAGCTAGAAATAGCAAGTTAAAATAAGGC |
| 213B | psg1590_relA_B | 9613 | TGTATGGTAATCCGTTTTTTgttttAGAGCTAGAAATAGCAAGTTAAAATAAGGC |
| 214B | psg1791_cbf1_B | 9614 | TTAACATGTACAATTTCTTCgttttAGAGCTAGAAATAGCAAGTTAAAATAAGGC |
| 219B | psg0015_rplL_B | 9615 | TTTAAGTTGTGTTGCCGCATgttttAGAGCTAGAAATAGCAAGTTAAAATAAGGC |
| 221B | psg0479_rplY_B | 9675 | ACGAAACTATTATACACGTTgttttAGAGCTAGAAATAGCAAGTTAAAATAAGGC |
| 222B | psg0522_rplK_B | 9676 | ACGAAACTATTATACACGTTgttttAGAGCTAGAAATAGCAAGTTAAAATAAGGC |
| 223B | psg0524_rplJ_B | 9616 | CTTCAGCTACTGTTAATCCAgttttAGAGCTAGAAATAGCAAGTTAAAATAAGGC |
| 225B | psg0990_rpoY_B | 9287 | TAACGATTTGGTTAATAACTgttttAGAGCTAGAAATAGCAAGTTAAAATAAGGC |
| 231B | psg1166_rpsO_B | 9677 | AGTACAGCGATTTGTACTTCgttttAGAGCTAGAAATAGCAAGTTAAAATAAGGC |
| 234B | psg1545_rpsT_B | 9620 | CTTTTAGGAGGTGACAGAAAgttttAGAGCTAGAAATAGCAAGTTAAAATAAGGC |
| 235B | psg1601_rpmA_B | 9621 | TTGTCGTCATAATTGATATCgttttAGAGCTAGAAATAGCAAGTTAAAATAAGGC |
| 236B | psg1625_rplT_B | 9622 | TAACACGTTTAGCTGCTCCGgttttAGAGCTAGAAATAGCAAGTTAAAATAAGGC |
| 237B | psg1666_rpsD_B | 9623 | GTTACGTGACACGTCCGCATgttttAGAGCTAGAAATAGCAAGTTAAAATAAGGC |
| 239B | psg2171_rpsI_B | 9624 | ACTGTGATGTTACCTTCACCgttttAGAGCTAGAAATAGCAAGTTAAAATAAGGC |
| 240B | psg2177_rplQ_B | 9625 | GTTGAGAAATTAATCACTTTgttttAGAGCTAGAAATAGCAAGTTAAAATAAGGC |
| 241B | psg0386_xpt_B | 10049 | TGCGATACCGGAAGCTTCAAgttttAGAGCTAGAAATAGCAAGTTAAAATAAGGC |

| | | | |
|---|---|---|---|
| 245B | psg1072_mraZ_B | 10057 | ATTTAAGTCATAACGAAACTgttttAGAGCTAGAAATAGCAAGTTAAAATAAGGC |
| 249B | psg1158_rimP_B | 10065 | TAAGACATTGAAAAGAAATAgttttAGAGCTAGAAATAGCAAGTTAAAATAAGGC |
| 254B | psg2029_B | 10075 | CGTATTTTAAGTTAATCGATgttttAGAGCTAGAAATAGCAAGTTAAAATAAGGC |
| 256B | psg0014_gdpP_B | 10079 | AGTCCTACAATATGTGTCGTgttttAGAGCTAGAAATAGCAAGTTAAAATAAGGC |

### Supplementary Table 3. Lisbon CRISPRi Mutant Library strain list

| Name | Description |
|---|---|
| LCML1 | JE2 *Δspa:P*$_{xyl/tetO3}$*-dcas9*$_{Spy}$ containing psg0001 (*dnaA*) |
| LCML2 | JE2 *Δspa:P*$_{xyl/tetO3}$*-dcas9*$_{Spy}$ containingsg0002 (*dnaN*) |
| LCML3 | JE2 *Δspa:P*$_{xyl/tetO3}$*-dcas9*$_{Spy}$ containing psg0003 |
| LCML4 | JE2 *Δspa:P*$_{xyl/tetO3}$*-dcas9*$_{Spy}$ containing psg0009 (*serS*) |
| LCML5 | JE2 *Δspa:P*$_{xyl/tetO3}$*-dcas9*$_{Spy}$ containing psg0016 (*dnaB*) |
| LCML6 | JE2 *Δspa:P*$_{xyl/tetO3}$*-dcas9*$_{Spy}$ containing psg0020 (*walR*) |
| LCML7 | JE2 *Δspa:P*$_{xyl/tetO3}$*-dcas9*$_{Spy}$ containing psg0024 (*walJ*) |
| LCML8 | JE2 *Δspa:P*$_{xyl/tetO3}$*-dcas9*$_{Spy}$ containing psg0089 |
| LCML9 | JE2 *Δspa:P*$_{xyl/tetO3}$*-dcas9*$_{Spy}$ containing psg0248 (*tarF*) |
| LCML10 | JE2 *Δspa:P*$_{xyl/tetO3}$*-dcas9*$_{Spy}$ containing psg0249 (*ispD*) |
| LCML11 | JE2 *Δspa:P*$_{xyl/tetO3}$*-dcas9*$_{Spy}$ containing psg0363 |
| LCML12 | JE2 *Δspa:P*$_{xyl/tetO3}$*-dcas9*$_{Spy}$ containing psg0367 (*ssb*) |
| LCML13 | JE2 *Δspa:P*$_{xyl/tetO3}$*-dcas9*$_{Spy}$ containing psg0388 (*guaB*) |
| LCML14 | JE2 *Δspa:P*$_{xyl/tetO3}$*-dcas9*$_{Spy}$ containing psg0452 (*dnaX*) |
| LCML15 | JE2 *Δspa:P*$_{xyl/tetO3}$*-dcas9*$_{Spy}$ containing psg0453 |
| LCML16 | JE2 *Δspa:P*$_{xyl/tetO3}$*-dcas9*$_{Spy}$ containing psg00459 (*tmk*) |
| LCML17 | JE2 *Δspa:P*$_{xyl/tetO3}$*-dcas9*$_{Spy}$ containing psg0461 (*holB*) |
| LCML18 | JE2 *Δspa:P*$_{xyl/tetO3}$*-dcas9*$_{Spy}$ containing psg0467 (*metS*) |
| LCML19 | JE2 *Δspa:P*$_{xyl/tetO3}$*-dcas9*$_{Spy}$ containing psg0477 (*glmU*) |
| LCML20 | JE2 *Δspa:P*$_{xyl/tetO3}$*-dcas9*$_{Spy}$ containing psg0478 (*prs*) |
| LCML21 | JE2 *Δspa:P*$_{xyl/tetO3}$*-dcas9*$_{Spy}$ containing psg0480 (*pth*) |
| LCML22 | JE2 *Δspa:P*$_{xyl/tetO3}$*-dcas9*$_{Spy}$ containing psg0484 |
| LCML23 | JE2 *Δspa:P*$_{xyl/tetO3}$*-dcas9*$_{Spy}$ containing psg0485 (*divIC*) |
| LCML24 | JE2 *Δspa:P*$_{xyl/tetO3}$*-dcas9*$_{Spy}$ containing psg0487 (*tilS*) |
| LCML25 | JE2 *Δspa:P*$_{xyl/tetO3}$*-dcas9*$_{Spy}$ containing psg0492 (*folP*) |
| LCML26 | JE2 *Δspa:P*$_{xyl/tetO3}$*-dcas9*$_{Spy}$ containing psg0496 (*lysS*) |
| LCML27 | JE2 *Δspa:P*$_{xyl/tetO3}$*-dcas9*$_{Spy}$ containing psg0513 (*gltX*) |
| LCML28 | JE2 *Δspa:P*$_{xyl/tetO3}$*-dcas9*$_{Spy}$ containing psg0514 (*cysE*) |
| LCML29 | JE2 *Δspa:P*$_{xyl/tetO3}$*-dcas9*$_{Spy}$ containing psg0532 (*fusA*) |
| LCML30 | JE2 *Δspa:P*$_{xyl/tetO3}$*-dcas9*$_{Spy}$ containing psg0533 (*tuf*) |
| LCML31 | JE2 *Δspa:P*$_{xyl/tetO3}$*-dcas9*$_{Spy}$ containing psg0551 (*folE2*) |
| LCML32 | JE2 *Δspa:P*$_{xyl/tetO3}$*-dcas9*$_{Spy}$ containing psg0570 (*eutD*) |
| LCML33 | JE2 *Δspa:P*$_{xyl/tetO3}$*-dcas9*$_{Spy}$ containing psg0572 (*mvaK1*) |
| LCML34 | JE2 *Δspa:P*$_{xyl/tetO3}$*-dcas9*$_{Spy}$ containing psg0596 (*argS*) |
| LCML35 | JE2 *Δspa:P*$_{xyl/tetO3}$*-dcas9*$_{Spy}$ containing psg0623 (*tagA*) |
| LCML36 | JE2 *Δspa:P*$_{xyl/tetO3}$*-dcas9*$_{Spy}$ containing psg0624 (*tagH*) |
| LCML37 | JE2 *Δspa:P*$_{xyl/tetO3}$*-dcas9*$_{Spy}$ containing psg0625 (*tagG*) |
| LCML38 | JE2 *Δspa:P*$_{xyl/tetO3}$*-dcas9*$_{Spy}$ containing psg0626 (*tagB*) |
| LCML39 | JE2 *Δspa:P*$_{xyl/tetO3}$*-dcas9*$_{Spy}$ containing psg0628 (*tagD*) |
| LCML40 | JE2 *Δspa:P*$_{xyl/tetO3}$*-dcas9*$_{Spy}$ containing psg0703 (*ltaS*) |
| LCML41 | JE2 *Δspa:P*$_{xyl/tetO3}$*-dcas9*$_{Spy}$ containing psg0715 (*nrdI*) |

| | |
|---|---|
| LCML42 | JE2 *Δspa:P$_{xyl/tetO3}$-dcas9$_{Spy}$* containing psg0722 (*murB*) |
| LCML43 | JE2 *Δspa:P$_{xyl/tetO3}$-dcas9$_{Spy}$* containing psg0731 (*tagO*) |
| LCML44 | JE2 *Δspa:P$_{xyl/tetO3}$-dcas9$_{Spy}$* containing psg0737 (*secA*) |
| LCML45 | JE2 *Δspa:P$_{xyl/tetO3}$-dcas9$_{Spy}$* containing psg0738 (*prfB*) |
| LCML46 | JE2 *Δspa:P$_{xyl/tetO3}$-dcas9$_{Spy}$* containing psg0743 (*hprK*) |
| LCML47 | JE2 *Δspa:P$_{xyl/tetO3}$-dcas9$_{Spy}$* containing psg0749 |
| LCML48 | JE2 *Δspa:P$_{xyl/tetO3}$-dcas9$_{Spy}$* containing psg0756 (*gapA*) |
| LCML49 | JE2 *Δspa:P$_{xyl/tetO3}$-dcas9$_{Spy}$* containing psg0761 |
| LCML50 | JE2 *Δspa:P$_{xyl/tetO3}$-dcas9$_{Spy}$* containing psg0765 (*smpB*) |
| LCML51 | JE2 *Δspa:P$_{xyl/tetO3}$-dcas9$_{Spy}$* containing psg0818 (*sufC*) |
| LCML52 | JE2 *Δspa:P$_{xyl/tetO3}$-dcas9$_{Spy}$* containing psg0835 (*dltA*) |
| LCML53 | JE2 *Δspa:P$_{xyl/tetO3}$-dcas9$_{Spy}$* containing psg0858 |
| LCML54 | JE2 *Δspa:P$_{xyl/tetO3}$-dcas9$_{Spy}$* containing psg0865 (*pgi*) |
| LCML55 | JE2 *Δspa:P$_{xyl/tetO3}$-dcas9$_{Spy}$* containing psg0868 (*spsB*) |
| LCML56 | JE2 *Δspa:P$_{xyl/tetO3}$-dcas9$_{Spy}$* containing psg0885 (*fabH*) |
| LCML57 | JE2 *Δspa:P$_{xyl/tetO3}$-dcas9$_{Spy}$* containing psg0897 (*trpS*) |
| LCML58 | JE2 *Δspa:P$_{xyl/tetO3}$-dcas9$_{Spy}$* containing psg0898 (*spxA*) |
| LCML59 | JE2 *Δspa:P$_{xyl/tetO3}$-dcas9$_{Spy}$* containing psg0908 (*ppnK*) |
| LCML60 | JE2 *Δspa:P$_{xyl/tetO3}$-dcas9$_{Spy}$* containing psg0912 (*fabI*) |
| LCML61 | JE2 *Δspa:P$_{xyl/tetO3}$-dcas9$_{Spy}$* containing psg0919 (*murE*) |
| LCML62 | JE2 *Δspa:P$_{xyl/tetO3}$-dcas9$_{Spy}$* containing psg0922 |
| LCML63 | JE2 *Δspa:P$_{xyl/tetO3}$-dcas9$_{Spy}$* containing psg0944 (*menA*) |
| LCML64 | JE2 *Δspa:P$_{xyl/tetO3}$-dcas9$_{Spy}$* containing psg0948 (*menB*) |
| LCML65 | JE2 *Δspa:P$_{xyl/tetO3}$-dcas9$_{Spy}$* containing psg0983 (*ptsH*) |
| LCML66 | JE2 *Δspa:P$_{xyl/tetO3}$-dcas9$_{Spy}$* containing psg0989 (*rnjA*) |
| LCML67 | JE2 *Δspa:P$_{xyl/tetO3}$-dcas9$_{Spy}$* containing psg0990 (*rpoY*) |
| LCML68 | JE2 *Δspa:P$_{xyl/tetO3}$-dcas9$_{Spy}$* containing psg0991 (*def*) |
| LCML69 | JE2 *Δspa:P$_{xyl/tetO3}$-dcas9$_{Spy}$* containing psg1013 (*ftsW*) |
| LCML70 | JE2 *Δspa:P$_{xyl/tetO3}$-dcas9$_{Spy}$* containing psg1024 (*coaD*) |
| LCML71 | JE2 *Δspa:P$_{xyl/tetO3}$-dcas9$_{Spy}$* containing psg1026 |
| LCML72 | JE2 *Δspa:P$_{xyl/tetO3}$-dcas9$_{Spy}$* containing psg1037 (*pheS*) |
| LCML73 | JE2 *Δspa:P$_{xyl/tetO3}$-dcas9$_{Spy}$* containing psg1044 (*trxA*) |
| LCML74 | JE2 *Δspa:P$_{xyl/tetO3}$-dcas9$_{Spy}$* containing psg1049 (*murI*) |
| LCML75 | JE2 *Δspa:P$_{xyl/tetO3}$-dcas9$_{Spy}$* containing psg1074 (*ftsL*) |
| LCML76 | JE2 *Δspa:P$_{xyl/tetO3}$-dcas9$_{Spy}$* containing psg1075 (*pbpA*) |
| LCML77 | JE2 *Δspa:P$_{xyl/tetO3}$-dcas9$_{Spy}$* containing psg1076 (*mraY*) |
| LCML78 | JE2 *Δspa:P$_{xyl/tetO3}$-dcas9$_{Spy}$* containing psg1077 (*murD*) |
| LCML79 | JE2 *Δspa:P$_{xyl/tetO3}$-dcas9$_{Spy}$* containing psg1078 (*divIB*) |
| LCML80 | JE2 *Δspa:P$_{xyl/tetO3}$-dcas9$_{Spy}$* containing psg1079 (*ftsA*) |
| LCML81 | JE2 *Δspa:P$_{xyl/tetO3}$-dcas9$_{Spy}$* containing psg1082 |
| LCML82 | JE2 *Δspa:P$_{xyl/tetO3}$-dcas9$_{Spy}$* containing psg1083 (*sepF*) |
| LCML83 | JE2 *Δspa:P$_{xyl/tetO3}$-dcas9$_{Spy}$* containing psg1087 (*ileS*) |
| LCML84 | JE2 *Δspa:P$_{xyl/tetO3}$-dcas9$_{Spy}$* containing psg1102 (*gmk*) |
| LCML85 | JE2 *Δspa:P$_{xyl/tetO3}$-dcas9$_{Spy}$* containing psg1104 (*coaBC*) |
| LCML86 | JE2 *Δspa:P$_{xyl/tetO3}$-dcas9$_{Spy}$* containing psg1105 (*priA*) |

| | |
|---|---|
| LCML87 | JE2 *Δspa:P*$_{xyl/tetO3}$-*dcas9*$_{Spy}$ containing psg1109 (*fmt*) |
| LCML88 | JE2 *Δspa:P*$_{xyl/tetO3}$-*dcas9*$_{Spy}$ containing psg1115 (*rpe*) |
| LCML89 | JE2 *Δspa:P*$_{xyl/tetO3}$-*dcas9*$_{Spy}$ containing psg1122 (*plsX*) |
| LCML90 | JE2 *Δspa:P*$_{xyl/tetO3}$-*dcas9*$_{Spy}$ containing psg1125 (*acpP*) |
| LCML91 | JE2 *Δspa:P*$_{xyl/tetO3}$-*dcas9*$_{Spy}$ containing psg1128 (*ftsY*) |
| LCML92 | JE2 *Δspa:P*$_{xyl/tetO3}$-*dcas9*$_{Spy}$ containing psg1133 (*trmD*) |
| LCML93 | JE2 *Δspa:P*$_{xyl/tetO3}$-*dcas9*$_{Spy}$ containing psg1136 (*rbgA*) |
| LCML94 | JE2 *Δspa:P*$_{xyl/tetO3}$-*dcas9*$_{Spy}$ containing psg1143 (*topA*) |
| LCML95 | JE2 *Δspa:P*$_{xyl/tetO3}$-*dcas9*$_{Spy}$ containing psg1150 (*tsf*) |
| LCML96 | JE2 *Δspa:P*$_{xyl/tetO3}$-*dcas9*$_{Spy}$ containing psg1151 (*pyrH*) |
| LCML97 | JE2 *Δspa:P*$_{xyl/tetO3}$-*dcas9*$_{Spy}$ containing psg1152 (*frr*) |
| LCML98 | JE2 *Δspa:P*$_{xyl/tetO3}$-*dcas9*$_{Spy}$ containing psg1153 (*uppS*) |
| LCML99 | JE2 *Δspa:P*$_{xyl/tetO3}$-*dcas9*$_{Spy}$ containing psg1156 (*proS*) |
| LCML100 | JE2 *Δspa:P*$_{xyl/tetO3}$-*dcas9*$_{Spy}$ containing psg1157 (*polC*) |
| LCML101 | JE2 *Δspa:P*$_{xyl/tetO3}$-*dcas9*$_{Spy}$ containing psg1159 (*nusA*) |
| LCML102 | JE2 *Δspa:P*$_{xyl/tetO3}$-*dcas9*$_{Spy}$ containing psg1168 (*rnjB*) |
| LCML103 | JE2 *Δspa:P*$_{xyl/tetO3}$-*dcas9*$_{Spy}$ containing psg1176 (*pgsA*) |
| LCML104 | JE2 *Δspa:P*$_{xyl/tetO3}$-*dcas9*$_{Spy}$ containing psg1200 (*glnR*) |
| LCML105 | JE2 *Δspa:P*$_{xyl/tetO3}$-*dcas9*$_{Spy}$ containing psg1237 (*lexA*) |
| LCML106 | JE2 *Δspa:P*$_{xyl/tetO3}$-*dcas9*$_{Spy}$ containing psg1239 (*tkt*) |
| LCML107 | JE2 *Δspa:P*$_{xyl/tetO3}$-*dcas9*$_{Spy}$ containing psg1249 (*plsY*) |
| LCML108 | JE2 *Δspa:P*$_{xyl/tetO3}$-*dcas9*$_{Spy}$ containing psg1250 (*parE*) |
| LCML109 | JE2 *Δspa:P*$_{xyl/tetO3}$-*dcas9*$_{Spy}$ containing psg1257 (*msrR*) |
| LCML110 | JE2 *Δspa:P*$_{xyl/tetO3}$-*dcas9*$_{Spy}$ containing psg1269 (*femA*) |
| LCML111 | JE2 *Δspa:P*$_{xyl/tetO3}$-*dcas9*$_{Spy}$ containing psg1270 (*femB*) |
| LCML112 | JE2 *Δspa:P*$_{xyl/tetO3}$-*dcas9*$_{Spy}$ containing psg1311 (*murG*) |
| LCML113 | JE2 *Δspa:P*$_{xyl/tetO3}$-*dcas9*$_{Spy}$ containing psg1319 (*folA*) |
| LCML114 | JE2 *Δspa:P*$_{xyl/tetO3}$-*dcas9*$_{Spy}$ containing psg1320 (*thyA*) |
| LCML115 | JE2 *Δspa:P*$_{xyl/tetO3}$-*dcas9*$_{Spy}$ containing psg1340 (*recU*) |
| LCML116 | JE2 *Δspa:P*$_{xyl/tetO3}$-*dcas9*$_{Spy}$ containing psg1341 (*pbp2*) |
| LCML117 | JE2 *Δspa:P*$_{xyl/tetO3}$-*dcas9*$_{Spy}$ containing psg1344 (*dnaD*) |
| LCML118 | JE2 *Δspa:P*$_{xyl/tetO3}$-*dcas9*$_{Spy}$ containing psg1347 (*birA*) |
| LCML119 | JE2 *Δspa:P*$_{xyl/tetO3}$-*dcas9*$_{Spy}$ containing psg1348 (*papS*) |
| LCML120 | JE2 *Δspa:P*$_{xyl/tetO3}$-*dcas9*$_{Spy}$ containing psg1351 |
| LCML121 | JE2 *Δspa:P*$_{xyl/tetO3}$-*dcas9*$_{Spy}$ containing psg1360 (*ubiE*) |
| LCML122 | JE2 *Δspa:P*$_{xyl/tetO3}$-*dcas9*$_{Spy}$ containing psg1362 (*hup*) |
| LCML123 | JE2 *Δspa:P*$_{xyl/tetO3}$-*dcas9*$_{Spy}$ containing psg1364 (*engA*) |
| LCML124 | JE2 *Δspa:P*$_{xyl/tetO3}$-*dcas9*$_{Spy}$ containing psg1367 (*cmk*) |
| LCML125 | JE2 *Δspa:P*$_{xyl/tetO3}$-*dcas9*$_{Spy}$ containing psg1373 (*fer*) |
| LCML126 | JE2 *Δspa:P*$_{xyl/tetO3}$-*dcas9*$_{Spy}$ containing psg1453 (*rnz*) |
| LCML127 | JE2 *Δspa:P*$_{xyl/tetO3}$-*dcas9*$_{Spy}$ containing psg1454 (*zwf*) |
| LCML128 | JE2 *Δspa:P*$_{xyl/tetO3}$-*dcas9*$_{Spy}$ containing psg1459 (*gnd*) |
| LCML129 | JE2 *Δspa:P*$_{xyl/tetO3}$-*dcas9*$_{Spy}$ containing psg1464 (*bmfBB*) |
| LCML130 | JE2 *Δspa:P*$_{xyl/tetO3}$-*dcas9*$_{Spy}$ containing psg1466 (*bmfBAA*) |
| LCML131 | JE2 *Δspa:P*$_{xyl/tetO3}$-*dcas9*$_{Spy}$ containing psg1475 (*accC*) |

| | |
|---|---|
| LCML132 | JE2 *Δspa:*$P_{xyl/tetO3}$-*dcas9*$_{Spy}$ containing psg1490 (*efp*) |
| LCML133 | JE2 *Δspa:*$P_{xyl/tetO3}$-*dcas9*$_{Spy}$ containing psg1492 |
| LCML134 | JE2 *Δspa:*$P_{xyl/tetO3}$-*dcas9*$_{Spy}$ containing psg1520 (*trmK*) |
| LCML135 | JE2 *Δspa:*$P_{xyl/tetO3}$-*dcas9*$_{Spy}$ containing psg1521(*rpoD*) |
| LCML136 | JE2 *Δspa:*$P_{xyl/tetO3}$-*dcas9*$_{Spy}$ containing psg1522 (*dnaG*) |
| LCML137 | JE2 *Δspa:*$P_{xyl/tetO3}$-*dcas9*$_{Spy}$ containing psg1525 (*glyS*) |
| LCML138 | JE2 *Δspa:*$P_{xyl/tetO3}$-*dcas9*$_{Spy}$ containing psg1530 (*ybeY*) |
| LCML139 | JE2 *Δspa:*$P_{xyl/tetO3}$-*dcas9*$_{Spy}$ containing psg1540 (*dnaK*) |
| LCML140 | JE2 *Δspa:*$P_{xyl/tetO3}$-*dcas9*$_{Spy}$ containing psg1541 (*grpE*) |
| LCML141 | JE2 *Δspa:*$P_{xyl/tetO3}$-*dcas9*$_{Spy}$ containing psg1546 (*holA*) |
| LCML142 | JE2 *Δspa:*$P_{xyl/tetO3}$-*dcas9*$_{Spy}$ containing psg1557 |
| LCML143 | JE2 *Δspa:*$P_{xyl/tetO3}$-*dcas9*$_{Spy}$ containing psg1567 (*greA*) |
| LCML144 | JE2 *Δspa:*$P_{xyl/tetO3}$-*dcas9*$_{Spy}$ containing psg1575 (*alaS*) |
| LCML145 | JE2 *Δspa:*$P_{xyl/tetO3}$-*dcas9*$_{Spy}$ containing psg1579 (*iscS*) |
| LCML146 | JE2 *Δspa:*$P_{xyl/tetO3}$-*dcas9*$_{Spy}$ containing psg1587 (*hisS*) |
| LCML147 | JE2 *Δspa:*$P_{xyl/tetO3}$-*dcas9*$_{Spy}$ containing psg1589 (*dtd*) |
| LCML148 | JE2 *Δspa:*$P_{xyl/tetO3}$-*dcas9*$_{Spy}$ containing psg1593 (*secF*) |
| LCML149 | JE2 *Δspa:*$P_{xyl/tetO3}$-*dcas9*$_{Spy}$ containing psg1600 (*obgE*) |
| LCML150 | JE2 *Δspa:*$P_{xyl/tetO3}$-*dcas9*$_{Spy}$ containing psg1602 |
| LCML151 | JE2 *Δspa:*$P_{xyl/tetO3}$-*dcas9*$_{Spy}$ containing psg1611 (*valS*) |
| LCML152 | JE2 *Δspa:*$P_{xyl/tetO3}$-*dcas9*$_{Spy}$ containing psg1619 (*hemA*) |
| LCML153 | JE2 *Δspa:*$P_{xyl/tetO3}$-*dcas9*$_{Spy}$ containing psg1620 (*engB*) |
| LCML154 | JE2 *Δspa:*$P_{xyl/tetO3}$-*dcas9*$_{Spy}$ containing psg1624 |
| LCML155 | JE2 *Δspa:*$P_{xyl/tetO3}$-*dcas9*$_{Spy}$ containing psg1627 (*infC*) |
| LCML156 | JE2 *Δspa:*$P_{xyl/tetO3}$-*dcas9*$_{Spy}$ containing psg1629 (*thrS*) |
| LCML157 | JE2 *Δspa:*$P_{xyl/tetO3}$-*dcas9*$_{Spy}$ containing psg1631 (*dnaB*) |
| LCML158 | JE2 *Δspa:*$P_{xyl/tetO3}$-*dcas9*$_{Spy}$ containing psg1632 (*nrdR*) |
| LCML159 | JE2 *Δspa:*$P_{xyl/tetO3}$-*dcas9*$_{Spy}$ containing psg1634 (*coaE*) |
| LCML160 | JE2 *Δspa:*$P_{xyl/tetO3}$-*dcas9*$_{Spy}$ containing psg1645 (*pfkA*) |
| LCML161 | JE2 *Δspa:*$P_{xyl/tetO3}$-*dcas9*$_{Spy}$ containing psg1647 (*accD*) |
| LCML162 | JE2 *Δspa:*$P_{xyl/tetO3}$-*dcas9*$_{Spy}$ containing psg1649 (*dnaE*) |
| LCML163 | JE2 *Δspa:*$P_{xyl/tetO3}$-*dcas9*$_{Spy}$ containing psg1657 (*ackA*) |
| LCML164 | JE2 *Δspa:*$P_{xyl/tetO3}$-*dcas9*$_{Spy}$ containing psg1673 (*plsC*) |
| LCML165 | JE2 *Δspa:*$P_{xyl/tetO3}$-*dcas9*$_{Spy}$ containing psg1675 (*tyrS*) |
| LCML166 | JE2 *Δspa:*$P_{xyl/tetO3}$-*dcas9*$_{Spy}$ containing psg1686 (*murC*) |
| LCML167 | JE2 *Δspa:*$P_{xyl/tetO3}$-*dcas9*$_{Spy}$ containing psg1688 |
| LCML168 | JE2 *Δspa:*$P_{xyl/tetO3}$-*dcas9*$_{Spy}$ containing psg1704 (*leuS*) |
| LCML169 | JE2 *Δspa:*$P_{xyl/tetO3}$-*dcas9*$_{Spy}$ containing psg1729 |
| LCML170 | JE2 *Δspa:*$P_{xyl/tetO3}$-*dcas9*$_{Spy}$ containing psg1730 (*metK*) |
| LCML171 | JE2 *Δspa:*$P_{xyl/tetO3}$-*dcas9*$_{Spy}$ containing psg1748 |
| LCML172 | JE2 *Δspa:*$P_{xyl/tetO3}$-*dcas9*$_{Spy}$ containing psg1749 |
| LCML173 | JE2 *Δspa:*$P_{xyl/tetO3}$-*dcas9*$_{Spy}$ containing psg1869 (*map*) |
| LCML174 | JE2 *Δspa:*$P_{xyl/tetO3}$-*dcas9*$_{Spy}$ containing psg1873 (*murT*) |
| LCML175 | JE2 *Δspa:*$P_{xyl/tetO3}$-*dcas9*$_{Spy}$ containing psg1879 (*dgkB*) |
| LCML176 | JE2 *Δspa:*$P_{xyl/tetO3}$-*dcas9*$_{Spy}$ containing psg1881 (*gatA*) |

| | |
|---|---|
| LCML177 | JE2 $\Delta spa{:}P_{xyl/tetO3}\text{-}dcas9_{Spy}$ containing psg1882 (*gatC*) |
| LCML178 | JE2 $\Delta spa{:}P_{xyl/tetO3}\text{-}dcas9_{Spy}$ containing psg1885 (*ligA*) |
| LCML179 | JE2 $\Delta spa{:}P_{xyl/tetO3}\text{-}dcas9_{Spy}$ containing psg1886 (*pcrA*) |
| LCML180 | JE2 $\Delta spa{:}P_{xyl/tetO3}\text{-}dcas9_{Spy}$ containing psg1894 (*pncB*) |
| LCML181 | JE2 $\Delta spa{:}P_{xyl/tetO3}\text{-}dcas9_{Spy}$ containing psg1900 (*ppaC*) |
| LCML182 | JE2 $\Delta spa{:}P_{xyl/tetO3}\text{-}dcas9_{Spy}$ containing psg1911 (*pmtC*) |
| LCML183 | JE2 $\Delta spa{:}P_{xyl/tetO3}\text{-}dcas9_{Spy}$ containing psg1913 (*pmtA*) |
| LCML184 | JE2 $\Delta spa{:}P_{xyl/tetO3}\text{-}dcas9_{Spy}$ containing psg1983 (*groES*) |
| LCML185 | JE2 $\Delta spa{:}P_{xyl/tetO3}\text{-}dcas9_{Spy}$ containing psg1988 (*hld*) |
| LCML186 | JE2 $\Delta spa{:}P_{xyl/tetO3}\text{-}dcas9_{Spy}$ containing psg2002 (*gcp*) |
| LCML187 | JE2 $\Delta spa{:}P_{xyl/tetO3}\text{-}dcas9_{Spy}$ containing psg2005 (*tsaE*) |
| LCML188 | JE2 $\Delta spa{:}P_{xyl/tetO3}\text{-}dcas9_{Spy}$ containing psg2028 (*acpS*) |
| LCML189 | JE2 $\Delta spa{:}P_{xyl/tetO3}\text{-}dcas9_{Spy}$ containing psg2039 (*ddl*) |
| LCML190 | JE2 $\Delta spa{:}P_{xyl/tetO3}\text{-}dcas9_{Spy}$ containing psg2046 (*oxaA*) |
| LCML191 | JE2 $\Delta spa{:}P_{xyl/tetO3}\text{-}dcas9_{Spy}$ containing psg2055 (*murA*) |
| LCML192 | JE2 $\Delta spa{:}P_{xyl/tetO3}\text{-}dcas9_{Spy}$ containing psg2062 (*atpF*) |
| LCML193 | JE2 $\Delta spa{:}P_{xyl/tetO3}\text{-}dcas9_{Spy}$ containing psg2072 (*prfA*) |
| LCML194 | JE2 $\Delta spa{:}P_{xyl/tetO3}\text{-}dcas9_{Spy}$ containing psg2079 (*fbaA*) |
| LCML195 | JE2 $\Delta spa{:}P_{xyl/tetO3}\text{-}dcas9_{Spy}$ containing psg2081 (*pyrG*) |
| LCML196 | JE2 $\Delta spa{:}P_{xyl/tetO3}\text{-}dcas9_{Spy}$ containing psg2084 (*coaA*) |
| LCML197 | JE2 $\Delta spa{:}P_{xyl/tetO3}\text{-}dcas9_{Spy}$ containing psg2104 (*glmS*) |
| LCML198 | JE2 $\Delta spa{:}P_{xyl/tetO3}\text{-}dcas9_{Spy}$ containing psg2113 (*dacA*) |
| LCML199 | JE2 $\Delta spa{:}P_{xyl/tetO3}\text{-}dcas9_{Spy}$ containing psg2182 (*infA*) |
| LCML200 | JE2 $\Delta spa{:}P_{xyl/tetO3}\text{-}dcas9_{Spy}$ containing psg2184 (*secY*) |
| LCML201 | JE2 $\Delta spa{:}P_{xyl/tetO3}\text{-}dcas9_{Spy}$ containing psg2214 (*femX*) |
| LCML202 | JE2 $\Delta spa{:}P_{xyl/tetO3}\text{-}dcas9_{Spy}$ containing psg2238 (*ureA*) |
| LCML203 | JE2 $\Delta spa{:}P_{xyl/tetO3}\text{-}dcas9_{Spy}$ containing psg2283 (*rpiA*) |
| LCML204 | JE2 $\Delta spa{:}P_{xyl/tetO3}\text{-}dcas9_{Spy}$ containing psg2292 (*fni*) |
| LCML205 | JE2 $\Delta spa{:}P_{xyl/tetO3}\text{-}dcas9_{Spy}$ containing psg2483 (*mvaA*) |
| LCML206 | JE2 $\Delta spa{:}P_{xyl/tetO3}\text{-}dcas9_{Spy}$ containing psg2646 (*trmE*) |
| LCML207 | JE2 $\Delta spa{:}P_{xyl/tetO3}\text{-}dcas9_{Spy}$ containing psg2484 (*mvaS*) |
| LCML208 | JE2 $\Delta spa{:}P_{xyl/tetO3}\text{-}dcas9_{Spy}$ containing psg0005 (*gyrB*) |
| LCML209 | JE2 $\Delta spa{:}P_{xyl/tetO33}\text{-}dcas9_{Spy}$ containing psg0527 (*rpoB*) |
| LCML210 | JE2 $\Delta spa{:}P_{xyl/tetO3}\text{-}dcas9_{Spy}$ containing psg0425 (*mpsA*) |
| LCML211 | JE2 $\Delta spa{:}P_{xyl/tetO3}\text{-}dcas9_{Spy}$ containing psg0639 |
| LCML212 | JE2 $\Delta spa{:}P_{xyl/tetO3}\text{-}dcas9_{Spy}$ containing psg1359 |
| LCML213 | JE2 $\Delta spa{:}P_{xyl/tetO3}\text{-}dcas9_{Spy}$ containing psg1590 (*relA*) |
| LCML214 | JE2 $\Delta spa{:}P_{xyl/tetO3}\text{-}dcas9_{Spy}$ containing psg1791 (*cbf1*) |
| LCML215 | JE2 $\Delta spa{:}P_{xyl/tetO3}\text{-}dcas9_{Spy}$ containing psg2183 (*adk*) |
| LCML216 | JE2 $\Delta spa{:}P_{xyl/tetO3}\text{-}dcas9_{Spy}$ containing psg2185 (*rplO*) |
| LCML217 | JE2 $\Delta spa{:}P_{xyl/tetO3}\text{-}dcas9_{Spy}$ containing psg2647 (*rnpA*) |
| LCML218 | JE2 $\Delta spa{:}P_{xyl/tetO3}\text{-}dcas9_{Spy}$ containing psg2648 (*rpmH*) |
| LCML219 | JE2 $\Delta spa{:}P_{xyl/tetO3}\text{-}dcas9_{Spy}$ containing psg0015 (*rplI*) |
| LCML220 | JE2 $\Delta spa{:}P_{xyl/tetO3}\text{-}dcas9_{Spy}$ containing psg0366 (*rpsF*) |
| LCML221 | JE2 $\Delta spa{:}P_{xyl/tetO3}\text{-}dcas9_{Spy}$ containing psg0479 (*rplY*) |

| | |
|---|---|
| LCML222 | JE2 *Δspa:*$P_{xyl/tetO3}$-*dcas9*$_{Spy}$ containing psg0522 (*rplK*) |
| LCML223 | JE2 *Δspa:*$P_{xyl/tetO3}$-*dcas9*$_{Spy}$ containing psg0524 (*rplJ*) |
| LCML224 | JE2 *Δspa:*$P_{xyl/tetO3}$-*dcas9*$_{Spy}$ containing psg0530 (*rpsL*) |
| LCML225 | JE2 *Δspa:*$P_{xyl/tetO3}$-*dcas9*$_{Spy}$ containing psg0990 (*rpoY*) |
| LCML226 | JE2 *Δspa:*$P_{xyl/tetO3}$-*dcas9*$_{Spy}$ containing psg1027 (*rpmF*) |
| LCML227 | JE2 *Δspa:*$P_{xyl/tetO3}$-*dcas9*$_{Spy}$ containing psg1117 (*rpmB*) |
| LCML228 | JE2 *Δspa:*$P_{xyl/tetO3}$-*dcas9*$_{Spy}$ containing psg1131 (*rpsP*) |
| LCML229 | JE2 *Δspa:*$P_{xyl/tetO3}$-*dcas9*$_{Spy}$ containing psg1134 (*rplS*) |
| LCML230 | JE2 *Δspa:*$P_{xyl/tetO3}$-*dcas9*$_{Spy}$ containing psg1149 (*rpsB*) |
| LCML231 | JE2 *Δspa:*$P_{xyl/tetO3}$-*dcas9*$_{Spy}$ containing psg1166 (*rpsO*) |
| LCML232 | JE2 *Δspa:*$P_{xyl/tetO3}$-*dcas9*$_{Spy}$ containing psg1511 (*rpmG*) |
| LCML233 | JE2 *Δspa:*$P_{xyl/tetO3}$-*dcas9*$_{Spy}$ containing psg1535 (*rpsU*) |
| LCML234 | JE2 *Δspa:*$P_{xyl/tetO3}$-*dcas9*$_{Spy}$ containing psg1545 (*rpsT*) |
| LCML235 | JE2 *Δspa:*$P_{xyl/tetO3}$-*dcas9*$_{Spy}$ containing psg1601 (*rpmA*) |
| LCML236 | JE2 *Δspa:*$P_{xyl/tetO3}$-*dcas9*$_{Spy}$ containing psg1625 (*rplT*) |
| LCML237 | JE2 *Δspa:*$P_{xyl/tetO3}$-*dcas9*$_{Spy}$ containing psg1666 (*rpsD*) |
| LCML238 | JE2 *Δspa:*$P_{xyl/tetO3}$-*dcas9*$_{Spy}$ containing psg2074 (*rpmE2*) |
| LCML239 | JE2 *Δspa:*$P_{xyl/tetO3}$-*dcas9*$_{Spy}$ containing psg2171 (*rpsI*) |
| LCML240 | JE2 *Δspa:*$P_{xyl/tetO3}$-*dcas9*$_{Spy}$ containing psg2177 (*rplQ*) |
| LCML241 | JE2 *Δspa:*$P_{xyl/tetO3}$-*dcas9*$_{Spy}$ containing psg0386 (*xpt*) |
| LCML242 | JE2 *Δspa:*$P_{xyl/tetO3}$-*dcas9*$_{Spy}$ containing psg0529 (*rplGB*) |
| LCML243 | JE2 *Δspa:*$P_{xyl/tetO3}$-*dcas9*$_{Spy}$ containing psg0755 (*gapR*) |
| LCML244 | JE2 *Δspa:*$P_{xyl/tetO3}$-*dcas9*$_{Spy}$ containing psg0906 |
| LCML245 | JE2 *Δspa:*$P_{xyl/tetO3}$-*dcas9*$_{Spy}$ containing psg1072 (*mraZ*) |
| LCML246 | JE2 *Δspa:*$P_{xyl/tetO3}$-*dcas9*$_{Spy}$ containing psg1108 (*def*) |
| LCML247 | JE2 *Δspa:*$P_{xyl/tetO3}$-*dcas9*$_{Spy}$ containing psg1121 (*fapR*) |
| LCML248 | JE2 *Δspa:*$P_{xyl/tetO3}$-*dcas9*$_{Spy}$ containing psg1155 (*rasP*) |
| LCML249 | JE2 *Δspa:*$P_{xyl/tetO3}$-*dcas9*$_{Spy}$ containing psg1158 (*rimP*) |
| LCML250 | JE2 *Δspa:*$P_{xyl/tetO3}$-*dcas9*$_{Spy}$ containing psg1558 (*mtnM*) |
| LCML251 | JE2 *Δspa:*$P_{xyl/tetO3}$-*dcas9*$_{Spy}$ containing psg1635 (*mutM*) |
| LCML252 | JE2 *Δspa:*$P_{xyl/tetO3}$-*dcas9*$_{Spy}$ containing psg1690 |
| LCML253 | JE2 *Δspa:*$P_{xyl/tetO3}$-*dcas9*$_{Spy}$ containing psg1914 (*pmtR*) |
| LCML254 | JE2 *Δspa:*$P_{xyl/tetO3}$-*dcas9*$_{Spy}$ containing psg2029 |
| LCML255 | JE2 *Δspa:*$P_{xyl/tetO3}$-*dcas9*$_{Spy}$ containing psg2205 (*rpsJ*) |
| LCML256 | JE2 *Δspa:*$P_{xyl/tetO3}$-*dcas9*$_{Spy}$ containing psg0014 (*gdpP*) |
| LCML257 | JE2 *Δspa:*$P_{xyl/tetO3}$-*dcas9*$_{Spy}$ containing psg1603 (*rplU*) |
| LCML258 | JE2 *Δspa:*$P_{xyl/tetO3}$-*dcas9*$_{Spy}$ containing psg2172 (*rplM*) |
| LCML259 | JE2 *Δspa:*$P_{xyl/tetO3}$-*dcas9*$_{Spy}$ containing psg1476 (*accB*) |
| LCML260 | JE2 *Δspa:*$P_{xyl/tetO3}$-*dcas9*$_{Spy}$ containing psg1700 (*murJ*) |
| LCML261 | JE2 *Δspa:*$P_{xyl/tetO3}$-*dcas9*$_{Spy}$ containing psg1080 (*ftsZ*) |
| B Clones | |
| LCML4_B | JE2 *Δspa:*$P_{xyl/tetO3}$-*dcas9*$_{Spy}$ containing psg0009_B (*serS*) |
| LCML6_B | JE2 *Δspa:*$P_{xyl/tetO3}$-*dcas9*$_{Spy}$ containing psg0020_B (*walR*) |
| LCML7_B | JE2 *Δspa:*$P_{xyl/tetO3}$-*dcas9*$_{Spy}$ containing psg0024_B (*walJ*) |
| LCML8_B | JE2 *Δspa:*$P_{xyl/tetO3}$-*dcas9*$_{Spy}$ containing psg0089_B |

| | |
|---|---|
| LCML9_B | JE2 *Δspa:P*$_{xyl/tetO3}$-*dcas9*$_{Spy}$ containing psg0248_B (*tarF*) |
| LCML10_B | JE2 *Δspa:P*$_{xyl/tetO3}$-*dcas9*$_{Spy}$ containing psg0249_B (*ispD*) |
| LCML15_B | JE2 *Δspa:P*$_{xyl/tetO3}$-*dcas9*$_{Spy}$ containing psg0453_B |
| LCML24_B | JE2 *Δspa:P*$_{xyl/tetO3}$-*dcas9*$_{Spy}$ containing psg0487_B (*tilS*) |
| LCML25_B | JE2 *Δspa:P*$_{xyl/tetO3}$-*dcas9*$_{Spy}$ containing psg0492_B (*folP*) |
| LCML28_B | JE2 *Δspa:P*$_{xyl/tetO3}$-*dcas9*$_{Spy}$ containing psg0514_B (*cysE*) |
| LCML31_B | JE2 *Δspa:P*$_{xyl/tetO3}$-*dcas9*$_{Spy}$ containing psg0551_B (*folE2*) |
| LCML32_B | JE2 *Δspa:P*$_{xyl/tetO3}$-*dcas9*$_{Spy}$ containing psg0570_B (*eutD*) |
| LCML33_B | JE2 *Δspa:P*$_{xyl/tetO3}$-*dcas9*$_{Spy}$ containing psg0572_B (*mvaK1*) |
| LCML34_B | JE2 *Δspa:P*$_{xyl/tetO3}$-*dcas9*$_{Spy}$ containing psg0596_B (*argS*) |
| LCML35_B | JE2 *Δspa:P*$_{xyl/tetO3}$-*dcas9*$_{Spy}$ containing psg0623_B (*tagA*) |
| LCML36_B | JE2 *Δspa:P*$_{xyl/tetO3}$-*dcas9*$_{Spy}$ containing psg0624_B (*tagH*) |
| LCML40_B | JE2 *Δspa:P*$_{xyl/tetO3}$-*dcas9*$_{Spy}$ containing psg0703_B (*ltaS*) |
| LCML42_B | JE2 *Δspa:P*$_{xyl/tetO3}$-*dcas9*$_{Spy}$ containing psg0722_B (*murB*) |
| LCML46_B | JE2 *Δspa:P*$_{xyl/tetO3}$-*dcas9*$_{Spy}$ containing psg0743_B (*hprK*) |
| LCML55_B | JE2 *Δspa:P*$_{xyl/tetO3}$-*dcas9*$_{Spy}$ containing psg0868_B (*spsB*) |
| LCML56_B | JE2 *Δspa:P*$_{xyl/tetO3}$-*dcas9*$_{Spy}$ containing psg0885_B (*fabH*) |
| LCML62_B | JE2 *Δspa:P*$_{xyl/tetO3}$-*dcas9*$_{Spy}$ containing psg0922_B |
| LCML65_B | JE2 *Δspa:P*$_{xyl/tetO3}$-*dcas9*$_{Spy}$ containing psg0983_B (*ptsH*) |
| LCML66_B | JE2 *Δspa:P*$_{xyl/tetO3}$-*dcas9*$_{Spy}$ containing psg0989_B (*rnjA*) |
| LCML67_B | JE2 *Δspa:P*$_{xyl/tetO3}$-*dcas9*$_{Spy}$ containing psg0990_B (*rpoY*) |
| LCML68_B | JE2 *Δspa:P*$_{xyl/tetO3}$-*dcas9*$_{Spy}$ containing psg0991_B (*def*) |
| LCML70_B | JE2 *Δspa:P*$_{xyl/tetO3}$-*dcas9*$_{Spy}$ containing psg1024_B (*coaD*) |
| LCML92_B | JE2 *Δspa:P*$_{xyl/tetO3}$-*dcas9*$_{Spy}$ containing psg1133_B (*trmD*) |
| LCML97_B | JE2 *Δspa:P*$_{xyl/tetO3}$-*dcas9*$_{Spy}$ containing psg1152_B (*frr*) |
| LCML98_B | JE2 *Δspa:P*$_{xyl/tetO3}$-*dcas9*$_{Spy}$ containing psg1153_B (*uppS*) |
| LCML100_B | JE2 *Δspa:P*$_{xyl/tetO3}$-*dcas9*$_{Spy}$ containing psg1157_B (*polC*) |
| LCML103_B | JE2 *Δspa:P*$_{xyl/tetO3}$-*dcas9*$_{Spy}$ containing psg1176_B (*pgsA)* |
| LCML107_B | JE2 *Δspa:P*$_{xyl/tetO3}$-*dcas9*$_{Spy}$ containing psg1249_B (*plsY*) |
| LCML109_B | JE2 *Δspa:P*$_{xyl/tetO3}$-*dcas9*$_{Spy}$ containing psg1257_B (*msrR*) |
| LCML115_B | JE2 *Δspa:P*$_{xyl/tetO3}$-*dcas9*$_{Spy}$ containing psg1340_B (*recU*) |
| LCML116_B | JE2 *Δspa:P*$_{xyl/tetO3}$-*dcas9*$_{Spy}$ containing psg1341_B (*pbp2*) |
| LCML117_B | JE2 *Δspa:P*$_{xyl/tetO3}$-*dcas9*$_{Spy}$ containing psg1344_B (*dnaD*) |
| LCML118_B | JE2 *Δspa:P*$_{xyl/tetO3}$-*dcas9*$_{Spy}$ containing psg1347_B (*birA*) |
| LCML119_B | JE2 *Δspa:P*$_{xyl/tetO3}$-*dcas9*$_{Spy}$ containing psg1348_B (*papS*) |
| LCML120_B | JE2 *Δspa:P*$_{xyl/tetO3}$-*dcas9*$_{Spy}$ containing psg1351_B |
| LCML121_B | JE2 *Δspa:P*$_{xyl/tetO3}$-*dcas9*$_{Spy}$ containing psg1360_B (*ubiE*) |
| LCML129_B | JE2 *Δspa:P*$_{xyl/tetO3}$-*dcas9*$_{Spy}$ containing psg1464_B (*bmfBB*) |
| LCML131_B | JE2 *Δspa:P*$_{xyl/tetO3}$-*dcas9*$_{Spy}$ containing psg1475_B (*accC*) |
| LCML132_B | JE2 *Δspa:P*$_{xyl/tetO3}$-*dcas9*$_{Spy}$ containing psg1490_B (*efp*) |
| LCML133_B | JE2 *Δspa:P*$_{xyl/tetO3}$-*dcas9*$_{Spy}$ containing psg1492_B |
| LCML134_B | JE2 *Δspa:P*$_{xyl/tetO3}$-*dcas9*$_{Spy}$ containing psg1520_B (*trmK*) |
| LCML136_B | JE2 *Δspa:P*$_{xyl/tetO3}$-*dcas9*$_{Spy}$ containing psg1522_B (*dnaG*) |
| LCML137_B | JE2 *Δspa:P*$_{xyl/tetO3}$-*dcas9*$_{Spy}$ containing psg1525_B (*glyS*) |
| LCML139_B | JE2 *Δspa:P*$_{xyl/tetO3}$-*dcas9*$_{Spy}$ containing psg1540_B (*dnaK*) |

| | |
|---|---|
| LCML140_B | JE2 *Δspa:P$_{xyl/tetO3}$-dcas9$_{Spy}$* containing psg1541_B (*grpE*) |
| LCML141_B | JE2 *Δspa:P$_{xyl/tetO3}$-dcas9$_{Spy}$* containing psg1546_B (*holA*) |
| LCML143_B | JE2 *Δspa:P$_{xyl/tetO3}$-dcas9$_{Spy}$* containing psg1567_B (*greA*) |
| LCML144_B | JE2 *Δspa:P$_{xyl/tetO3}$-dcas9$_{Spy}$* containing psg1575_B (*alaS*) |
| LCML147_B | JE2 *Δspa:P$_{xyl/tetO3}$-dcas9$_{Spy}$* containing psg1589_B (*dtd*) |
| LCML148_B | JE2 *Δspa:P$_{xyl/tetO3}$-dcas9$_{Spy}$* containing psg1593_B (*secF*) |
| LCML151_B | JE2 *Δspa:P$_{xyl/tetO3}$-dcas9$_{Spy}$* containing psg1611_B (*valS*) |
| LCML152_B | JE2 *Δspa:P$_{xyl/tetO3}$-dcas9$_{Spy}$* containing psg1619_B (*hemA*) |
| LCML154_B | JE2 *Δspa:P$_{xyl/tetO3}$-dcas9$_{Spy}$* containing psg1624_B |
| LCML156_B | JE2 *Δspa:P$_{xyl/tetO3}$-dcas9$_{Spy}$* containing psg1629_B (*thrS*) |
| LCML157_B | JE2 *Δspa:P$_{xyl/tetO3}$-dcas9$_{Spy}$* containing psg1631_B (*dnaB*) |
| LCML158_B | JE2 *Δspa:P$_{xyl/tetO3}$-dcas9$_{Spy}$* containing psg1632_B (*nrdR*) |
| LCML159_B | JE2 *Δspa:P$_{xyl/tetO3}$-dcas9$_{Spy}$* containing psg1634_B (*coaE*) |
| LCML161_B | JE2 *Δspa:P$_{xyl/tetO3}$-dcas9$_{Spy}$* containing psg1647_B (*accD*) |
| LCML162_B | JE2 *Δspa:P$_{xyl/tetO3}$-dcas9$_{Spy}$* containing psg1649_B (*dnaE*) |
| LCML163_B | JE2 *Δspa:P$_{xyl/tetO3}$-dcas9$_{Spy}$* containing psg1657_B (*ackA*) |
| LCML165_B | JE2 *Δspa:P$_{xyl/tetO3}$-dcas9$_{Spy}$* containing psg1675_B (*tyrS*) |
| LCML168_B | JE2 *Δspa:P$_{xyl/tetO3}$-dcas9$_{Spy}$* containing psg1704_B (*leuS*) |
| LCML169_B | JE2 *Δspa:P$_{xyl/tetO3}$-dcas9$_{Spy}$* containing psg1729_B |
| LCML170_B | JE2 *Δspa:P$_{xyl/tetO3}$-dcas9$_{Spy}$* containing psg1730_B (*metK*) |
| LCML171_B | JE2 *Δspa:P$_{xyl/tetO3}$-dcas9$_{Spy}$* containing psg1748_B |
| LCML172_B | JE2 *Δspa:P$_{xyl/tetO3}$-dcas9$_{Spy}$* containing psg1749_B |
| LCML173_B | JE2 *Δspa:P$_{xyl/tetO3}$-dcas9$_{Spy}$* containing psg1869_B (*map_*) |
| LCML180_B | JE2 *Δspa:P$_{xyl/tetO3}$-dcas9$_{Spy}$* containing psg1894_B (*pncB*) |
| LCML181_B | JE2 *Δspa:P$_{xyl/tetO3}$-dcas9$_{Spy}$* containing psg1900_B (*ppaC*) |
| LCML182_B | JE2 *Δspa:P$_{xyl/tetO3}$-dcas9$_{Spy}$* containing psg1911_B (*pmtC*) |
| LCML184_B | JE2 *Δspa:P$_{xyl/tetO3}$-dcas9$_{Spy}$* containing psg1983_B (*groES*) |
| LCML185_B | JE2 *Δspa:P$_{xyl/tetO3}$-dcas9$_{Spy}$* containing psg1988_B (*hld*) |
| LCML188_B | JE2 *Δspa:P$_{xyl/tetO3}$-dcas9$_{Spy}$* containing psg2028_B (*acpS*) |
| LCML191_B | JE2 *Δspa:P$_{xyl/tetO3}$-dcas9$_{Spy}$* containing psg2055_B (*murA*) |
| LCML192_B | JE2 *Δspa:P$_{xyl/tetO3}$-dcas9$_{Spy}$* containing psg2062_B (*atpF*) |
| LCML198_B | JE2 *Δspa:P$_{xyl/tetO3}$-dcas9$_{Spy}$* containing psg2113_B (*dacA*) |
| LCML200_B | JE2 *Δspa:P$_{xyl/tetO3}$-dcas9$_{Spy}$* containing psg2184_B (*secY*) |
| LCML202_B | JE2 *Δspa:P$_{xyl/tetO3}$-dcas9$_{Spy}$* containing psg2238_B (*ureA*) |
| LCML203_B | JE2 *Δspa:P$_{xyl/tetO3}$-dcas9$_{Spy}$* containing psg2283_B (*rpiA*) |
| LCML208_B | JE2 *Δspa:P$_{xyl/tetO3}$-dcas9$_{Spy}$* containing psg0005_B (*gyrB*) |
| LCML209_B | JE2 *Δspa:P$_{xyl/tetO3}$-dcas9$_{Spy}$* containing psg0527_B (*rpoB*) |
| LCML211_B | JE2 *Δspa:P$_{xyl/tetO3}$-dcas9$_{Spy}$* containing psg0639_B |
| LCML212_B | JE2 *Δspa:P$_{xyl/tetO3}$-dcas9$_{Spy}$* containing psg1359_B |
| LCML213_B | JE2 *Δspa:P$_{xyl/tetO3}$-dcas9$_{Spy}$* containing psg1590_B (*relA*) |
| LCML214_B | JE2 *Δspa:P$_{xyl/tetO3}$-dcas9$_{Spy}$* containing psg1791_B (*cbf1*) |
| LCML219_B | JE2 *Δspa:P$_{xyl/tetO3}$-dcas9$_{Spy}$* containing psg0015_B (*rplI*) |
| LCML221_B | JE2 *Δspa:P$_{xyl/tetO3}$-dcas9$_{Spy}$* containing psg0479_B (*rplY*) |
| LCML222_B | JE2 *Δspa:P$_{xyl/tetO3}$-dcas9$_{Spy}$* containing psg0522_B (*rplK*) |
| LCML223_B | JE2 *Δspa:P$_{xyl/tetO3}$-dcas9$_{Spy}$* containing psg0524_B (*rplJ*) |

| | |
|---|---|
| LCML225_B | JE2 *Δspa:*$P_{xyl/tetO3}$-*dcas9*$_{Spy}$ containing psg0990_B (*rpoY*) |
| LCML231_B | JE2 *Δspa:*$P_{xyl/tetO3}$-*dcas9*$_{Spy}$ containing psg1166_B (*rpsO*) |
| LCML234_B | JE2 *Δspa:*$P_{xyl/tetO3}$-*dcas9*$_{Spy}$ containing psg1545_B (*rpsT*) |
| LCML235_B | JE2 *Δspa:*$P_{xyl/tetO3}$-*dcas9*$_{Spy}$ containing psg1601_B (*rmpA*) |
| LCML236_B | JE2 *Δspa:*$P_{xyl/tetO3}$-*dcas9*$_{Spy}$ containing psg1625_B (*rplT*) |
| LCML237_B | JE2 *Δspa:*$P_{xyl/tetO3}$-*dcas9*$_{Spy}$ containing psg1666_B (*rpsD*) |
| LCML239_B | JE2 *Δspa:*$P_{xyl/tetO3}$-*dcas9*$_{Spy}$ containing psg2171_B (*rpsI)* |
| LCML240_B | JE2 *Δspa:*$P_{xyl/tetO3}$-*dcas9*$_{Spy}$ containing psg2177_B (*rplQ*) |
| LCML241_B | JE2 *Δspa:*$P_{xyl/tetO3}$-*dcas9*$_{Spy}$ containing psg0386_B (*xpt*) |
| LCML245_B | JE2 *Δspa:*$P_{xyl/tetO3}$-*dcas9*$_{Spy}$ containing psg1072_B (*mraZ*) |
| LCML249_B | JE2 *Δspa:*$P_{xyl/tetO3}$-*dcas9*$_{Spy}$ containing psg1158_B (*rimP*) |
| LCML254_B | JE2 *Δspa:*$P_{xyl/tetO3}$-*dcas9*$_{Spy}$ containing psg2029_B |
| LCML256_B | JE2 *Δspa:*$P_{xyl/tetO3}$-*dcas9*$_{Spy}$ containing psg0014_B (*gdpP*) |

### Supplementary Table 4. Lisbon CRISPRi Mutant Library clone list

| LCML Number | Locus Tag | Name | Best Clone | R Value | Temp | sgRNA Sequence |
|---|---|---|---|---|---|---|
| 1 | SAUSA300_0001 | sg0001_dnaA | A | 0.083 | 30 | ATACACAAAAATAACAGCCGgttttAGAGCTAGAAATAGCAAGTTAAAATAAGGC |
| 2 | SAUSA300_0002 | sg0002_dnaN | A | 0.027 | 30 | TCGTTTATATATAATTATATgttttAGAGCTAGAAATAGCAAGTTAAAATAAGGC |
| 3 | SAUSA300_0003 | sg0003 | A | 0.385 | 30 | ACAAATTTCATTTAAAATAGgttttAGAGCTAGAAATAGCAAGTTAAAATAAGGC |
| 4 | SAUSA300_0009 | sg0009_serS | A | 0.267 | 37 | CAAATAATTATCATTTATTAgttttAGAGCTAGAAATAGCAAGTTAAAATAAGGC |
| 5 | SAUSA300_0016 | sg0016_dnaB | A | 0.261 | 30 | TCATACATTCTATCCATGAAgttttAGAGCTAGAAATAGCAAGTTAAAATAAGGC |
| 6 | SAUSA300_0020 | sg0020_walR | A | 0.469 | 37 | ATGTTTGTGTAAAAAATCACgttttAGAGCTAGAAATAGCAAGTTAAAATAAGGC |
| 7 | SAUSA300_0024 | sg0024_walJ | A | 0.389 | 37 | TGTAACTTTAGTTCATCGACgttttAGAGCTAGAAATAGCAAGTTAAAATAAGGC |
| 8 | SAUSA300_0089 | sg0089 | A | 0.407 | 37 | GCCAAAATAAAAAATGGACGgttttAGAGCTAGAAATAGCAAGTTAAAATAAGGC |
| 9 | SAUSA300_0248 | sg0248_tarF | B | 0.401 | 30 | ATATAATTACAAAAACACGTgttttAGAGCTAGAAATAGCAAGTTAAAATAAGGC |
| 10 | SAUSA300_0249 | sg0249_ispD | B | 0.422 | 30 | TCTAATGTATGGATTAAAATgttttAGAGCTAGAAATAGCAAGTTAAAATAAGGC |
| 11 | SAUSA300_0363 | sg0363 | A | 0.278 | 30 | TGATTATAAGCAGTCATAATgttttAGAGCTAGAAATAGCAAGTTAAAATAAGGC |
| 12 | SAUSA300_0367 | sg0367_ssb | A | 0.111 | 30 | TTTATATTTGCACCTCCTTGgttttAGAGCTAGAAATAGCAAGTTAAAATAAGGC |
| 13 | SAUSA300_0388 | sg0388_guaB | A | 0.177 | 30 | GGTAAAATATCAGATTGTGCgttttAGAGCTAGAAATAGCAAGTTAAAATAAGGC |
| 14 | SAUSA300_0452 | sg0452_dnaX | A | 0.364 | 30 | GGAAATTTTACGATTCCGTGgttttAGAGCTAGAAATAGCAAGTTAAAATAAGGC |
| 15 | SAUSA300_0453 | sg0453 | B | 0.367 | 30 | GTTTGTTGTGCTTGTTGCTTgttttAGAGCTAGAAATAGCAAGTTAAAATAAGGC |
| 16 | SAUSA300_0459 | sg00459_tmk | A | 0.021 | 30 | GTTTCTCCTTTGAAAATAATgttttAGAGCTAGAAATAGCAAGTTAAAATAAGGC |
| 17 | SAUSA300_0461 | sg0461_holB | A | 0.239 | 30 | CCTTTTGCTTTTATATAAAAgttttAGAGCTAGAAATAGCAAGTTAAAATAAGGC |
| 18 | SAUSA300_0467 | sg0467_metS | A | 0.015 | 30 | AAGATTTCTATGCATTTCAAgttttAGAGCTAGAAATAGCAAGTTAAAATAAGGC |
| 19 | SAUSA300_0477 | sg0477_glmU | A | 0.223 | 30 | TTCAGCACCATGTCCTACGAgttttAGAGCTAGAAATAGCAAGTTAAAATAAGGC |
| 20 | SAUSA300_0478 | sg0478_prs | A | 0.476 | 30 | AGATTACATTAATATTACATgttttAGAGCTAGAAATAGCAAGTTAAAATAAGGC |
| 21 | SAUSA300_0480 | sg0480_pth | A | 0.492 | 30 | GACAAATTCATCATTGTCATgttttAGAGCTAGAAATAGCAAGTTAAAATAAGGC |
| 22 | SAUSA300_0484 | sg0484 | B | 0.032 | 30 | TTCTAGCCACTGCTTTAACGgttttAGAGCTAGAAATAGCAAGTTAAAATAAGGC |
| 23 | SAUSA300_0485 | sg0485_divIC | A | 0.294 | 30 | CAATATCATTGCGATGTTTTgttttAGAGCTAGAAATAGCAAGTTAAAATAAGGC |

| LCML Number | Locus Tag | Name | Best Clone | R Value | Temp | sgRNA Sequence |
|---|---|---|---|---|---|---|
| 24 | SAUSA300_0487 | sg0487_tilS | B | 0.083 | 30 | TAGAAACAGCGACAACAATAgttttAGAGCTAGAAATAGCAAGTTAAAATAAGGC |
| 25 | SAUSA300_0492 | sg0492_folP | B | 0.434 | 30 | GTCAGCACCTTCATCTATCAgttttAGAGCTAGAAATAGCAAGTTAAAATAAGGC |
| 26 | SAUSA300_0496 | sg0496_lysS | A | 0.216 | 30 | TAATCCTTTAGTATTCCAACgttttAGAGCTAGAAATAGCAAGTTAAAATAAGGC |
| 27 | SAUSA300_0513 | sg0513_gltX | A | 0.059 | 30 | TGAAGATACCCAGTTGGACTgttttAGAGCTAGAAATAGCAAGTTAAAATAAGGC |
| 28 | SAUSA300_0514 | sg0514_cysE | B | 0.625 | 30 | TCTAATGTTGAACGTGCCGCgttttAGAGCTAGAAATAGCAAGTTAAAATAAGGC |
| 29 | SAUSA300_0532 | sg0532_fusA | A | 0.006 | 30 | CTGAATAAATACGATAGATAgttttAGAGCTAGAAATAGCAAGTTAAAATAAGGC |
| 30 | SAUSA300_0533 | sg0533_tuf | A | 0.008 | 30 | GTGACCGATAGTACCGATATgttttAGAGCTAGAAATAGCAAGTTAAAATAAGGC |
| 31 | SAUSA300_0551 | sg0551_folE2 | B | 0.431 | 30 | GTCATTTCGTTCTTAGTAGTgttttAGAGCTAGAAATAGCAAGTTAAAATAAGGC |
| 32 | SAUSA300_0570 | sg0570_eutD | A | 0.486 | 30 | TAAATCTTATTAATCATTCAgttttAGAGCTAGAAATAGCAAGTTAAAATAAGGC |
| 33 | SAUSA300_0572 | sg0572_mvaK1 | B | 0.501 | 30 | CGAATAGTTCCCGCTCTCTAgttttAGAGCTAGAAATAGCAAGTTAAAATAAGGC |
| 34 | SAUSA300_0596 | sg0596_argS | B | 0.216 | 30 | GGAACTTCAATTTTAATATCgttttAGAGCTAGAAATAGCAAGTTAAAATAAGGC |
| 35 | SAUSA300_0623 | sg0623_tagA | B | 0.342 | 30 | CGTTGATTGATTTGCAAAAAgttttAGAGCTAGAAATAGCAAGTTAAAATAAGGC |
| 36 | SAUSA300_0624 | sg0624_tagH | B | 0.216 | 30 | AAATGTTTTGTTTTTATGTTgttttAGAGCTAGAAATAGCAAGTTAAAATAAGGC |
| 37 | SAUSA300_0625 | sg0625_tagG | A | 0.213 | 30 | AAACCATAATTTGCATAACAgttttAGAGCTAGAAATAGCAAGTTAAAATAAGGC |
| 38 | SAUSA300_0626 | sg0626_tagB | A | 0.347 | 30 | TGCGGTTTATCAATCACTTGgttttAGAGCTAGAAATAGCAAGTTAAAATAAGGC |
| 39 | SAUSA300_0628 | sg0628_tagD | A | 0.369 | 30 | TCCTCATATTGTCACATCATgttttAGAGCTAGAAATAGCAAGTTAAAATAAGGC |
| 40 | SAUSA300_0703 | sg0703_ltaS | A | 0.597 | 30 | CTTCAAGGTAATCGTTATTAgttttAGAGCTAGAAATAGCAAGTTAAAATAAGGC |
| 41 | SAUSA300_0715 | sg0715_nrdI | A | 0.405 | 30 | TAAAAAATGCTTTAACAACAgttttAGAGCTAGAAATAGCAAGTTAAAATAAGGC |
| 42 | SAUSA300_0722 | sg0722_murB | B | 0.157 | 30 | TGATATTAAAGACATGAGAAgttttAGAGCTAGAAATAGCAAGTTAAAATAAGGC |
| 43 | SAUSA300_0731 | sg0731_tagO | A | 0.460 | 30 | TTCGATATTGCAATAACAATgttttAGAGCTAGAAATAGCAAGTTAAAATAAGGC |
| 44 | SAUSA300_0737 | sg0737_secA | A | 0.365 | 30 | CCTTTAGCTAAAAAACTGTTgttttAGAGCTAGAAATAGCAAGTTAAAATAAGGC |
| 45 | SAUSA300_0738 | sg0738_prfB | A | 0.417 | 30 | TTTGGTTATCCCAAAAATTgttttAGAGCTAGAAATAGCAAGTTAAAATAAGGC |
| 46 | SAUSA300_0743 | sg0743_hprK | A | 0.550 | 30 | TATTGCTTGCTTAATTTACAgttttAGAGCTAGAAATAGCAAGTTAAAATAAGGC |
| 47 | SAUSA300_0749 | sg0749 | A | 0.416 | 30 | TTGTCCACCCGTACAACCGAgttttAGAGCTAGAAATAGCAAGTTAAAATAAGGC |
| 48 | SAUSA300_0756 | sg0756_gapA | A | 0.460 | 30 | TTCAAGTATTATCTTTGCTGgttttAGAGCTAGAAATAGCAAGTTAAAATAAGGC |
| 49 | SAUSA300_0761 | sg_0761 | A | 0.393 | 30 | CGTTAAATTAAGTAATGCTTgttttAGAGCTAGAAATAGCAAGTTAAAATAAGGC |

| LCML Number | Locus Tag | Name | Best Clone | R Value | Temp | sgRNA Sequence |
|---|---|---|---|---|---|---|
| 50 | SAUSA300_0765 | sg0765_smpB | A | 0.018 | 30 | CGATTTTCCGCTAATGTACCgttttAGAGCTAGAAATAGCAAGTTAAAATAAGGC |
| 51 | SAUSA300_0818 | sg0818_sufC | A | 0.423 | 30 | TATCCTCAATAGACACATGTgttttAGAGCTAGAAATAGCAAGTTAAAATAAGGC |
| 52 | SAUSA300_0835 | sg0835_dltA | A | 0.426 | 30 | TGTCTAACAGCAATGCTTTGgttttAGAGCTAGAAATAGCAAGTTAAAATAAGGC |
| 53 | SAUSA300_0858 | sg0858 | A | 0.165 | 30 | GTCTCAACAAACGCACCGTAgttttAGAGCTAGAAATAGCAAGTTAAAATAAGGC |
| 54 | SAUSA300_0865 | sg0865_pgi | A | 0.397 | 30 | TTTCAACAAACATTTCAAACgttttAGAGCTAGAAATAGCAAGTTAAAATAAGGC |
| 55 | SAUSA300_0868 | sg0868_spsB | A | 0.684 | 30 | CGCTCGCCATCTTTCAAAGTgttttAGAGCTAGAAATAGCAAGTTAAAATAAGGC |
| 56 | SAUSA300_0885 | sg0885_fabH | A | 0.554 | 30 | GCATTGTCAATAATCTTTTCgttttAGAGCTAGAAATAGCAAGTTAAAATAAGGC |
| 57 | SAUSA300_0897 | sg0897_trpS | A | 0.020 | 30 | TAGTAGGAATTCCACTAGGTgttttAGAGCTAGAAATAGCAAGTTAAAATAAGGC |
| 58 | SAUSA300_0898 | sg0898_spxA | A | 0.104 | 30 | TTACGGCAAGATGTGCAACTgttttAGAGCTAGAAATAGCAAGTTAAAATAAGGC |
| 59 | SAUSA300_0908 | sg0908_ppnK | A | 0.118 | 30 | GTTCATCATTTTATGCTTTAgttttAGAGCTAGAAATAGCAAGTTAAAATAAGGC |
| 60 | SAUSA300_0912 | sg0912_fabI | A | 0.423 | 30 | CCTTATAATAATTAATTTAAgttttAGAGCTAGAAATAGCAAGTTAAAATAAGGC |
| 61 | SAUSA300_0919 | sg0919_murE | A | 0.674 | 30 | ACTATTAAAAATAAAACATAgttttAGAGCTAGAAATAGCAAGTTAAAATAAGGC |
| 62 | SAUSA300_0922 | sg0922 | B | 0.597 | 30 | AAATAAGGTAAGATCAAACTgttttAGAGCTAGAAATAGCAAGTTAAAATAAGGC |
| 63 | SAUSA300_0944 | sg0944_menA | A | 0.470 | 30 | ACGGAAGCAGTTAATGTATGgttttAGAGCTAGAAATAGCAAGTTAAAATAAGGC |
| 64 | SAUSA300_0948 | sg0948_menB | A | 0.187 | 30 | GTAAACGCATTGCGTACTTCgttttAGAGCTAGAAATAGCAAGTTAAAATAAGGC |
| 65 | SAUSA300_0983 | sg0983_ptsH | B | 0.302 | 30 | AAATTCAATTCGTTTAAAACgttttAGAGCTAGAAATAGCAAGTTAAAATAAGGC |
| 66 | SAUSA300_0989 | sg0989_rnjA | B | 0.089 | 30 | ATCCCTAATAAGTTATCATCgttttAGAGCTAGAAATAGCAAGTTAAAATAAGGC |
| 67 | SAUSA300_0990 | sg0990_rpoY | B | 0.760 | 30 | GGATAAAAAGTTTGGTAGAAgttttAGAGCTAGAAATAGCAAGTTAAAATAAGGC |
| 68 | SAUSA300_0991 | sg0991_def | A | 0.910 | 30 | GCTGCTTTTTGACGCAAAGTgttttAGAGCTAGAAATAGCAAGTTAAAATAAGGC |
| 69 | SAUSA300_1013 | sg1013_ftsW | A | 0.075 | 30 | CGGATAATCAATAAACTTTGgttttAGAGCTAGAAATAGCAAGTTAAAATAAGGC |
| 70 | SAUSA300_1024 | sg1024_coaD | A | 0.648 | 37 | GTAATGGGGTCAAAACTACCgttttAGAGCTAGAAATAGCAAGTTAAAATAAGGC |
| 71 | SAUSA300_1026 | sg1026 | A | 0.155 | 30 | ACCGTTTGATCAAATTCAAAgttttAGAGCTAGAAATAGCAAGTTAAAATAAGGC |
| 72 | SAUSA300_1037 | sg1037_pheS | A | 0.032 | 30 | AACGCAGGTTTATCTTCATTgttttAGAGCTAGAAATAGCAAGTTAAAATAAGGC |
| 73 | SAUSA300_1044 | sg1044_trxA | A | 0.319 | 30 | ACCGGAGCGATCATTTTACAgttttAGAGCTAGAAATAGCAAGTTAAAATAAGGC |
| 74 | SAUSA300_1049 | sg1049_murI | A | 0.418 | 30 | CCAGAGTCTATTACACCTATgttttAGAGCTAGAAATAGCAAGTTAAAATAAGGC |
| 75 | SAUSA300_1074 | sg1074_ftsL | A | 0.009 | 30 | AAACTTGTTCGTCATATGGTgttttAGAGCTAGAAATAGCAAGTTAAAATAAGGC |

| LCML Number | Locus Tag | Name | Best Clone | R Value | Temp | sgRNA Sequence |
|---|---|---|---|---|---|---|
| 76 | SAUSA300_1075 | sg1075_pbpA | A | 0.019 | 30 | GTCCGAATAAACCAACAAGTgttttAGAGCTAGAAATAGCAAGTTAAAATAAGGC |
| 77 | SAUSA300_1076 | sg1076_mraY | A | 0.001 | 30 | TTTAATGTAGGTATTAAAACgttttAGAGCTAGAAATAGCAAGTTAAAATAAGGC |
| 78 | SAUSA300_1077 | sg1077_murD | A | 0.656 | 30 | AATAAAAATGTATTAGTTGTgttttAGAGCTAGAAATAGCAAGTTAAAATAAGGC |
| 79 | SAUSA300_1078 | sg1078_divIB | A | 0.095 | 30 | GGAATACATCTAAGAAAAGAgttttAGAGCTAGAAATAGCAAGTTAAAATAAGGC |
| 80 | SAUSA300_1079 | sg1079_ftsA | A | 0.038 | 30 | ATTTTTTATACCGCTCGTGTgttttAGAGCTAGAAATAGCAAGTTAAAATAAGGC |
| 81 | SAUSA300_1082 | sg1082 | A | 0.485 | 37 | TTTGTAACTGCAATCACGTTgttttAGAGCTAGAAATAGCAAGTTAAAATAAGGC |
| 82 | SAUSA300_1083 | sg1083_sepF | A | 0.269 | 30 | TTTACCTGTTGTTGTTTGTCgttttAGAGCTAGAAATAGCAAGTTAAAATAAGGC |
| 83 | SAUSA300_1087 | sg1087_ileS | A | 0.206 | 30 | CGCATTGGGAAATCTGTTTTgttttAGAGCTAGAAATAGCAAGTTAAAATAAGGC |
| 84 | SAUSA300_1102 | sg1102_gmk | A | 0.100 | 30 | GTACCTTTACCTACTCCAGAgttttAGAGCTAGAAATAGCAAGTTAAAATAAGGC |
| 85 | SAUSA300_1104 | sg1104_coaBC | A | 0.200 | 30 | TGCCGCAATGCCACCTGTAAgttttAGAGCTAGAAATAGCAAGTTAAAATAAGGC |
| 86 | SAUSA300_1105 | sg1105_priA | A | 0.181 | 30 | GATGACAGATTCGAGTTGTTgttttAGAGCTAGAAATAGCAAGTTAAAATAAGGC |
| 87 | SAUSA300_1109 | sg1109_fmt | A | 0.235 | 30 | AAAACAGTTGTTGAAAAGTCgttttAGAGCTAGAAATAGCAAGTTAAAATAAGGC |
| 88 | SAUSA300_1115 | sg1115_rpe | A | 0.097 | 30 | AAATCAACAGATAATAATGAgttttAGAGCTAGAAATAGCAAGTTAAAATAAGGC |
| 89 | SAUSA300_1122 | sg1122_plsX | A | 0.279 | 30 | GATATCGTATTAGAAGCCGTgttttAGAGCTAGAAATAGCAAGTTAAAATAAGGC |
| 90 | SAUSA300_1125 | sg1125_acpP | A | 0.224 | 30 | TATCAGCGTCTACACCTAAAgttttAGAGCTAGAAATAGCAAGTTAAAATAAGGC |
| 91 | SAUSA300_1128 | sg1128_ftsY | A | 0.189 | 30 | TTGACCTTGTTCTTCTGTTAgttttAGAGCTAGAAATAGCAAGTTAAAATAAGGC |
| 92 | SAUSA300_1133 | sg1133_trmD | B | 0.014 | 30 | TTTACAACATCAGCAATATAgttttAGAGCTAGAAATAGCAAGTTAAAATAAGGC |
| 93 | SAUSA300_1136 | sg1136_rbgA | A | 0.309 | 30 | TTGGCTTTCGCCATATGTCCgttttAGAGCTAGAAATAGCAAGTTAAAATAAGGC |
| 94 | SAUSA300_1143 | sg1143_topA | A | 0.324 | 30 | TCAATGGTTTTTGCTTTTGCgttttAGAGCTAGAAATAGCAAGTTAAAATAAGGC |
| 95 | SAUSA300_1150 | sg1150_tsf | A | 0.066 | 30 | AATACCTTTTTCACGTgttttAGAGCTAGAAATAGCAAGTTAAAATAAGGC |
| 96 | SAUSA300_1151 | sg1151_pyrH | A | 0.280 | 30 | GCAACACTTTTAATAATTACgttttAGAGCTAGAAATAGCAAGTTAAAATAAGGC |
| 97 | SAUSA300_1152 | sg1152_frr | B | 0.528 | 30 | GCTAATTGTTGTACAGGTGTgttttAGAGCTAGAAATAGCAAGTTAAAATAAGGC |
| 98 | SAUSA300_1153 | sg1153_uppS | B | 0.334 | 30 | TCGTAATGACCTTTAATTCTgttttAGAGCTAGAAATAGCAAGTTAAAATAAGGC |
| 99 | SAUSA300_1156 | sg1156_proS | A | 0.221 | 30 | CATCGTTGGTATAAAAACTTgttttAGAGCTAGAAATAGCAAGTTAAAATAAGGC |
| 100 | SAUSA300_1157 | sg1157_polC | B | 0.415 | 30 | AAAATTAAATGCTTTATATTgttttAGAGCTAGAAATAGCAAGTTAAAATAAGGC |
| 101 | SAUSA300_1159 | sg1159_nusA | A | 0.122 | 30 | GCATCAATTAATACTGCTCTgttttAGAGCTAGAAATAGCAAGTTAAAATAAGGC |

| LCML Number | Locus Tag | Name | Best Clone | R Value | Temp | sgRNA Sequence |
|---|---|---|---|---|---|---|
| 102 | SAUSA300_1168 | sg1168_rnjB | A | 0.312 | 30 | TGAATAGTGAGTTTATATATgttttAGAGCTAGAAATAGCAAGTTAAAATAAGGC |
| 103 | SAUSA300_1176 | sg1176_pgsA | B | 0.129 | 30 | AACCATCAACAAAATCGCTAgttttAGAGCTAGAAATAGCAAGTTAAAATAAGGC |
| 104 | SAUSA300_1200 | sg1200_glnR | A | 0.490 | 30 | TGATGCAATCAGACGAAATAgttttAGAGCTAGAAATAGCAAGTTAAAATAAGGC |
| 105 | SAUSA300_1237 | sg1237_lexA | A | 0.296 | 30 | CCAATTTCGCGAACACTAGGgttttAGAGCTAGAAATAGCAAGTTAAAATAAGGC |
| 106 | SAUSA300_1239 | sg1239_tkt | A | 0.279 | 30 | TTGAAGTAATCTTTAGATTGgttttAGAGCTAGAAATAGCAAGTTAAAATAAGGC |
| 107 | SAUSA300_1249 | sg1249_plsY | B | 0.334 | 30 | TATCATATTCATTTAAATTAgttttAGAGCTAGAAATAGCAAGTTAAAATAAGGC |
| 108 | SAUSA300_1250 | sg1250_parE | A | 0.987 | 30 | GTTGATCCAATATACATACCgttttAGAGCTAGAAATAGCAAGTTAAAATAAGGC |
| 109 | SAUSA300_1257 | sg1257_msrR | B | 0.161 | 30 | TTTCTTAATAAATCGTACTAgttttAGAGCTAGAAATAGCAAGTTAAAATAAGGC |
| 110 | SAUSA300_1269 | sg1269_femA | A | 0.297 | 30 | TTACAGATAGCATGCCATACgttttAGAGCTAGAAATAGCAAGTTAAAATAAGGC |
| 111 | SAUSA300_1270 | sg1270_femB | A | 0.121 | 30 | TTGTACAAAGTTGTCAAATTgttttAGAGCTAGAAATAGCAAGTTAAAATAAGGC |
| 112 | SAUSA300_1311 | sg1311_murG | A | 0.300 | 30 | TGTCCAACTGTTCCCCCTCCgttttAGAGCTAGAAATAGCAAGTTAAAATAAGGC |
| 113 | SAUSA300_1319 | sg1319_folA | A | 0.164 | 30 | CAAGTCATGTGCAACTAGAAgttttAGAGCTAGAAATAGCAAGTTAAAATAAGGC |
| 114 | SAUSA300_1320 | sg1320_thyA | A | 0.077 | 30 | ACTTTCTTTGTCGTTAATAGgttttAGAGCTAGAAATAGCAAGTTAAAATAAGGC |
| 115 | SAUSA300_1340 | sg1340_recU | B | 0.596 | 30 | AATACATTTACCGTTAATGTgttttAGAGCTAGAAATAGCAAGTTAAAATAAGGC |
| 116 | SAUSA300_1341 | sg1341_pbp2 | B | 0.119 | 30 | AATTTAGCTTCGGTAAAAGCgttttAGAGCTAGAAATAGCAAGTTAAAATAAGGC |
| 117 | SAUSA300_1344 | sg1344_dnaD | B | 0.173 | 30 | GATTTTAACAATGAACTTTAgttttAGAGCTAGAAATAGCAAGTTAAAATAAGGC |
| 118 | SAUSA300_1347 | sg1347_birA | B | 0.462 | 30 | ATACCTTGATACCAAATATCgttttAGAGCTAGAAATAGCAAGTTAAAATAAGGC |
| 119 | SAUSA300_1348 | sg1348_papS | B | 0.574 | 30 | CGAACAAATGTAACACCACTgttttAGAGCTAGAAATAGCAAGTTAAAATAAGGC |
| 120 | SAUSA300_1351 | sg1351 | A | 0.767 | 30 | CTTTGTGTTGTGCCCATAAgttttAGAGCTAGAAATAGCAAGTTAAAATAAGGC |
| 121 | SAUSA300_1360 | sg1360_ubiE | B | 0.500 | 30 | ATACCAGTAACTTCACCTGTgttttAGAGCTAGAAATAGCAAGTTAAAATAAGGC |
| 122 | SAUSA300_1362 | sg1362_hup | A | 0.042 | 30 | CATTAGACATTCACCTCCTGgttttAGAGCTAGAAATAGCAAGTTAAAATAAGGC |
| 123 | SAUSA300_1364 | sg1364_engA | A | 0.476 | 30 | ATTGTAGATTTACCTACATTgttttAGAGCTAGAAATAGCAAGTTAAAATAAGGC |
| 124 | SAUSA300_1367 | sg1367_cmk | A | 0.346 | 30 | ACCATCTAATGCAATATTAAgttttAGAGCTAGAAATAGCAAGTTAAAATAAGGC |
| 125 | SAUSA300_1373 | sg1373_fer | A | 0.613 | 30 | TCGTCGTAATCATATATATCgttttAGAGCTAGAAATAGCAAGTTAAAATAAGGC |
| 126 | SAUSA300_1453 | sg1453_rnz | A | 0.877 | 37 | TGTGTATTTCTCTCTTTTGTgttttAGAGCTAGAAATAGCAAGTTAAAATAAGGC |
| 127 | SAUSA300_1454 | sg1454_zwf | A | 0.259 | 30 | CCAAAGATTGTGATTAAACAgttttAGAGCTAGAAATAGCAAGTTAAAATAAGGC |

| LCML Number | Locus Tag | Name | Best Clone | R Value | Temp | sgRNA Sequence |
|---|---|---|---|---|---|---|
| 128 | SAUSA300_1459 | sg1459_gnd | A | 0.024 | 30 | AGCTAGGTTTTTACCCATAAgttttAGAGCTAGAAATAGCAAGTTAAAATAAGGC |
| 129 | SAUSA300_1464 | sg1464_bmfBB | B | 0.332 | 30 | ACAGGAGAAAATGGCATAGAgttttAGAGCTAGAAATAGCAAGTTAAAATAAGGC |
| 130 | SAUSA300_1466 | sg1466_bmfBAA | A | 0.026 | 30 | CTTTTAGGTCTTCTTCGCTAgttttAGAGCTAGAAATAGCAAGTTAAAATAAGGC |
| 131 | SAUSA300_1475 | sg1475_accC | B | 0.284 | 30 | TGCACATATGCTTCTTCGTCgttttAGAGCTAGAAATAGCAAGTTAAAATAAGGC |
| 132 | SAUSA300_1490 | sg1490_efp | A | 0.594 | 30 | CTTTACCAGGCTTTACATGTgttttAGAGCTAGAAATAGCAAGTTAAAATAAGGC |
| 133 | SAUSA300_1492 | sg1492 | B | 0.466 | 37 | AAATGCACAAACGTTTCACTgttttAGAGCTAGAAATAGCAAGTTAAAATAAGGC |
| 134 | SAUSA300_1520 | sg1520_trmK | A | 0.523 | 30 | ATTCTCCTTTATGAAAAAAGgttttAGAGCTAGAAATAGCAAGTTAAAATAAGGC |
| 135 | SAUSA300_1521 | sg1521_rpoD | A | 0.113 | 30 | CTGTGTTATCAGACATGAAAgttttAGAGCTAGAAATAGCAAGTTAAAATAAGGC |
| 136 | SAUSA300_1522 | sg1522_dnaG | A | 0.696 | 30 | ATTTATTTTGTTCAATATAAgttttAGAGCTAGAAATAGCAAGTTAAAATAAGGC |
| 137 | SAUSA300_1525 | sg1525_glyS | A | 0.143 | 37 | TCATACATGAAAACGCCCCAgttttAGAGCTAGAAATAGCAAGTTAAAATAAGGC |
| 138 | SAUSA300_1530 | sg1530_ybeY | A | 0.202 | 30 | TCAATGATCTTACTTACCAAgttttAGAGCTAGAAATAGCAAGTTAAAATAAGGC |
| 139 | SAUSA300_1540 | sg1540_dnaK | A | 0.726 | 30 | TCAGGGTTTTGAATTACTTTgttttAGAGCTAGAAATAGCAAGTTAAAATAAGGC |
| 140 | SAUSA300_1541 | sg1541_grpE | A | 0.805 | 30 | TCATTAATTTTTTGATCTTTgttttAGAGCTAGAAATAGCAAGTTAAAATAAGGC |
| 141 | SAUSA300_1546 | sg1546_holA | A | 0.815 | 30 | CTTTGTTTTTCAACCAATTCgttttAGAGCTAGAAATAGCAAGTTAAAATAAGGC |
| 142 | SAUSA300_1557 | sg1557 | A | 0.016 | 30 | ATTGATTGAACATATGAATTgttttAGAGCTAGAAATAGCAAGTTAAAATAAGGC |
| 143 | SAUSA300_1567 | sg1567_greA | A | 0.687 | 30 | TCAAAACCTTCTTGAGTCATgttttAGAGCTAGAAATAGCAAGTTAAAATAAGGC |
| 144 | SAUSA300_1575 | sg1575_alaS | A | 0.772 | 30 | ATTGGCACTAATGGTGCAGAgttttAGAGCTAGAAATAGCAAGTTAAAATAAGGC |
| 145 | SAUSA300_1579 | sg1579_iscS | A | 0.135 | 30 | TACTTCAGGTTTTACTGGTGgttttAGAGCTAGAAATAGCAAGTTAAAATAAGGC |
| 146 | SAUSA300_1587 | sg1587_hisS | A | 0.481 | 30 | AAAATATCCTGCGTCCCTCTgttttAGAGCTAGAAATAGCAAGTTAAAATAAGGC |
| 147 | SAUSA300_1589 | sg1589_dtd | A | 0.460 | 37 | GTATTTCACCATTCATTTGTgttttAGAGCTAGAAATAGCAAGTTAAAATAAGGC |
| 148 | SAUSA300_1593 | sg1593_secF | A | 0.763 | 30 | TTTATAAGTTGCAGCCATTCgttttAGAGCTAGAAATAGCAAGTTAAAATAAGGC |
| 149 | SAUSA300_1600 | sg1600_obgE | A | 0.045 | 30 | AATACCATTACCACCATCACgttttAGAGCTAGAAATAGCAAGTTAAAATAAGGC |
| 150 | SAUSA300_1602 | sg1602 | A | 0.205 | 30 | CATATTCACCATGGTCAGCAgttttAGAGCTAGAAATAGCAAGTTAAAATAAGGC |
| 151 | SAUSA300_1611 | sg1611_valS | B | 0.363 | 30 | TACCAGTTACATTTGGTGGCgttttAGAGCTAGAAATAGCAAGTTAAAATAAGGC |
| 152 | SAUSA300_1619 | sg1619_hemA | B | 0.185 | 30 | TACAAAAACTATGAAATATAgttttAGAGCTAGAAATAGCAAGTTAAAATAAGGC |
| 153 | SAUSA300_1620 | sg1620_engB | A | 0.093 | 30 | ATGATTAATTCTATATTATTgttttAGAGCTAGAAATAGCAAGTTAAAATAAGGC |

| LCML Number | Locus Tag | Name | Best Clone | R Value | Temp | sgRNA Sequence |
|---|---|---|---|---|---|---|
| 154 | SAUSA300_1624 | sg1624 | B | 0.395 | 30 | AAATGAGTTGTTTATATGAgttttAGAGCTAGAAATAGCAAGTTAAAATAAGGC |
| 155 | SAUSA300_1627 | sg1627_infC | A | 0.040 | 30 | TTGAGTTTGATCTTTTGCTAgttttAGAGCTAGAAATAGCAAGTTAAAATAAGGC |
| 156 | SAUSA300_1629 | sg1629_thrS | B | 0.703 | 30 | ATTGATCCATCAGTTTCAAGgttttAGAGCTAGAAATAGCAAGTTAAAATAAGGC |
| 157 | SAUSA300_1631 | sg1631_dnaB | A | 0.425 | 30 | TGGTCTTAAGCCGAATTCGAgttttAGAGCTAGAAATAGCAAGTTAAAATAAGGC |
| 158 | SAUSA300_1632 | sg1632_nrdR | B | 0.569 | 30 | CTAACTCCGAAGTCAGAGTTgttttAGAGCTAGAAATAGCAAGTTAAAATAAGGC |
| 159 | SAUSA300_1634 | sg1634_coaE | B | 0.501 | 30 | CTTCCACACACTTTGCATACgttttAGAGCTAGAAATAGCAAGTTAAAATAAGGC |
| 160 | SAUSA300_1645 | sg1645_pfkA | A | 0.311 | 30 | TGTACGAACAACTGCTCTTAgttttAGAGCTAGAAATAGCAAGTTAAAATAAGGC |
| 161 | SAUSA300_1647 | sg1647_accD | B | 0.332 | 30 | TGAACTTGACCAGTTAAAAAgttttAGAGCTAGAAATAGCAAGTTAAAATAAGGC |
| 162 | SAUSA300_1649 | sg1649_dnaE | B | 0.229 | 30 | ACTGAAACTCCTGACGCATTgttttAGAGCTAGAAATAGCAAGTTAAAATAAGGC |
| 163 | SAUSA300_1657 | sg1657_ackA | B | 0.652 | 30 | ACTATTACAGATTATTTTTTgttttAGAGCTAGAAATAGCAAGTTAAAATAAGGC |
| 164 | SAUSA300_1673 | sg1673_plsC | A | 0.054 | 30 | ACGACATATTTACTATCCTTgttttAGAGCTAGAAATAGCAAGTTAAAATAAGGC |
| 165 | SAUSA300_1675 | sg1675_tyrS | A | 0.485 | 37 | TAAACTATCTGCCGTTGGATgttttAGAGCTAGAAATAGCAAGTTAAAATAAGGC |
| 166 | SAUSA300_1686 | sg1686_murC | A | 0.116 | 30 | TTTATGTTATTAGCATCAAAgttttAGAGCTAGAAATAGCAAGTTAAAATAAGGC |
| 167 | SAUSA300_1688 | sg1688 | A | 0.152 | 30 | GCGACATCTCCTACATATTTgttttAGAGCTAGAAATAGCAAGTTAAAATAAGGC |
| 168 | SAUSA300_1704 | sg1704_leuS | A | 0.380 | 37 | ATTTCTTTTCAATTTGATTGgttttAGAGCTAGAAATAGCAAGTTAAAATAAGGC |
| 169 | SAUSA300_1729 | sg1729 | A | 0.624 | 30 | ATAATGAGTCCAATGACTATgttttAGAGCTAGAAATAGCAAGTTAAAATAAGGC |
| 170 | SAUSA300_1730 | sg1730_metK | B | 0.338 | 30 | TGTTGTTGTAGAAATTTCGCgttttAGAGCTAGAAATAGCAAGTTAAAATAAGGC |
| 171 | SAUSA300_1748 | sg1748 | B | 0.492 | 30 | AGTAGAGTCGCCTATCTCTCgttttAGAGCTAGAAATAGCAAGTTAAAATAAGGC |
| 172 | SAUSA300_1749 | sg1749 | B | 0.401 | 30 | GTTACAACCCATATGATTGTgttttAGAGCTAGAAATAGCAAGTTAAAATAAGGC |
| 173 | SAUSA300_1869 | sg1869_map | B | 0.247 | 30 | AAGATGTTACTTTAGTATTTgttttAGAGCTAGAAATAGCAAGTTAAAATAAGGC |
| 174 | SAUSA300_1873 | sg1873_murT | A | 0.241 | 30 | TACGCGCCAATTTCGCTAGAgttttAGAGCTAGAAATAGCAAGTTAAAATAAGGC |
| 175 | SAUSA300_1879 | sg1879_dgkB | A | 0.009 | 30 | ATAGCTCTTTACCTGATGTCgttttAGAGCTAGAAATAGCAAGTTAAAATAAGGC |
| 176 | SAUSA300_1881 | sg1881_gatA | A | 0.119 | 30 | ATATCTTTAACAACATCAGAgttttAGAGCTAGAAATAGCAAGTTAAAATAAGGC |
| 177 | SAUSA300_1882 | sg1882_gatC | A | 0.069 | 30 | GCCATTTCTTCCGTTTCTTCgttttAGAGCTAGAAATAGCAAGTTAAAATAAGGC |
| 178 | SAUSA300_1885 | sg1885_ligA | A | 0.252 | 30 | TATTCACTATCTGGTACAGAgttttAGAGCTAGAAATAGCAAGTTAAAATAAGGC |
| 179 | SAUSA300_1886 | sg1886_pcrA | A | 0.358 | 30 | GCACCTGCCATAATTAACAAgttttAGAGCTAGAAATAGCAAGTTAAAATAAGGC |

| LCML Number | Locus Tag | Name | Best Clone | R Value | Temp | sgRNA Sequence |
|---|---|---|---|---|---|---|
| 180 | SAUSA300_1894 | sg1894_pncB | B | 0.555 | 30 | GTAGACTATAATATAAAGCGgttttAGAGCTAGAAATAGCAAGTTAAAATAAGGC |
| 181 | SAUSA300_1900 | sg1900_ppaC | B | 0.515 | 30 | AATGAAGCCACTCCCTCAGCgttttAGAGCTAGAAATAGCAAGTTAAAATAAGGC |
| 182 | SAUSA300_1911 | sg1911_pmtC | B | 0.496 | 30 | AAAATATATGAATATAAATCgttttAGAGCTAGAAATAGCAAGTTAAAATAAGGC |
| 183 | SAUSA300_1913 | sg1913_pmtA | A | 0.496 | 30 | ATTAACATTACTTAATTCTAgttttAGAGCTAGAAATAGCAAGTTAAAATAAGGC |
| 184 | SAUSA300_1983 | sg1983_groES | A | 0.648 | 30 | ATAATCACACGATTTCCAATgttttAGAGCTAGAAATAGCAAGTTAAAATAAGGC |
| 185 | SAUSA300_1988 | sg1988_hld | B | 0.420 | 30 | AGTATTTATTTCCTACAGTTgttttAGAGCTAGAAATAGCAAGTTAAAATAAGGC |
| 186 | SAUSA300_2002 | sg2002_gcp | A | 0.199 | 30 | TGATGTCTACTTGCCACTTCgttttAGAGCTAGAAATAGCAAGTTAAAATAAGGC |
| 187 | SAUSA300_2005 | sg2005_tsaE | A | 0.195 | 30 | TTTAATGATGTTAAATGTCGgttttAGAGCTAGAAATAGCAAGTTAAAATAAGGC |
| 188 | SAUSA300_2028 | sg2028_acpS | B | 0.280 | 30 | CTTGACCACCCGCTGTATAAgttttAGAGCTAGAAATAGCAAGTTAAAATAAGGC |
| 189 | SAUSA300_2039 | sg2039_ddl | A | 0.066 | 30 | CTTTCTCCAATCACCATCATgttttAGAGCTAGAAATAGCAAGTTAAAATAAGGC |
| 190 | SAUSA300_2046 | sg2046_oxaA | A | 0.102 | 30 | ACCATAATACCTAAAAATAAgttttAGAGCTAGAAATAGCAAGTTAAAATAAGGC |
| 191 | SAUSA300_2055 | sg2055_murA | A | 0.509 | 30 | AATAAAGATGCTGTCAATATgttttAGAGCTAGAAATAGCAAGTTAAAATAAGGC |
| 192 | SAUSA300_2062 | sg2062_atpF | B | 0.278 | 30 | CTGAATTTATGCGATAGGCAgttttAGAGCTAGAAATAGCAAGTTAAAATAAGGC |
| 193 | SAUSA300_2072 | sg2072_prfA | A | 0.221 | 30 | CTGAATCATTTACAACATCTgttttAGAGCTAGAAATAGCAAGTTAAAATAAGGC |
| 194 | SAUSA300_2079 | sg2079_fbaA | A | 0.437 | 30 | ATTTCTTTCATTGAAACTAAgttttAGAGCTAGAAATAGCAAGTTAAAATAAGGC |
| 195 | SAUSA300_2081 | sg2081_pyrG | A | 0.229 | 30 | CCTGGGTCAACATTTAAGTAgttttAGAGCTAGAAATAGCAAGTTAAAATAAGGC |
| 196 | SAUSA300_2084 | sg2084_coaA | A | 0.523 | 30 | ATTCAGTTTTAAAAGTACGTgttttAGAGCTAGAAATAGCAAGTTAAAATAAGGC |
| 197 | SAUSA300_2104 | sg2104_glmS | A | 0.379 | 30 | ACCTTTTAATAATAATTCTTgttttAGAGCTAGAAATAGCAAGTTAAAATAAGGC |
| 198 | SAUSA300_2113 | sg2113_dacA | A | 0.321 | 37 | ACTGAGGTTTTGAAAAAAGTgttttAGAGCTAGAAATAGCAAGTTAAAATAAGGC |
| 199 | SAUSA300_2182 | sg2182_infA | A | 0.438 | 30 | AACGCAATGTTTAAAGTAGAgttttAGAGCTAGAAATAGCAAGTTAAAATAAGGC |
| 200 | SAUSA300_2184 | sg2184_secY | A | 0.135 | 37 | TGTTCTAAAGAAGTTCACAAgttttAGAGCTAGAAATAGCAAGTTAAAATAAGGC |
| 201 | SAUSA300_2214 | sg2214_femX | A | 0.244 | 30 | TAATAAATCTCCATTTGGGTgttttAGAGCTAGAAATAGCAAGTTAAAATAAGGC |
| 202 | SAUSA300_2238 | sg2238_ureA | B | 0.536 | 30 | ATCTAATCGAAAACAAATAGgttttAGAGCTAGAAATAGCAAGTTAAAATAAGGC |
| 203 | SAUSA300_2283 | sg2283_rpiA | B | 0.380 | 30 | ATTTTAAATAATACTCGTTAgttttAGAGCTAGAAATAGCAAGTTAAAATAAGGC |
| 204 | SAUSA300_2292 | sg2292_fni | A | 0.357 | 30 | TGAATGCATTGCGTCAGATTgttttAGAGCTAGAAATAGCAAGTTAAAATAAGGC |
| 205 | SAUSA300_2483 | sg2483_mvaA | A | 0.282 | 30 | ATTCTTATCTAAATTTTGCAgttttAGAGCTAGAAATAGCAAGTTAAAATAAGGC |

| LCML Number | Locus Tag | Name | Best Clone | R Value | Temp | sgRNA Sequence |
|---|---|---|---|---|---|---|
| 206 | SAUSA300_2646 | sg2646_trmE | A | 0.283 | 30 | CCAATTGCCCCTTCACCCATgttttAGAGCTAGAAATAGCAAGTTAAAATAAGGC |
| 207 | SAUSA300_2484 | sg2484_mvaS | A | 0.277 | 30 | GCCATGTCTACATAGTACTTgttttAGAGCTAGAAATAGCAAGTTAAAATAAGGC |
| 208 | SAUSA300_0005 | sg0005_gyrB | B | 0.352 | 30 | GTCGATCCTATATACATACCgttttAGAGCTAGAAATAGCAAGTTAAAATAAGGC |
| 209 | SAUSA300_0527 | sg0527_rpoB | B | 0.312 | 30 | ATAAAAAGACAAAAAGAAAAgttttAGAGCTAGAAATAGCAAGTTAAAATAAGGC |
| 210 | SAUSA300_0425 | sg0425_mspA | A | 0.497 | 30 | ATAGTAGATTCTGTACATAAgttttAGAGCTAGAAATAGCAAGTTAAAATAAGGC |
| 211 | SAUSA300_0639 | sg0639 | B | 0.697 | 30 | ATAAAACTGCCTTCAACAATgttttAGAGCTAGAAATAGCAAGTTAAAATAAGGC |
| 212 | SAUSA300_1359 | sg1359 | B | 0.483 | 30 | TACTCAGAATAACAAATGCTgttttAGAGCTAGAAATAGCAAGTTAAAATAAGGC |
| 213 | SAUSA300_1590 | sg1590_relA | B | 0.445 | 37 | TGTATGGTAATCCGTTTTTTgttttAGAGCTAGAAATAGCAAGTTAAAATAAGGC |
| 214 | SAUSA300_1791 | sg1791_cbf1 | B | 0.367 | 37 | TTAACATGTACAATTTCTTCgttttAGAGCTAGAAATAGCAAGTTAAAATAAGGC |
| 215 | SAUSA300_2183 | sg2183_adk | A | 0.310 | 30 | TGAGTTCCTTTACCTGCGCCgttttAGAGCTAGAAATAGCAAGTTAAAATAAGGC |
| 216 | SAUSA300_2185 | sg2185_rplO | A | 0.372 | 30 | TAACTCATGTAATTTCATTTgttttAGAGCTAGAAATAGCAAGTTAAAATAAGGC |
| 217 | SAUSA300_2647 | sg2647_rnpA | A | 0.385 | 30 | AATCTGCATTCTTTTTAATTgttttAGAGCTAGAAATAGCAAGTTAAAATAAGGC |
| 218 | SAUSA300_2648 | sg2648_rpmH | A | 0.314 | 30 | TTACTATGTTTACGTTTATTgttttAGAGCTAGAAATAGCAAGTTAAAATAAGGC |
| 219 | SAUSA300_0015 | sg0015_rplI | B | 0.602 | 30 | TTTAAGTTGTGTTGCCGCATgttttAGAGCTAGAAATAGCAAGTTAAAATAAGGC |
| 220 | SAUSA300_0366 | sg0366_rpsF | A | 0.347 | 30 | TTTATATTTGCACCTCCTTGgttttAGAGCTAGAAATAGCAAGTTAAAATAAGGC |
| 221 | SAUSA300_0479 | sg0479_rplY | A | 0.506 | 30 | AACGTGTTTGTTTACCTTGAgttttAGAGCTAGAAATAGCAAGTTAAAATAAGGC |
| 222 | SAUSA300_0522 | sg0522_rplK | B | 0.321 | 30 | GCTGGACCAACTGGTGGTGCgttttAGAGCTAGAAATAGCAAGTTAAAATAAGGC |
| 223 | SAUSA300_0524 | sg0524_rplJ | B | 0.314 | 30 | CTTCAGCTACTGTTAATCCAgttttAGAGCTAGAAATAGCAAGTTAAAATAAGGC |
| 224 | SAUSA300_0530 | sg0530_rpsL | A | 0.191 | 30 | CGTACTAATTGGTTAATAGTgttttAGAGCTAGAAATAGCAAGTTAAAATAAGGC |
| 225 | SAUSA300_0990 | sg0990_rpoY | A | 0.358 | 37 | TAACCCTAGTAAAATCGTATgttttAGAGCTAGAAATAGCAAGTTAAAATAAGGC |
| 226 | SAUSA300_1027 | sg1027_rpmF | A | 0.404 | 30 | GTTTTAGAAGTTCTTCTTTTgttttAGAGCTAGAAATAGCAAGTTAAAATAAGGC |
| 227 | SAUSA300_1117 | sg1117_rpmB | A | 0.497 | 30 | CAGTTTACTCAAAATATAATgttttAGAGCTAGAAATAGCAAGTTAAAATAAGGC |
| 228 | SAUSA300_1131 | sg1131_rpsP | A | 0.260 | 30 | GCTACTACGATACGATAGAAgttttAGAGCTAGAAATAGCAAGTTAAAATAAGGC |
| 229 | SAUSA300_1134 | sg1134_rplS | A | 0.443 | 30 | CCTCGACAAATATATAGCAGgttttAGAGCTAGAAATAGCAAGTTAAAATAAGGC |
| 230 | SAUSA300_1149 | sg1149_rpsB | A | 0.286 | 30 | CCATTATAAATTCCTCCTATgttttAGAGCTAGAAATAGCAAGTTAAAATAAGGC |
| 231 | SAUSA300_1166 | sg1166_rpsO | B | 0.425 | 30 | AGTACAGCGATTTGTACTTCgttttAGAGCTAGAAATAGCAAGTTAAAATAAGGC |

| LCML Number | Locus Tag | Name | Best Clone | R Value | Temp | sgRNA Sequence |
|---|---|---|---|---|---|---|
| 232 | SAUSA300_1511 | sg1511_rpmG | A | 0.368 | 30 | TAAAGTTACGTTTACGCGCAgttttAGAGCTAGAAATAGCAAGTTAAAATAAGGC |
| 233 | SAUSA300_1535 | sg1535_rpsU | A | 0.321 | 30 | TCCCTCCCTCCAAATATCAAgttttAGAGCTAGAAATAGCAAGTTAAAATAAGGC |
| 234 | SAUSA300_1545 | sg1545_rpsT | B | 0.171 | 37 | CTTTTAGGAGGTGACAGAAAgttttAGAGCTAGAAATAGCAAGTTAAAATAAGGC |
| 235 | SAUSA300_1601 | sg1601_rpmA | B | 0.236 | 37 | TTGTCGTCATAATTGATATCgttttAGAGCTAGAAATAGCAAGTTAAAATAAGGC |
| 236 | SAUSA300_1625 | sg1625_rplT | B | 0.326 | 37 | TAACACGTTTAGCTGCTCCGgttttAGAGCTAGAAATAGCAAGTTAAAATAAGGC |
| 237 | SAUSA300_1666 | sg1666_rpsD | A | 0.457 | 37 | CCATGTTGTCCTGGTGCGTAgttttAGAGCTAGAAATAGCAAGTTAAAATAAGGC |
| 238 | SAUSA300_2074 | sg2074_rpmE2 | A | 0.276 | 30 | CTCCTTTGCCCTGAACCATCgttttAGAGCTAGAAATAGCAAGTTAAAATAAGGC |
| 239 | SAUSA300_2171 | sg2171_rpsI | B | 0.577 | 30 | ACTGTGATGTTACCTTCACCgttttAGAGCTAGAAATAGCAAGTTAAAATAAGGC |
| 240 | SAUSA300_2177 | sg2177_rplQ | A | 0.839 | 30 | GAAAAGAAGATTGATAAAGGgttttAGAGCTAGAAATAGCAAGTTAAAATAAGGC |
| 241 | SAUSA300_0386 | sg0386_xpt | A | 0.899 | 30 | CCAAACTAAATAATAGTTTCgttttAGAGCTAGAAATAGCAAGTTAAAATAAGGC |
| 242 | SAUSA300_0529 | sg0529_rplGB | A | 0.355 | 30 | ATTTACATAAAAATAACAAGgttttAGAGCTAGAAATAGCAAGTTAAAATAAGGC |
| 243 | SAUSA300_0775 | sg0755_gapR | A | 0.417 | 30 | CAACTTTCAAAGTAGAAAAAgttttAGAGCTAGAAATAGCAAGTTAAAATAAGGC |
| 244 | SAUSA300_0906 | sg0906 | A | 0.438 | 30 | CTTACGAACACTATTTTTAAgttttAGAGCTAGAAATAGCAAGTTAAAATAAGGC |
| 245 | SAUSA300_1072 | sg1072_mraZ | B | 0.208 | 37 | ATTTAAGTCATAACGAAACTgttttAGAGCTAGAAATAGCAAGTTAAAATAAGGC |
| 246 | SAUSA300_1108 | sg1108_def | A | 0.479 | 30 | GTAATCCTAAATTTATTGTAgttttAGAGCTAGAAATAGCAAGTTAAAATAAGGC |
| 247 | SAUSA300_1121 | sg1121_fapR | A | 0.415 | 30 | AGTTCATGGTCTGTGATGAAgttttAGAGCTAGAAATAGCAAGTTAAAATAAGGC |
| 248 | SAUSA300_1155 | sg1155_rasP | A | 0.307 | 30 | CCCATACCGATCGCAAATTCgttttAGAGCTAGAAATAGCAAGTTAAAATAAGGC |
| 249 | SAUSA300_1158 | sg1158_rimP | B | 0.279 | 30 | TAAGACATTGAAAAGAAATAgttttAGAGCTAGAAATAGCAAGTTAAAATAAGGC |
| 250 | SAUSA300_1558 | sg1558_mtnN | A | 0.406 | 30 | TATTGTTACTTCTTCTTCCAgttttAGAGCTAGAAATAGCAAGTTAAAATAAGGC |
| 251 | SAUSA300_1635 | sg1635_mutM | A | 0.493 | 30 | CTTTTTACATGTTCTACTTCgttttAGAGCTAGAAATAGCAAGTTAAAATAAGGC |
| 252 | SAUSA300_1690 | sg1690 | A | 0.444 | 30 | GGTTCTATCACTCTACAATCgttttAGAGCTAGAAATAGCAAGTTAAAATAAGGC |
| 253 | SAUSA300_1914 | sg1914_pmtR | A | 0.357 | 30 | TGCTTAATCTGTTCATAAATgttttAGAGCTAGAAATAGCAAGTTAAAATAAGGC |
| 254 | SAUSA300_2029 | sg2029 | B | 0.637 | 30 | CGTATTTTAAGTTAATCGATgttttAGAGCTAGAAATAGCAAGTTAAAATAAGGC |
| 255 | SAUSA300_2205 | sg2205_rpsJ | A | 0.298 | 30 | CTGCTGATTGATCAATTACGgttttAGAGCTAGAAATAGCAAGTTAAAATAAGGC |
| 256 | SAUSA300_0014 | sg0014_gdpP | A | 0.532 | 30 | TATTAGTAAAGCTTTCTTAGgttttAGAGCTAGAAATAGCAAGTTAAAATAAGGC |
| 257 | SAUSA300_1603 | sg1603_rplU | A | 0.273 | 30 | TAATAAGTCACGCCATACATgttttAGAGCTAGAAATAGCAAGTTAAAATAAGGC |

| LCML Number | Locus Tag | Name | Best Clone | R Value | Temp | sgRNA Sequence |
|---|---|---|---|---|---|---|
| 258 | SAUSA300_2172 | sg2172_rplM | A | 0.453 | 30 | ATGATAAACGACCTAATGTTgttttAGAGCTAGAAATAGCAAGTTAAAATAAGGC |
| 259 | SAUSA300_1476 | sg1476_accB | A | 0.310 | 30 | TTCAAGCATGTTCATATTGCgttttAGAGCTAGAAATAGCAAGTTAAAATAAGGC |
| 260 | SAUSA300_1700 | sg1700_murJ | A | 0.027 | 30 | CTTGGTAATTAATATACTAAgttttAGAGCTAGAAATAGCAAGTTAAAATAAGGC |
| 261 | SAUSA300_1080 | sg1080_ftsZ | A | 0.004 | 30 | AAATTTCCTCCTAGTTTTAgttttAGAGCTAGAAATAGCAAGTTAAAATAAGGC |
